# Mutations outside the MR1 antigen binding groove differentially inhibit presentation of exogenous antigens

**DOI:** 10.1101/2025.05.14.654109

**Authors:** Corinna A. Kulicke, Chance Lemon, Jason R. Krawic, Luisa Maria Nieto Ramirez, Se-Jin Kim, Gitanjali Narayanan, Fikadu G. Tafesse, William H. Hildebrand, Karen M. Dobos, David M. Lewinsohn

## Abstract

The antigen presenting molecule MHC class I-related protein 1 (MR1) binds small molecule metabolites derived from microbial riboflavin biosynthetic pathways and presents them at the cell surface for surveillance by MR1-restricted mucosal-associated invariant T cells (MAIT cells). MR1 ligands can originate in the extracellular space or in endosomal compartments that contain microbial pathogens. Distinct, complementary antigen processing and presentation pathways enable MR1 to survey diverse intracellular locations and present both exogenous and intracellular antigens. Here, we generated a panel of BEAS-2B MR1 KO cells reconstituted with MR1 proteins mutated at amino acids 9 – 16. The mutated MR1 molecules differentially translocated to the cell surface in response to 6-formylpterin and differed in their ability to present mycobacterial antigens to MAIT cell clones. While they barely presented Mycobacterium smegmatis supernatant and other exogenous MAIT cell antigens, their ability to present antigens derived from mycobacterial infection and a 5-A-RU prodrug requiring endosomal processing remained largely intact. Protein co-immunoprecipitation and mass spectrometry-based proteomic analysis showed that mutated MR1 differentially associated with calnexin and β_2_-microglobulin (B2M). Knock-down of B2M in cells over-expressing MR1 phenocopied the loss of exogenous antigen presentation but did not impact presentation of intracellular antigens. Thus, the MR1-mediated presentation of exogenous antigen appears to be limited by binding to B2M whereas the lower sensitivity to B2M deficiency implies that MAIT cell activation via the endosomal antigen presentation pathway may be limited by the availability of MR1 itself.

## Introduction

Major histocompatibility complex (MHC) class I-related protein 1 (MR1) presents small molecule antigens to MR1-restricted T (MR1T) cells, including mucosal associated invariant T (MAIT) cells, which express the canonical TRAV1-2 T cell receptor (TCR) alpha chain [1]. The MR1-MR1T cell axis has recently attracted much interest in the context of bacterial and viral infections as well as sterile inflammatory and non-inflammatory conditions such as auto-immunity and cancer [2–4]. MAIT cells are a particularly promising therapeutic target due to their abundance, especially in tissues, their restriction by a non-polymorphic antigen presenting molecule, and their capacity to rapidly respond to immune insults [2, 3, 5]. While the broader group of MR1T cells shares the restricting molecule MR1, both its TCR repertoire and its effector phenotypes are more diverse than those of TRAV1-2^+^ MAIT cells [1, 6]. Unlike peptide-specific human leukocyte antigen class Ia (HLA-Ia)-restricted CD4^+^ and CD8^+^ T cells, MAIT cells respond to small molecules. The first MAIT cell antigens identified were a group of ribityllumazine compounds [7] and 5-(2-oxopropylideneamino)-6-D-ribitylaminouracil (5-OP-RU) [8]. These molecules are derivatives of metabolites of bacterial and fungal riboflavin biosynthesis which can either cyclize or react with byproducts of cellular metabolism to form the MR1 ligands [7, 8]. Since humans do not express the enzymes catalyzing riboflavin synthesis, the MR1-MAIT cell axis is a sensitive detection mechanism of microbial products [9].

While 5-OP-RU remains the most potent MAIT cell-activating MR1 ligand discovered to date, it has become increasingly clear that antigens presented by MR1 are more diverse than initially appreciated. Studies by various research groups have identified diverse small molecules derived from bacterial metabolism [10, 11], the host itself [12], and environmental sources [13–15]. Interestingly, the composition of the MR1 ligand repertoire generated by a particular microbe appears to be unique and has been linked to the expression levels of enzymes in the riboflavin synthesis pathway [16, 17]. How these diverse microbial antigens are loaded onto and presented by MR1 is only partially understood. Research has traditionally focused on the presentation of purified, exogenous antigens added directly to the medium in tissue culture systems. Presentation of ligands supplied to the extracellular milieu in this fashion is primarily mediated by an HLA-Ia-like, endoplasmic reticulum (ER)-based antigen presentation pathway which has been widely studied [18–21]. However, mounting evidence supports the hypothesis that MR1 ligands derived from bacteria in the context of intracellular infection use a different cellular mechanism, possibly more akin to an MHC class II or CD1 pathway [20–22]. Most recently, Karamooz et al showed that endosomal calcium signaling plays a unique and specific role in the MR1-mediated presentation of intracellular Mycobacterium tuberculosis (Mtb) but not Mycobacterium smegmatis (M. smeg) supernatant or antigenic peptide from Mtb protein CFP10 [22]. Thus, the presence of multiple, non-exclusive, complementary pathways for MR1 antigen presentation has been proposed [23, 24].

In this study, we characterize a panel of MR1 mutants which are differentially impaired in their ability to present soluble, exogenous ligands while retaining the ability to present antigen in the context of intracellular Mtb infection and endosomal processing. We find that this antigen presentation pathway is independent of the ability to translocate to the cell surface in response to the MR1-stabilising ligand 6-formylpterin (6-FP) but linked to the extent of binding to β_2_-microglobulin (B2M). These mutants present a unique opportunity to study the presentation of MR1T cell antigens from intracellular infection since the pathways are differentially affected by the mutations.

## Results

### A panel of MR1 mutants differentially translocates to the cell surface in response to stabilizing ligand

Our lab has previously used CRISPR/Cas9 technology to generate a BEAS-2B cell line deficient for MR1 [25, 26]. Subsequent transduction of these cells with a lentiviral vector encoding GFP-tagged MR1A under the control of a tetracycline (tet)-inducible promoter reconstituted MR1-mediated antigen presentation [25], but resulted in a heterogeneous population of cells with varying levels of MR1 surface expression (Figure 1A, [13]). As the reconstituted cell line was sorted based on GFP expression, the majority of the cells expressed GFP upon induction with doxycycline (dox). However, we were surprised to find that only a subset of the GFP^+^ cells expressed MR1 at the cell surface. We postulated that this was due to constitutive expression of the Cas9 protein and the single guide RNA (sgRNA) targeting MR1 in the BEAS-2B MR1 knock out (KO) parent cell line and, thus, continuous editing of the MR1 coding sequence, including the MR1-GFP sequence encoded by the lentiviral insertions. Here, we sub-cloned the polyclonal BEAS-2B MR1 KO tet MR1-GFP parent cell line and characterized a panel of MR1 mutants which differed in their ability to present mycobacterial MAIT cell antigens (Figures 1 and 2). As expected, all four clonal cell lines expressed GFP when induced with dox overnight (Figure 1B, bottom). However, MR1 surface expression as detected by the anti-MR1 antibody clone 26.5 [27] differed dramatically between the mutant cell lines (Figure 1B). D4 expressed MR1 at the cell surface at baseline, characteristic for most overexpression systems, and further increased MR1 surface expression after incubation with the stabilizing ligand 6-formylpterin (6-FP). The MR1 in D6, on the other hand, was undetectable at baseline, but consistently translocated to the cell surface in the presence of 6-FP. D8 and D16 had no 26.5-reactive MR1 at their cell surface at baseline. While D8 expressed a very small amount of MR1 at the cell surface in response to 6-FP, surface levels of MR1 in D16 remained undetectable irrespective of 6-FP incubation. As expected, HLA-Ia surface expression as measured with the W6/32 antibody was unaffected by 6-FP incubation and dox induction. Interestingly, HLA-Ia surface expression tended to be higher in cell lines with lower MR1 surface expression, a phenomenon described previously [28].

**Figure 1:**
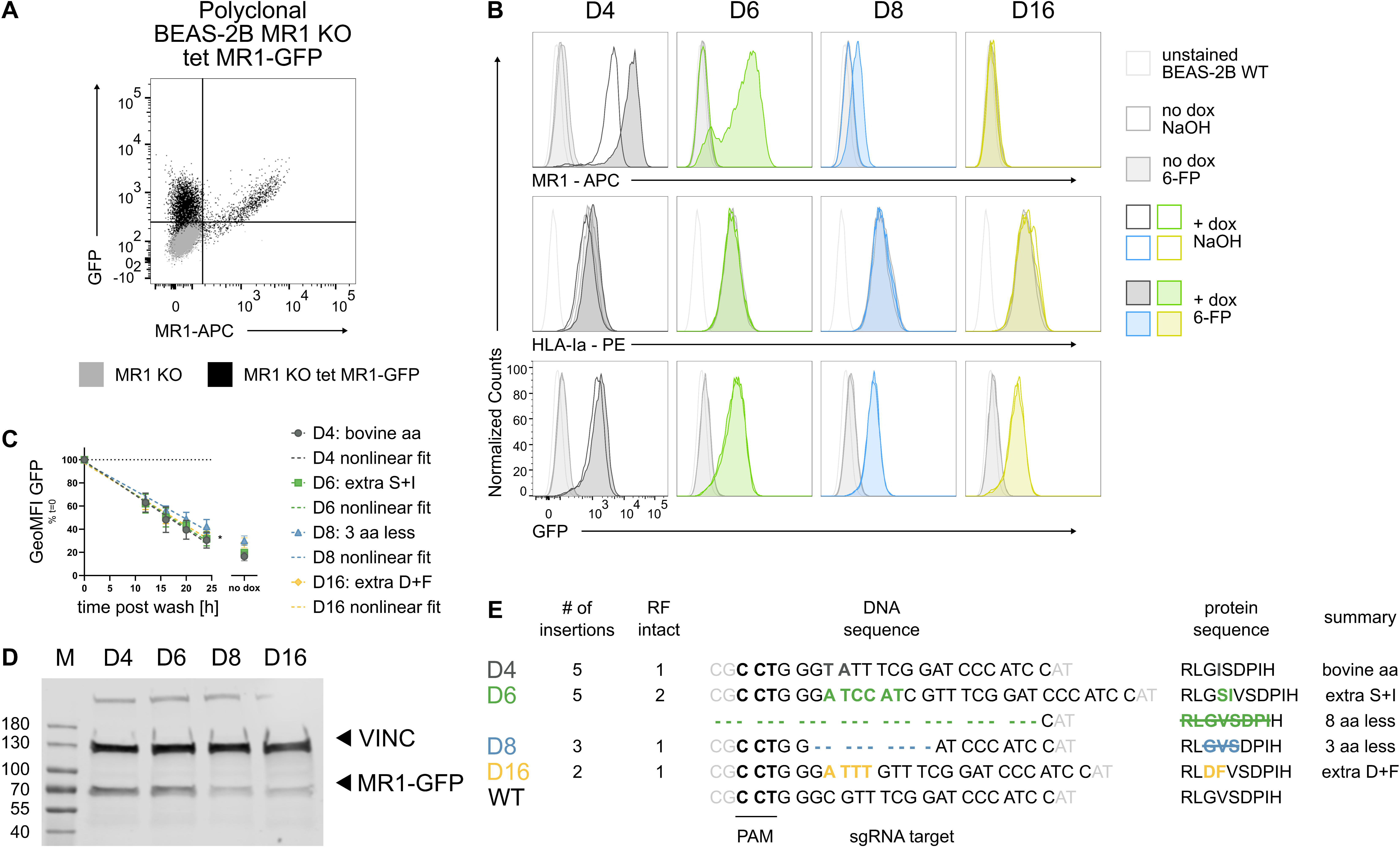
A panel of MR1 mutants differentially translocates to the cell surface in response to stabilizing ligand. A. Polyclonal BEAS-2B MR1 KO tet MR1-GFP cells were induced to express MR1-GFP with 2 μg/ml doxycycline (dox) overnight and stained for MR1 surface expression. The BEAS-2B MR1 KO parent cell line is shown for comparison. Data are representative of three independent experiments. B. Clonal cell lines derived from the polyclonal parent cell line were induced with 2 μg/ml of dox (dark colors) or not (light grey) and either treated with 100 μM of 6-FP (filled histograms) or an equal amount of solvent control 0.01 M NaOH (empty histograms) overnight. Data are representative of three independent experiments. The same unstained BEAS-2B WT control is included in each graph for reference. Calibration beads were included in each experiment. C. MR1-GFP expression was measured over time after removal of 2 μg/ml doxycycline. Data are pooled from three independent experiments, each cell line normalized to its starting level in each experiment and shown as mean with standard deviation (SD). Best fit parameters for straight lines were determined by least-squares regression. The fit of one curve fit to all the data sets was compared with the fit of individual curves fit to each data set using the extra sum-of-squares F test. * = p ≤ 0.05 ** = p ≤ 0.01 *** = p ≤ 0.001 **** = p ≤ 0.0001. For full test results see Table S1. D. Whole cell lysates from cell lines induced with 2 μg/ml dox overnight were analyzed for expression of MR1-GFP and loading control Vinculin (VINC) by WB. Primary antibodies were from different species and detected in parallel with species-specific secondary antibodies conjugated to distinct IRDyes and both channels exported as greyscale. Molecular weight markers are indicated in kDa on the left. Data are representative of three independent experiments. See Figure S1 for remaining blots. E. The region around the sgRNA target site was analyzed by next-generation sequencing and mutations resulting in intact reading frames are summarized. See Figure S2 for complete sequence analysis. RF = reading frame; aa = amino acid; GeoMFI = geometric mean fluorescence intensity.

**Figure 2:**
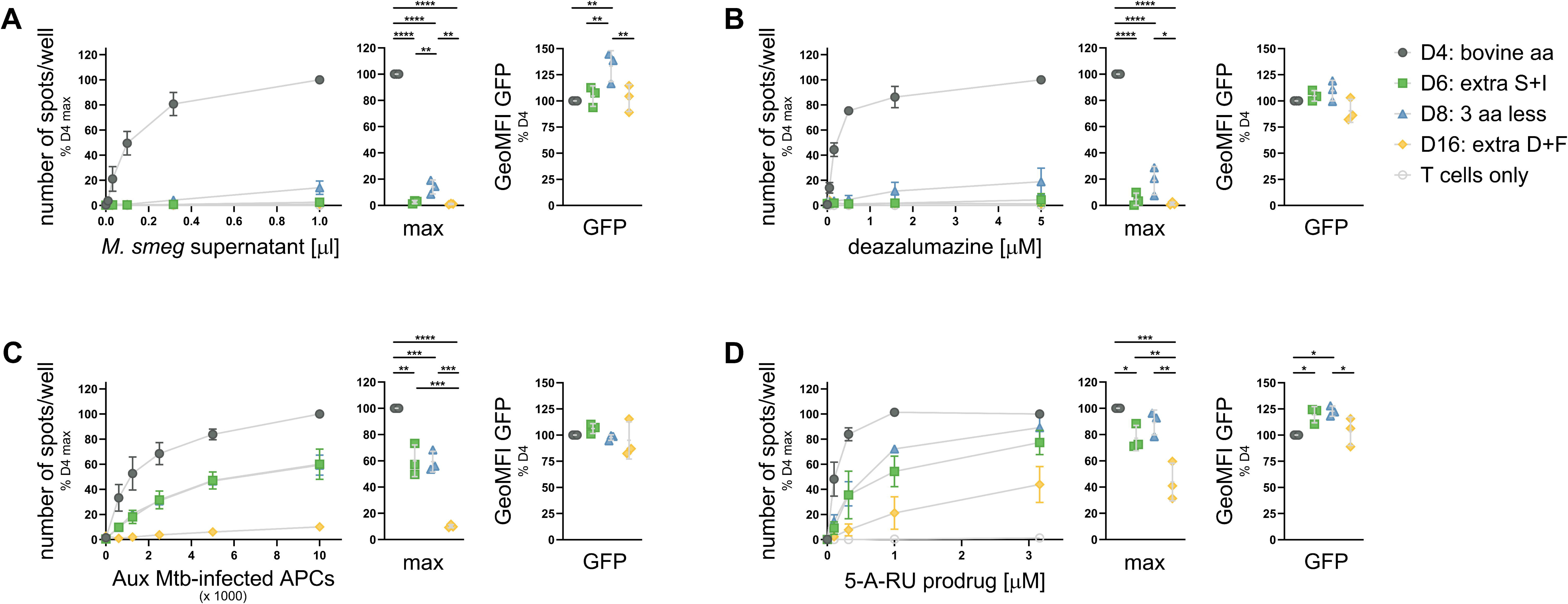
MR1 mutants have a defect in presenting exogenous ligands but present ligands derived from intracellular infection and endosomal processing. MR1-GFP expression was induced with adjusted dox concentrations (see Figure S3 and text) overnight and cells were used as APCs in IFNγ ELISpots. Cells were pre-incubated with indicated dilutions of M. smeg supernatant (A), deazalumazine (B), or 5-A-RU prodrug (D) for at least one hour before addition of MAIT cell clone D481-C7. APCs in C were infected with auxotroph Mtb (Aux Mtb) overnight and diluted as indicated. IFNγ responses at the highest antigen concentrations are additionally shown as dot plots. MR1-GFP expression at the time of the ELISpot was measured and is shown to the right of each plot. Data are pooled from three independent experiments, normalized to D4 at the highest antigen concentration and shown as mean with SD. Experimental groups were compared by repeated-measures ANOVA with Tukey’s multiple comparisons test and statistically significant differences are indicated. * = p ≤ 0.05 ** = p ≤ 0.01 *** = p ≤ 0.001 **** = p ≤ 0.0001. aa=amino acid; GeoMFI = geometric mean fluorescence intensity.

A time course analysis of GFP expression levels after removal of dox indicated that total MR1 protein stability differed in at least one of the cell lines (nonlinear regression analysis; p = 0.0126; Figure 1C and Table S1). Subsequent pairwise comparisons supported the conclusion that MR1 in D8 was degraded more slowly than in the other cell lines in the time frame investigated. Western blot (WB) analysis further confirmed full-length expression of the MR1-GFP fusion protein in all four cell lines (Figures 1D and S1). Thus, neither decreased protein stability nor integrity explain the reduced MR1 cell surface expression in three of the four clonal cell lines.

The sgRNA used to generate the parent BEAS-2B MR1 KO cell line targets a region within exon 2 of the MR1 gene [26]. On the protein level, this translates to amino acids L10 – I16 in the loop connecting two of the strands forming the beta sheet that constitutes the base of the MR1 antigen binding cleft [7]. The loop itself is outside the antigen binding cleft. To better understand the nature of the mutations within our panel of cell lines, we performed next generation sequencing on the region within the lentiviral insertion that was targeted by the sgRNA. Analysis with CRISPResso 2.0 [29] identified multiple gene-edited integrations of the MR1-GFP cassette in each of the four cell lines (Figure S2). Importantly, only one integration in each cell line was edited in a manner that preserved the reading frame except for D6 which had two integrations with in-frame mutations (Figures 1E and S2). The substitutions in the D4 sequence resulted in an amino acid change from valine to isoleucine at position 12. Multiple MR1 homologs in other species, including cattle and swine, have an isoleucine in this position [30], explaining the wildtype (WT)-like phenotype of D4 and allowing us to utilize this cell line as a control. The two insertions with a preserved reading frame in D6 featured a 24-basepair deletion (D6del) and a 6-basepair insertion which resulted in an additional polar serine and an additional isoleucine in the loop (D6ins). The D8 sequence has a 9 nucleotide deletion, resulting in the loss of G11, V12, and S13. D16 has insertions and substitutions which introduce a bulky Phenylalanine and a negatively charged Aspartate (Figures S2 and 1E). The diverse MR1 surface expression phenotypes and the intriguing sequence analysis led us to hypothesize that these mutant cell lines may differ functionally.

### MR1 mutants have a defect in presenting exogenous ligands but present ligands derived from intracellular infection and endosomal processing

We next investigated the antigen presentation capacity of the BEAS-2B MR1 KO tet MR1-GFP clonal cell lines using interferon-γ (IFNγ) production by two MR1-restricted, TRAV1-2^+^ MAIT cell clones established in our laboratory (D481-C7 [31] and D426-G11 [10]) as the readout. Since the four cell lines expressed different levels of MR1-GFP when induced with the same amount of dox (Figure 1B), we titrated the antibiotic and determined dox concentrations for each cell line that resulted in comparable MR1-GFP expression (Figure S3). Importantly, when pre-treated with the adjusted dox concentrations, the four clonal cell lines still expressed different levels of MR1 at the cell surface (compare Figures 1B and Figure S4). When testing the mutant cell lines for their ability to present antigen to the D481-C7 MAIT cell clone, we found a striking dichotomy: while D6 and D8 were greatly impaired in their capacity to present soluble, exogenous antigen such as M. smeg supernatant or the synthetic ligand deazalumazine (DZ) [32] (Figure 2A and B), they retained the ability to present antigen derived from intracellular infection (Figure 2C), albeit with a lower efficiency than D4 (Figure 2C). D6 and D8 also presented a 5-Amino-6-D-ribitylaminouracil (5-A-RU) prodrug that required endosomal processing [33] and at the highest antigen concentration their antigen presentation capacity was minimally different from that of D4 (Figure 2D). The 5-A-RU prodrug was the only antigen presented by D16. Importantly, flow cytometric analysis at the time of IFNγ enzyme-linked immunosorbent spot (ELISpot) confirmed that the adjusted dox concentrations resulted in MR1-GFP expression levels in D6, D8, and D16 that were not lower than in D4 (Figure 2). Thus, the reduced antigen presentation is not explained by lower MR1 expression levels. Similar results were obtained with MAIT cell clone D426-G11 (Figure S5). We also tested 5-OP-RU, live M. smeg bacteria, Bacillus Calmette–Guérin (BCG) infection, and photolumazine I (PLI) using the same dox concentration for all four cell lines (Figure S6). As with the normalized MR1-GFP expression, presentation of soluble, exogenous ligands was strongly reduced when MR1 was mutated in the antigen presenting cells (APCs) whereas MAIT cells were activated in response to cell lines infected with live bacteria (Figure S6).

To establish which of the MR1 mutants expressed in D6 was responsible for the cell surface expression and functional phenotypes, we expressed the in-frame MR1 sequence from D4, the D6del sequence, and the D6ins sequence in the BEAS-2B MR1 KO parental cell line by transient transfection. Despite comparable transfection efficiency and full-length MR1 protein expression from all plasmids, D6del MR1 was undetectable at the cell surface and presented neither M. smeg supernatant nor Mtb-derived antigen or the 5-A-RU prodrug (Figure S7). The D6ins MR1, on the other hand, phenocopied the D6 cell line (Figure S7). Thus, we concluded that D6del MR1 was non-functional and did not contribute to the surface translocation and antigen presentation seen in the D6 cell line. Interestingly, a V12S mutant of MR1 similarly phenocopied the D6 cell line, allowing us to attribute the effects seen to the presence of a polar residue at position 12 rather than the addition of two amino acids (Figure S7).

### Mutated MR1 does not accumulate in vesicular compartments

We were particularly intrigued by the D6 cell line as MR1 in this clonal line was clearly able to bind ligand and translocate to the cell surface but was strongly impaired in the presentation of exogenous antigens to MAIT cells. Thus, we focused our efforts on the comparison between D4 and D6. Our lab has previously shown that in BEAS-2B cells, GFP-tagged MR1A resides both in the ER and in an endosomal compartment labeled by Rab7 [20]. Since the mutated MR1 expressed by D6 was not detectable at the cell surface at baseline (Figure 1B), we hypothesized that it may not be able to leave the ER in the absence of a stabilizing ligand. Indeed, we found greatly reduced numbers of MR1-GFP^+^ vesicles in the D6 cell line, with the MR1-GFP signal distributed in a reticular pattern only (Figure 3A and B). By contrast, MR1 in D4 trafficked to vesicular compartments and a proportion of these co-stained with the late endosomal marker Rab7 as reported for WT MR1-GFP (Figure 3, [20]).

**Figure 3:**
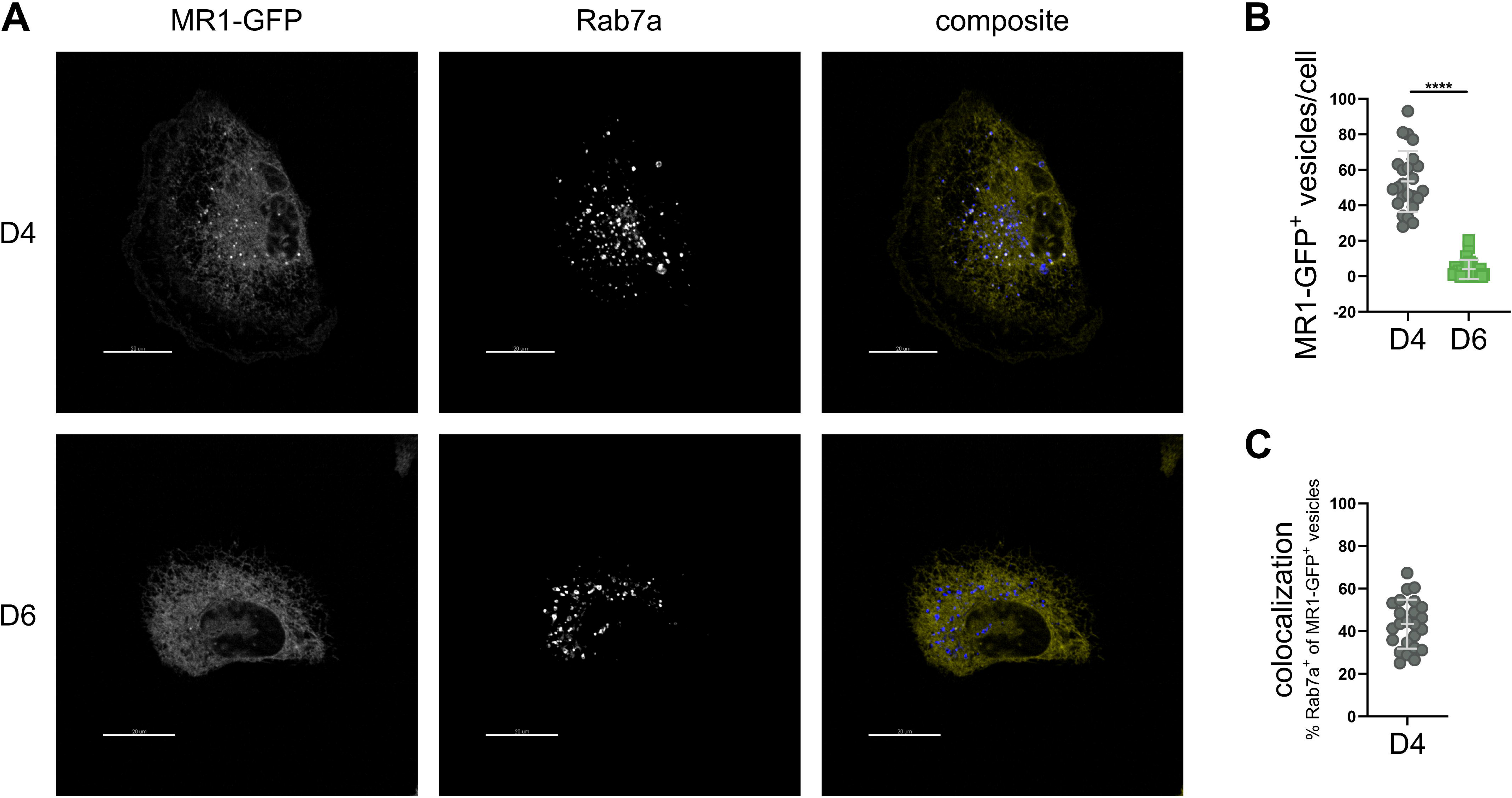
Mutated MR1 does not accumulate in vesicular compartments at baseline. Clonal cell lines D4 and D6 were induced to express MR1-GFP with 1 μg/ml and 2 μg/ml of dox, respectively, and treated with CellLight BacMam 2.0 for late endosomes. A. Representative images. Scale bar indicates 20 μm. B. Enumeration of MR1-GFP^+^ vesicles per cell. C. Colocalization of MR1-GFP^+^ vesicles and Rab7a^+^ vesicles. Data in B and C represent 24 cells from 24 images for each cell line, pooled from three independent experiments. Experimental groups in B were compared by two-tailed, unpaired t-test. * = p ≤ 0.05 ** = p ≤ 0.01 *** = p ≤ 0.001 **** = p ≤ 0.0001.

### Mutated MR1 differentially interacts with calnexin, B2M, HLA-Ia, TPP1, OLFML2A and SQSTM1/p62

Differential antigen presentation pathways dependent on distinct chaperones and trafficking proteins have been proposed for ER-based versus endosomal loading of MR1 [23, 34]. Thus, we hypothesized that MR1 in the D6 mutant may interact differentially with co-factors required for endosomal antigen presentation, but not ER-based presentation of exogenous MR1 ligands. To identify such interaction partners, we performed co-immunoprecipitation of MR1-GFP in D4 and D6 in the presence of 6-FP, followed by mass spectrometry-based proteomic analysis. This allowed the identification of MR1-interacting proteins in a semi-quantitative way using label-free mass spectrometry (Figure 4). Using on-bead Trypsin digestion and a data-dependent acquisition (DDA) approach, we initially found more calnexin (protein: CALX, gene: CANX) in the D6 samples whereas MR1 in D4 preferentially associated with B2M (Figure 4A, Table S4). To confirm and extend the analysis, we repeated the experiment using in-gel digestion and data-independent acquisition (DIA) (Figure 4B and C, Table S5). This more quantitative method confirmed the differential and opposite associations of MR1 in D4 and D6 with B2M and CANX, respectively (Figure 4B and C). In addition, the second mass spectrometry-based analysis showed that MR1 in D6 associated with HLA-Ia proteins, the lysosomal tripeptidyl-peptidase 1 (TPP1; also known as CLN2) [35], the pleiotropic olfactomedin-like protein 2A (OLFML2A) [36, 37] and the autophagy mediator sequestosome 1 (SQSTM1; also known as p62) [38] more frequently than MR1 in D4. Analysis of the lysates from the three independent immunoprecipitation experiments analyzed by DIA for B2M, CANX, and SQSTM1/p62 by WB supported the mass spectrometry results (Figure 4D).

**Figure 4:**
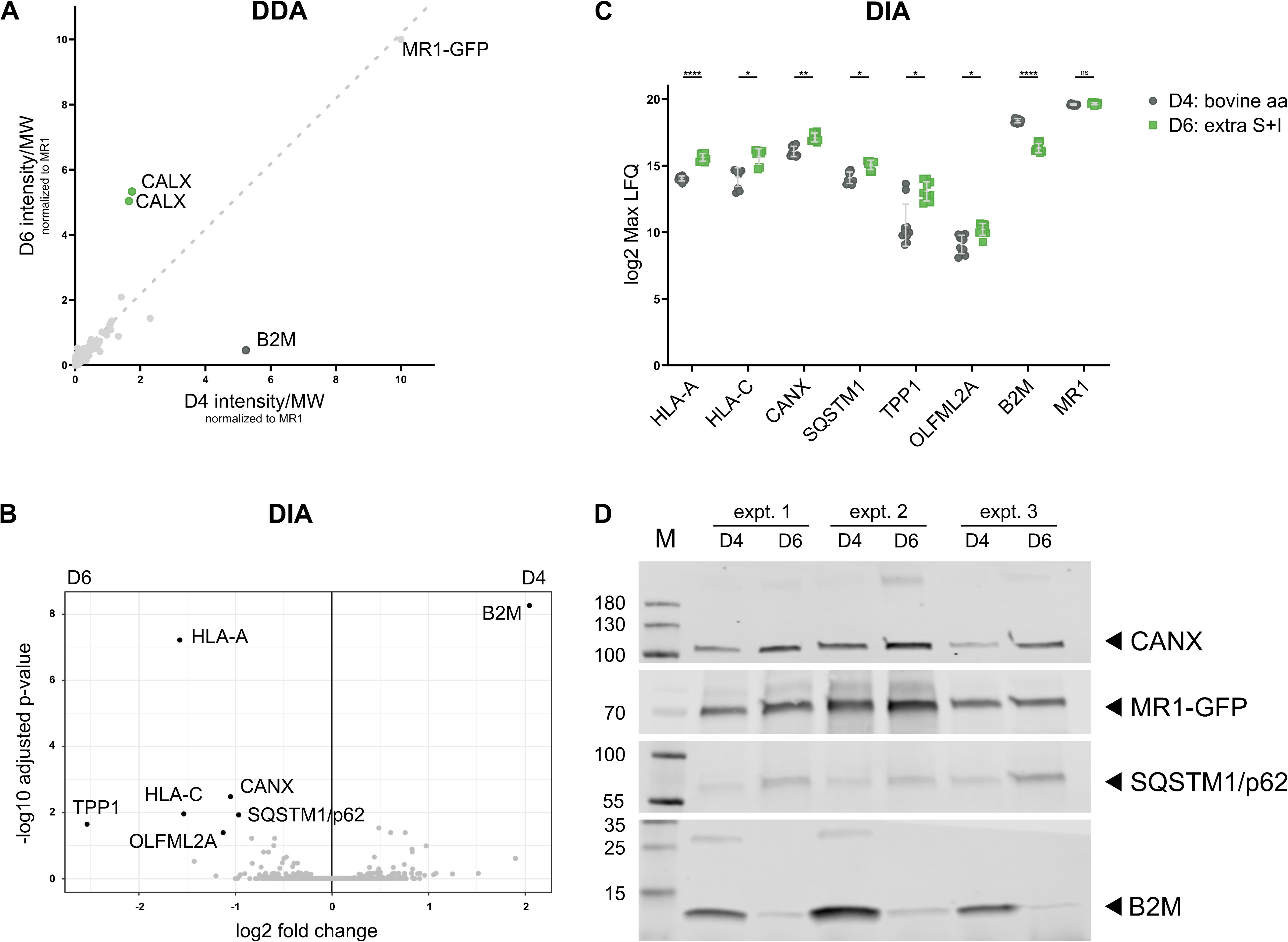
Mutated MR1 differentially interacts with calnexin, B2M, HLA-Ia, TPP1, OLFML2A and SQSTM1/p62. Expression of MR1-GFP was induced in clonal cell lines D4 and D6 with 10 μg/ml of dox overnight before incubation with 20 μg/ml dox and 100 μM 6-FP overnight. MR1-GFP was immuno-precipitated and bound proteins were analyzed by mass spectrometry in DDA mode (A) or DIA mode (B+C). A. Average size-adjusted intensity of proteins identified in both D4 and D6 normalized to MR1-GFP intensity in each sample. Data in A are pooled from three independent experiments. Data in B and C are pooled from three independent experiments with three or four technical replicates each. Each dot in C represents one mass spectrometry injection for proteins with p adjusted ≤ 0.01 and |log2 fold-change| ≥ 1. For statistical analysis see mass spectrometry methods section. * = p ≤ 0.05 ** = p ≤ 0.01 *** = p ≤ 0.001 **** = p ≤ 0.0001. For full lists of identified proteins see Tables S4 and S5. D. The lysates of the three experiments shown pooled in B and C were analyzed by WB. CANX and B2M were analyzed on one gel with the membrane cut horizontally before primary antibody incubation. Both the red and the green channel were exported in greyscale to allow visualization of the molecular weight marker. MR1-GFP and SQSTM1/p62 were analyzed on another gel with primary antibodies from different species. Species-specific secondary antibodies conjugated to distinct IRDyes were used and each channel was exported individually in greyscale. DDA = Data dependent acquisition; DIA = Data independent acquisition; aa = amino acid; MW = molecular weight.

### Differential association with SQSTM1/p62 does not explain the functional differences in MR1-mediated antigen presentation

SQSTM1/p62 is a selective autophagy receptor for ubiquitinated substrates [38]. As such, SQSTM1/p62 is critical for the clearance of intracellular bacterial pathogens by xenophagy [39, 40], which provided a rationale to further investigate this protein in the context of MR1-mediated presentation of mycobacterial antigens. Furthermore, a recent preprint publication suggested a role for autophagy proteins, including SQSTM1/p62, in MR1-mediated presentation of antigens derived from fixed bacteria [41]. Thus, we transfected small interfering RNAs (siRNAs) targeting the SQSTM1/p62 transcript into both the D4 and the D6 cell lines. We confirmed reduced expression of SQSTM1/p62 in our experimental setup at the protein level by WB (Figure S8) and found that despite successful knock-down of SQSTM1/p62, presentation of antigens from M. smeg supernatant, the 5-A-RU prodrug, and live infection with auxotroph Mtb (Aux Mtb) to an MR1-restricted T cell clone was indistinguishable from those in cells that had received a scrambled control siRNA (Figure 5A and B). We repeated the experiment in BEAS-2B WT cells to test for a role of SQSTM1/p62 under physiological expression of MR1 and again found no effect (Figure 5 C and D). Thus, we conclude that despite its importance in xenophagy and its evident interaction with MR1 in both D4 and D6, SQSTM1/p62 is not required for the presentation of either exogenous or endosomal MR1 ligands. We hypothesize that the increased association of SQSTM1/p62 with MR1 in D6 may instead be explained by the autophagy receptor being involved in the clearance of the non-functional D6del MR1 molecules.

**Figure 5:**
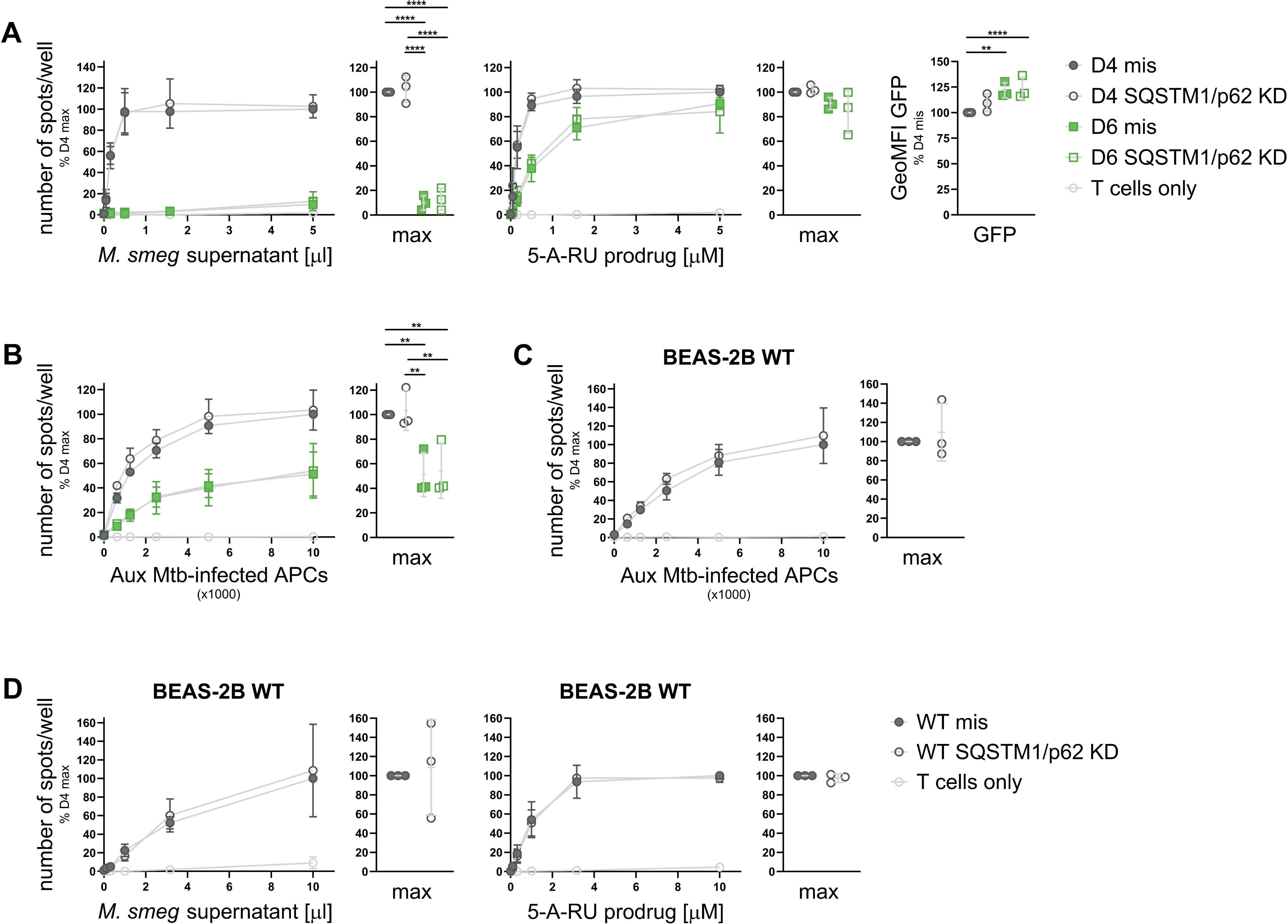
SQSTM1/p62 is not required for MR1-mediated antigen presentation. Clonal cell lines D4 and D6 (A+B) or BEAS-2B WT cells (C+D) were transfected with missense (mis, filled symbols) siRNA or siRNA targeting SQSTM1/p62 (KD, empty symbols) after inducing MR1-GFP expression with adjusted dox concentrations (see Figure S3, A+B only) and used as APCs in IFNγ ELISpots with the indicated antigens. MR1-GFP expression was measured by flow cytometry at the time of the ELISpot and is shown to the right in A. Data are pooled from three independent experiments, normalized to D4 at the highest antigen concentration and shown as mean with SD. IFNγ responses at the highest antigen concentrations are additionally shown as dot plots. Experimental groups in A and B were compared by repeated-measures ANOVA with Tukey’s multiple comparisons test and statistically significant differences are indicated. Comparisons in C and D were performed with two-tailed, paired t-tests. * = p ≤ 0.05 ** = p ≤ 0.01 *** = p ≤ 0.001 **** = p ≤ 0.0001. SQSTM1/p62 KD was confirmed by WB as shown in Figure S8. GeoMFI = geometric mean fluorescence intensity.

### MR1 mutant D6 binds B2M with lower apparent affinity

Research in the MR1 field established early on that B2M was required for the presentation of MAIT cell antigens [42, 43]. Since less B2M was present in immunoprecipitation samples from D6 than D4, we sought to rescue the functional defect in D6 by over-expressing B2M. To this end, we transiently transfected both D4 and D6 with a plasmid encoding B2M under the control of a constitutive cytomegalovirus (CMV) promoter. The complementary DNA (cDNA) sequence of B2M was followed by an internal ribosomal entry site and GFP to assess transfection efficiency by microscopy before induction of MR1-GFP with dox. Over-expression of B2M did not allow D6 to present M. smeg supernatant and did not result in increased presentation of the 5-A-RU prodrug (Figure 6A). B2M overexpression was confirmed by WB (Figure S9). This finding is consistent with the hypothesis that the MR1 mutation in D6 resulted in a reduced affinity for B2M rather than reduced availability of B2M. Thus, providing more B2M did not remedy its reduced ability to form a full trimer with ligand and light chain. Since it is not possible to refold recombinant MR1 without B2M for biochemical affinity measurements, we adapted a method used by Hein et al. [44] to measure the dissociation of B2M from MR1 in immunoprecipitates from D4 and D6. Using this technique, we found that B2M more rapidly disappeared from the fractions containing the magnetic resin bound to the MR1 molecules immunoprecipitated from D6 compared to those from D4 and, conversely, appeared in the unbound fractions sooner than for D4 (Figures 6C and S10). Although our data did not allow us to compute best fit values for the rate of decay for D4 with confidence (Table S6), the comparison of the model curves supports the conclusion that B2M dissociated more rapidly from MR1 immunoprecipitated from D6 than D4. This is consistent with a lower affinity for B2M. Of note, association with CANX and B2M has been reported to be mutually exclusive for certain HLA-Ia molecules [45, 46]. Thus, an alternative explanation for the inverse associations of D4 and D6 with CANX and B2M may be an increased affinity of D6 for CANX.

**Figure 6:**
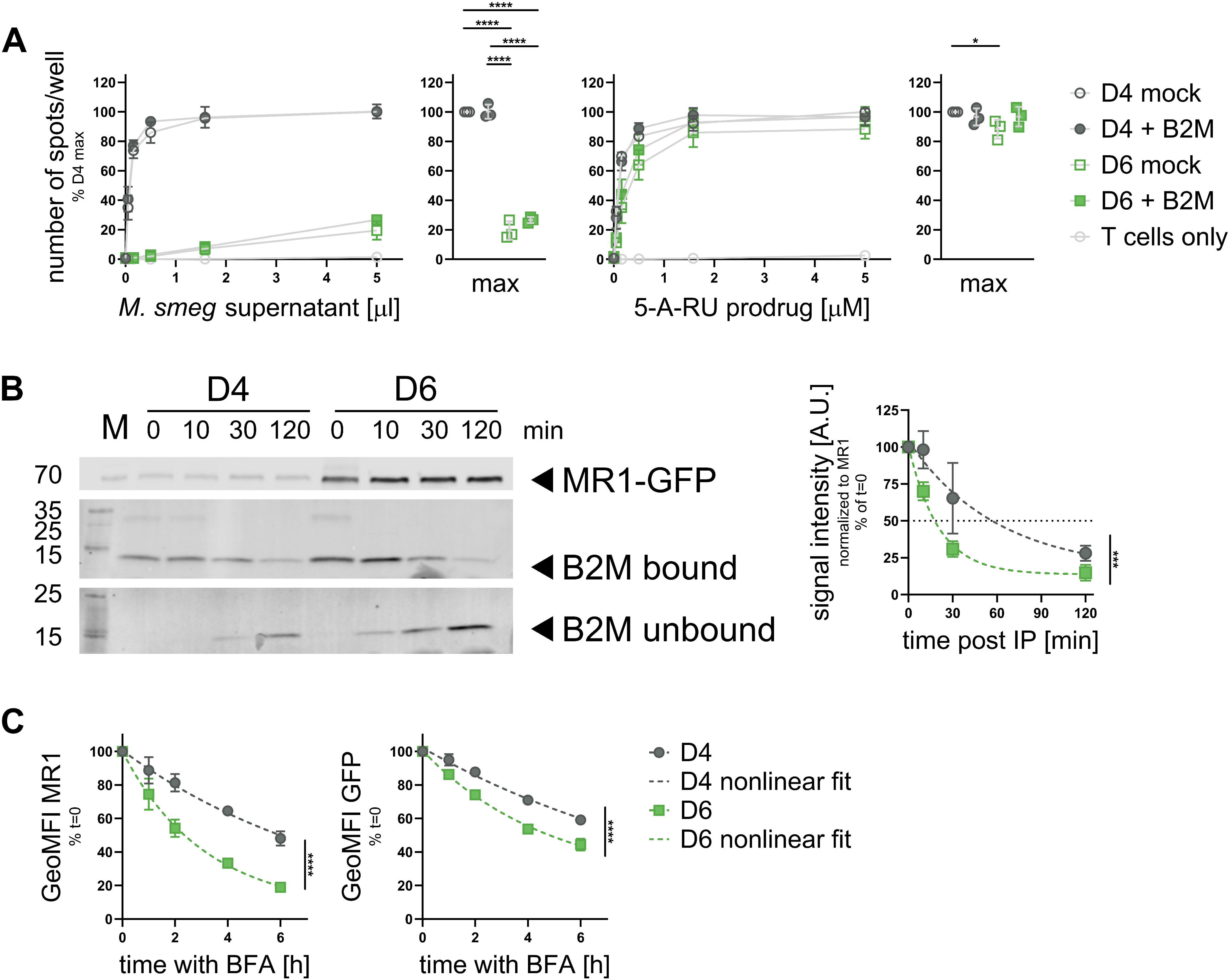
Mutant D6 binds B2M with lower apparent affinity. A. Clonal cell lines D4 and D6 were transiently transfected to overexpress B2M (full symbols) or mock transfected (empty symbols) before use as APCs in an ELISpot. Data are pooled from three independent experiments, normalized to D4 at the highest antigen concentration and shown as mean with SD. IFNγ responses at the highest antigen concentrations are additionally shown as dot plots. Experimental groups were compared by repeated-measures ANOVA with Tukey’s multiple comparisons test and statistically significant differences are indicated. * = p ≤ 0.05 ** = p ≤ 0.01 *** = p ≤ 0.001 **** = p ≤ 0.0001. B2M over-expression was confirmed by WB as shown in Figure S9. B. MR1-GFP was immunoprecipitated from clonal cell lines D4 and D6 induced to express MR1-GFP with 0.5 or 8 μg/ml dox, respectively, overnight followed by incubation with the same concentrations of dox plus 100 μM 6-FP overnight. Immunoprecipitated samples were incubated at 37°C for the indicated time periods and MR1 bound to the beads, B2M bound to the beads, and B2M in the supernatant (unbound) were measured by WB. A representative blot is shown on the left and pooled data from three experiments, normalized to MR1 for each sample and t=0 for each cell line is shown on the right as mean with SD. Primary antibodies were detected in parallel with species-specific secondary antibodies conjugated to IRDye800 and both channels exported as greyscale to visualize the molecular weight markers. The 800 channel was exported individually for MR1-GFP. See Figure S10 for the remaining blots. D. D4 and D6 were induced to express MR1-GFP with 4 and 8 μg/ml dox, respectively, overnight followed by incubation with the same concentrations of dox with 100 μM 6-FP overnight to bring MR1 to the cell surface before incubation with brefeldin A (BFA) for the indicated amounts of time. MR1 surface levels (left) and total MR1 expression (right) were measured by flow cytometry. Best fit parameters for one-phase exponential decay curves were determined by least-squares regression in B and C. The fit of one curve fit to both data sets was compared with the fit of individual curves fit to each data set using the extra sum-of-squares F test. For full test results see Tables S6 – S8. * = p ≤ 0.05 ** = p ≤ 0.01 *** = p ≤ 0.001 **** = p ≤ 0.0001. IP = immunoprecipitation; A.U. = arbitrary units; GeoMFI = geometric mean fluorescence intensity.

B2M affinity has been linked to the stability of murine MHC class Ia:peptide complexes at the cell surface [47]. Adapting the brefeldin A (BFA) assay used by Montealegre et al., we investigated the stability of fully folded trimers of MR1, B2M and ligand at the cell surface after treatment with BFA, which prevents anterograde transport via the Golgi apparatus and thereby halts supply of new MR1 molecules to the cell surface. After 6 hours, approximately 50% of the MR1 molecules recognized by the conformation-dependent antibody clone 26.5 on the cell surface remained in D4 (Figure 6D, left). Only 20% of the fully folded MR1 remained at the cell surface in D6 at the same time point. Of note, the anti-MR1 antibody clone used here is specific for the full trimer of MR1, B2M, and a ligand and, thus, the assay does not allow us to distinguish between internalization of the protein and loss of the epitope due to dissociation of B2M, the ligand, or both. Therefore, we cannot investigate the possibility that MR1 persists at the cell surface as ligand-free heavy chains as can be the case for MHC class Ia [48]. Interestingly, total MR1 protein levels as measured by the intensity of GFP also decreased slightly over the 6-hour experiment and again, this occured faster in D6 compared to D4 (Figure 6D, right). For both MR1 surface levels and total MR1-GFP expression we were unable to compute complete confidence intervals for some of the nonlinear model parameters (Tables S7 and S8), but comparisons of the model fits nevertheless indicated different curves for D4 and D6, further supporting the idea that D6 has a lower affinity for B2M.

### Loss of B2M differentially affects exogenous and endosomal antigen presentation pathways when MR1 is not limiting

Having established that overexpression of B2M did not rescue the inability of D6 to present exogenous ligands (Figure 6A and B), we next tested whether loss of B2M in D4 would phenocopy the effects seen in D6. We used siRNA-mediated knock-down (KD) and confirmed the loss of B2M on the protein level by WB (Figure S11). As expected, loss of B2M in D4 resulted in a significant decrease of the presentation of M. smeg supernatant (Figure 7A). However, to our surprise, the presentation of 5-A-RU prodrug via the endosomal pathway was not impacted in D4 even when B2M expression was knocked-down. Presentation of antigen derived from intracellular infection with Aux Mtb was similarly unaffected by B2M knock-down in D4 (Figure 7B). Interestingly, loss of B2M did abrogate the ability of D6 to present both endosomal antigens (Figure 7A and B).

**Figure 7:**
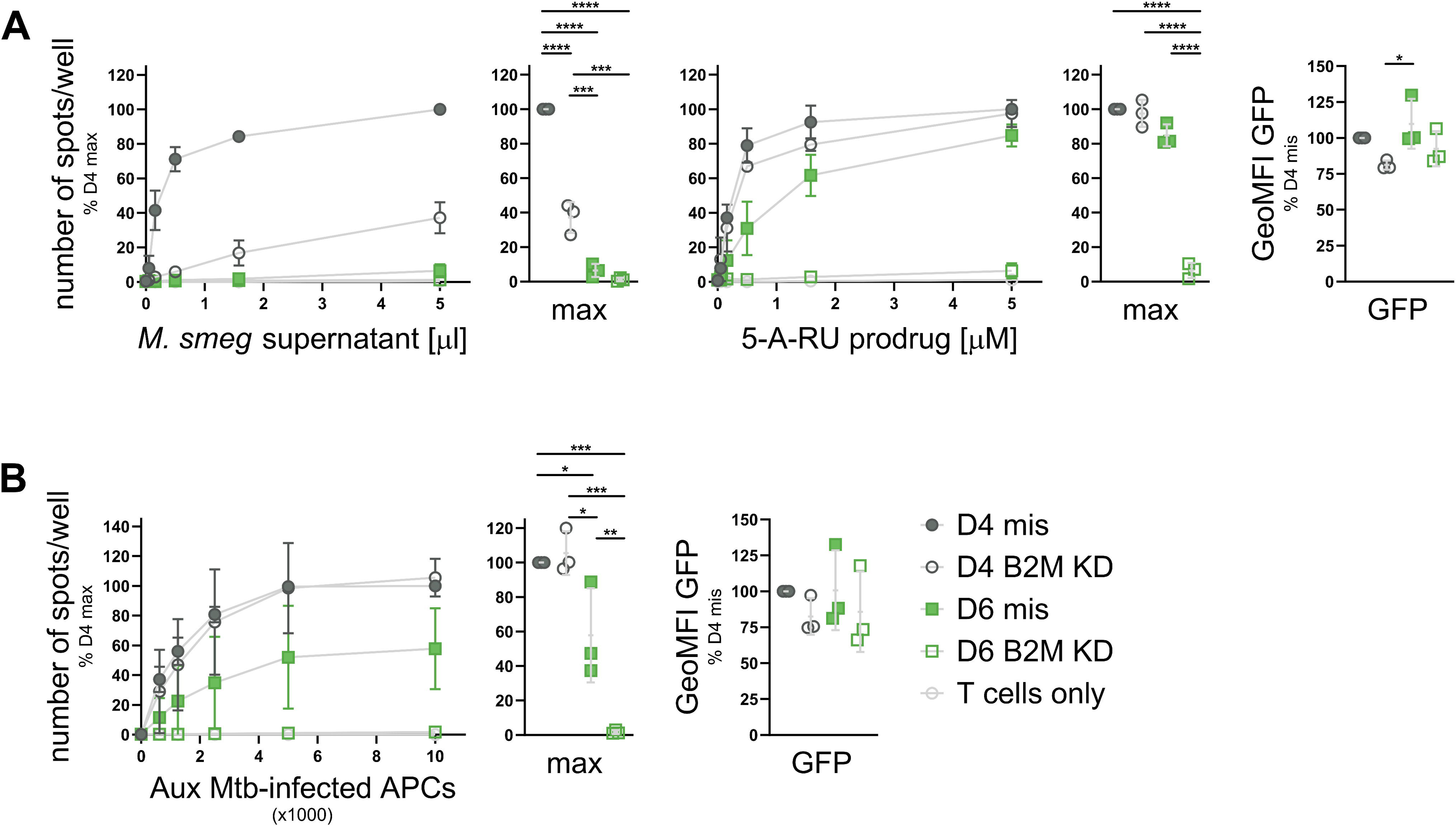
Knock-down of B2M differentially affects exogenous and endosomal antigen presentation pathways. Clonal cell lines D4 and D6 were transfected with missense (mis, filled symbols) siRNA or siRNA targeting B2M (KD, empty symbols) before induction with adjusted dox concentrations (see Figure S3) and use as APCs in IFNγ ELISpots with the indicated antigens. Dox concentrations during Aux Mtb infection were 0.4 and 0.8 μg/ml, respectively. MR1-GFP expression was measured by flow cytometry at the time of the ELISpot and shown to the right. Data are pooled from three independent experiments, normalized to D4 at the highest antigen concentration and shown as mean with SD. IFNγ responses at the highest antigen concentrations are additionally shown as dot plots. Experimental groups were compared by repeated-measures ANOVA with Tukey’s multiple comparisons test and statistically significant differences are indicated. * = p ≤ 0.05 ** = p ≤ 0.01 *** = p ≤ 0.001 **** = p ≤ 0.0001. B2M KD was confirmed by WB as shown in Figure S11. GeoMFI = geometric mean fluorescence intensity.

As in the D6 cell line, B2M KD in BEAS-2B WT cells significantly abrogated presentation of both exogenous and endosomal antigen (Figure 8A). We considered three differences between BEAS-2B WT cells and the D4 cell line which may explain the differential dependence on B2M expression: firstly, unlike endogenous MR1, the MR1 in D4 is tagged with GFP. We reasoned that any effects of the GFP tag should equally affect MR1 in the other clonal cell lines and, thus, discarded this explanation. Secondly, MR1 expression levels in the clonal cell lines are higher than in BEAS-2B cells as indicated by the ability to detect MR1-GFP by Western blot (Figure 1D). Thirdly, the clonal cell line D4 may carry an unknown off-target mutation which has unexpected consequences for MR1 trafficking and antigen presentation unique to this cell line. To investigate the latter two possibilities, we repeated the B2M KD experiment in a previously published polyclonal BEAS-2B cell line, which constitutively over-expresses MR1-GFP under a minimal CMV promoter (OE, [49, 50]). As in D4, B2M KD in these cells reduced the presentation of exogenous antigen, but not endosomal antigen (Figure 8B). This confirmed that the apparent lack of B2M dependence of endosomal antigen presentation was not unique to D4 but might indeed be attributable to MR1 overexpression. The BEAS-2B MR1 KO cell line from which both D4 and D6 were derived was included as a negative control and did not induce MAIT cell activation in any of the assays (Figure 8B). The lack of endosomal antigen presentation in the parental cell line makes it appear unlikely that the D4 cell line inherited unexpected mutations which render this pathway less sensitive to B2M levels.

**Figure 8:**
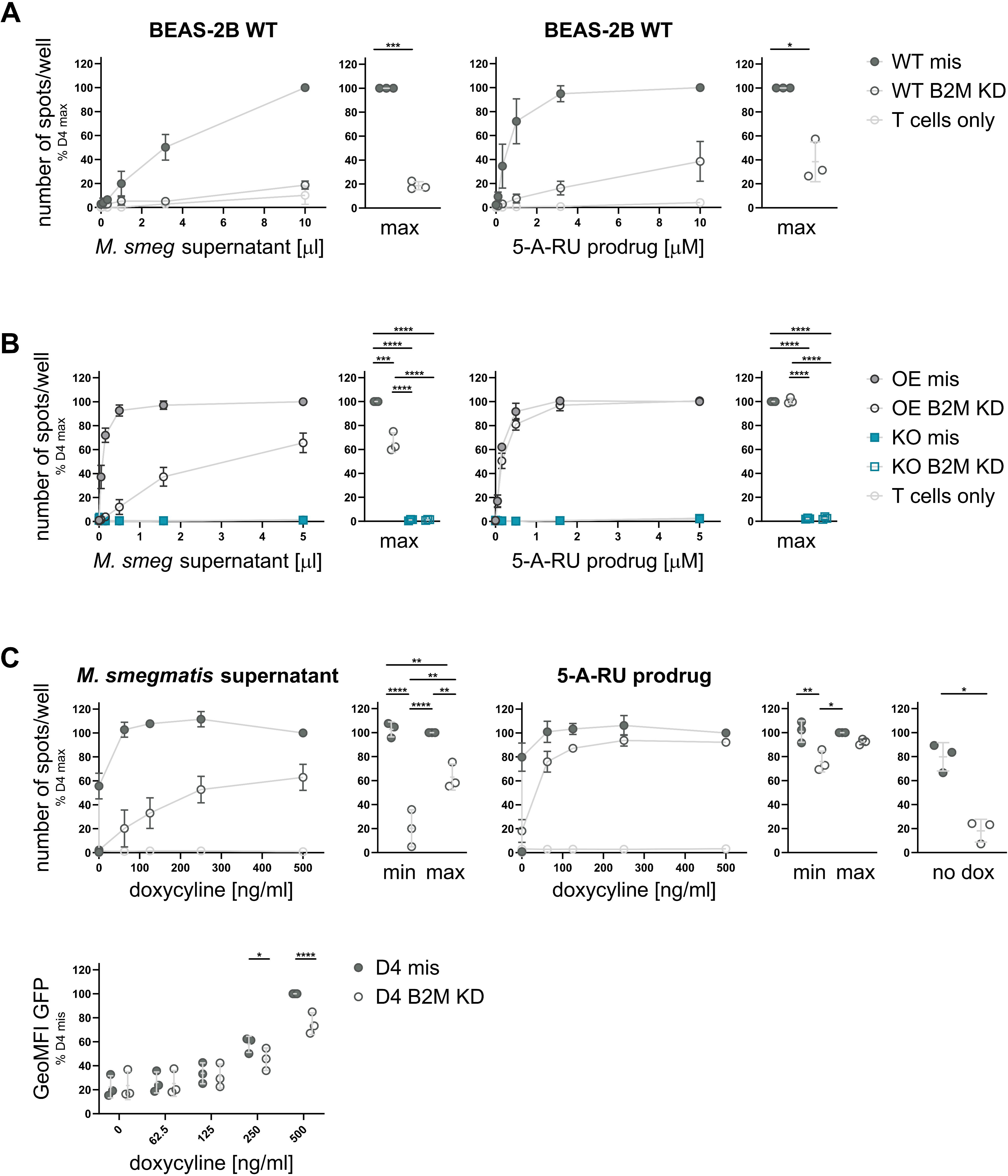
The effect of B2M knock-down depends on MR1 expression levels. BEAS-2B WT cells (A), or BEAS-2B MR1 KO (KO) and BEAS-2B MR1 overexpressing (OE) cells (B) were transfected with missense (mis, filled symbols) siRNA or siRNA targeting B2M (KD, empty symbols) before induction with adjusted dox concentrations (see Figure S3) and use as APCs in IFNγ ELISpots with the indicated antigens. C. B2M was knocked-down in the clonal cell line D4 as in A and B and MR1-GFP expression was induced with the indicated concentrations of dox before use as APCs in IFNγ ELISpots with the indicated antigens. MR1-GFP expression was measured by flow cytometry at the time of the ELISpot and is shown to the right. Data in all panels are pooled from three independent experiments, normalized to D4 at the highest antigen concentration and shown as mean with SD. IFNγ responses at the highest antigen concentrations (A+B) or the highest and lowest dox concentrations (C) are additionally shown as dot plots. Experimental groups were compared by repeated-measures ANOVA with Tukey’s multiple comparisons test for more than two groups or a two-tailed, paired t-test for two groups and statistically significant differences are indicated. * = p ≤ 0.05 ** = p ≤ 0.01 *** = p ≤ 0.001 **** = p ≤ 0.0001. Comparisons of all experimental groups for the flow cytometry analysis in C are shown in Table S9. B2M KD was confirmed by WB as shown in Figure S11. GeoMFI = geometric mean fluorescence intensity.

**Figure 9:**
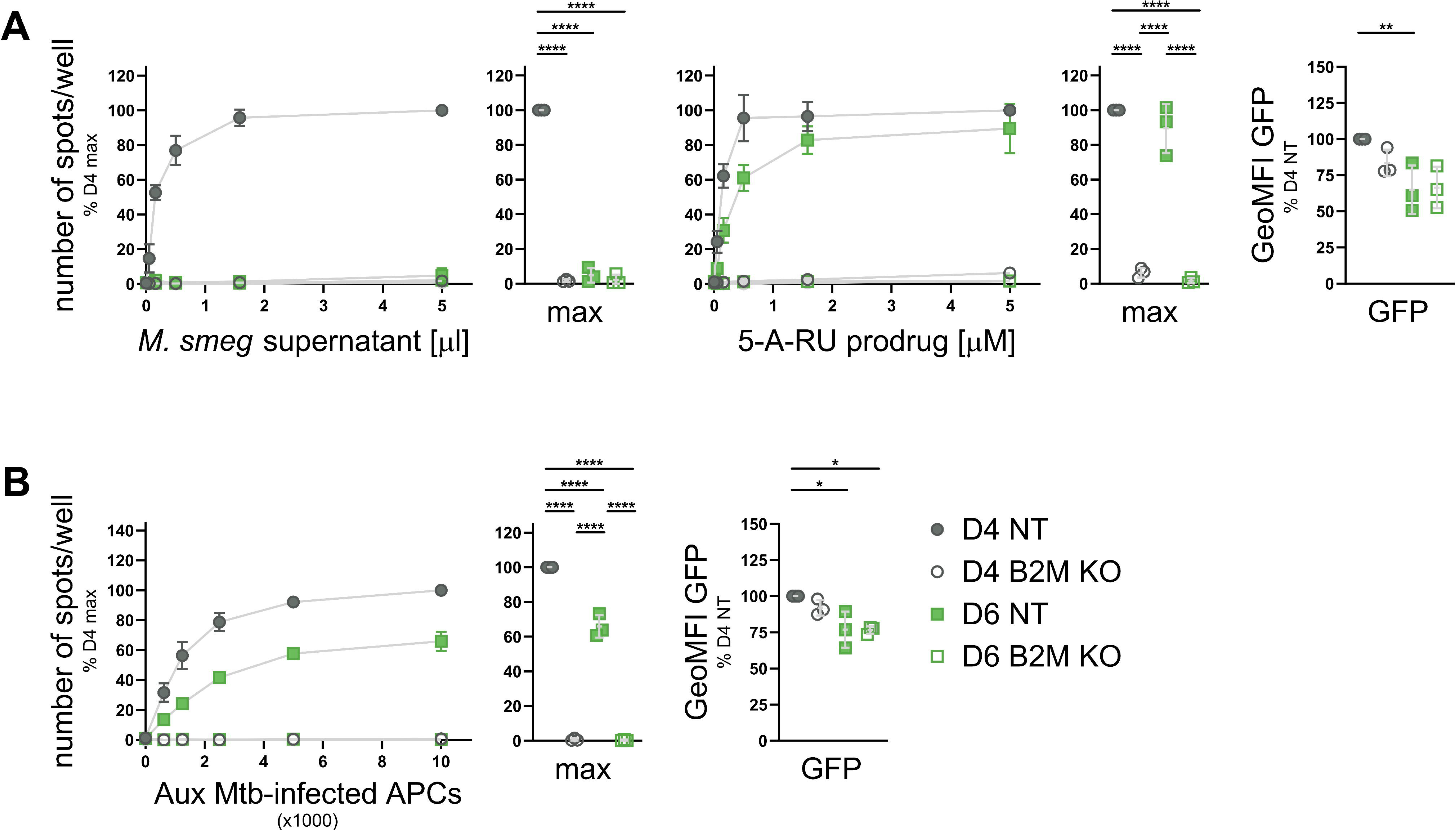
B2M is required for endosomal antigen presentation in the presence of high levels of MR1. Clonal cell lines D4 and D6 cells were electroporated with RNPs containing scrambled non-target control sgRNAs (NT) or three pooled sgRNAs targeting B2M (KO). Single clones were induced to express MR1-GFP with adjusted dox concentrations (see Figure S3) and used as APCs in IFNγ ELISpots with the indicated antigens. MR1-GFP expression was measured by flow cytometry at the time of the ELISpot and is shown to the right. Data are pooled from three independent experiments, normalized to D4 at the highest antigen concentration and shown as mean with SD. IFNγ responses at the highest antigen concentrations are additionally shown as dot plots. Experimental groups were compared by repeated-measures ANOVA with Tukey’s multiple comparisons test and statistically significant differences are indicated. * = p ≤ 0.05 ** = p ≤ 0.01 *** = p ≤ 0.001 **** = p ≤ 0.0001. B2M KO was confirmed by WB, PCR, and qRT-PCR as shown in Figure S12. GeoMFI = geometric mean fluorescence intensity.

To further test the hypothesis that the extent to which endosomal antigen presentation depends on B2M protein levels depends on the expression levels of MR1, we titrated down the amount of doxycycline used to induce MR1-GFP expression in D4. As expected, B2M KD significantly reduced the presentation of exogenous M. smeg supernatant regardless of the level of MR1 expression in D4 (Figure 8C). Presentation of the 5-A-RU prodrug, however, was unaffected at the highest dox concentration, but substantially decreased at the lowest concentration (Figure 8C). Indeed, we found that even without addition of doxycycline, this cell line produced enough MR1-GFP to activate the MAIT cell clone, presumably due to leakiness of the tet promoter system, and this baseline level of antigen presentation was sensitive to B2M KD. Flow cytometric analysis confirmed that the doxycycline titration resulted in varying levels of MR1-GFP expression (Figure 8C). Interestingly, B2M KD resulted in lower MR1-GFP levels at the highest dox concentrations, suggesting that stability of the MR1-GFP protein was dependent on B2M levels. This makes the lack of a functional effect of B2M KD under these conditions even more intriguing.

Finally, we used CRISPR/Cas9 technology to knock out B2M completely in both D4 and D6. As opposed to the siRNA-mediated KD, these cells were unable to express any B2M (Figure S12) and under these conditions, both exogenous and endosomal antigen presentation in D4 was abrogated (Figure 8A and B). We confirmed these results with a second set of B2M KO and non-target (NT)-control clonal cell lines (Figure S13). Thus, endosomal antigen presentation in D4 is still dependent on B2M but appears to require only very low levels of the protein such that the minimal levels remaining upon B2M KD, though undetectable by WB, suffice to support presentation of endosomal antigen.

We conclude that presentation of exogenous antigens is limited by the availability of B2M and, thus, acutely sensitive to reductions in the levels of this protein, whereas the endosomal antigen presentation pathway is limited by MR1 as it is functional even if B2M levels are drastically lowered (as in the knock-down experiments) or B2M is less able to bind MR1 (as in D6) as long as sufficient MR1 molecules are available.

## Discussion

Multiple, complementary pathways have been proposed for the surveillance of diverse subcellular compartments by MR1 [23, 34]. In particular, the cellular mechanisms employed for loading MR1 appear to differ between exogenously supplied antigens and those derived in the context of intracellular infection [20–22]. For example, the SNARE Syntaxin 4 plays a role in MR1-mediated presentation of antigen derived from exogenous M. smeg supernatant, but not Mtb infection [21] whereas the calcium-sensitive vesicular trafficking protein Synaptotagmin 7 selectively promotes presentation of antigens from intracellular microbes [22]. Here we add to the body of literature supporting this model by characterizing a panel of MR1 mutants which are greatly impaired in their ability to present exogenous ligands but retain the ability to present antigens generated in the endosomal compartment. The antigen presentation defect is most profound with synthetic ligands PLI and DZ, which none of the MR1 mutants present to an appreciable extent. The MR1 expressed in clonal cell lines D6 and D8 does present exogenous antigens such as M. smeg supernatant and 5-OP-RU, but only if provided at concentrations much higher than those required for the control cell line D4 to reach maximum stimulatory capacity. Interestingly, the D426-G11 MAIT cell clone appears to be more sensitive to exogenous ligands presented by mutated MR1 as it responds to lower concentrations of 5-OP-RU than the D481-C7 MAIT cell clone. The pattern differs markedly for antigens that are derived from bacterial infection or require endosomal processing. Although the maximum responses measured also tend to be lower than those in the control cell line, D6, D8, and, for some antigens, D16 achieve MAIT cell stimulatory capacity above background even at lower antigen concentrations. This diverse panel of cell lines gave us an opportunity to investigate the cellular mechanisms underlying the different pathways involved in the presentation of exogenous compared to endosomal MAIT cell antigens.

Intriguingly, MR1 in one of the mutated cell lines displayed an increased interaction with the ER chaperone calnexin and a concomitant decrease in association with B2M. It has previously been shown that MHC class Ia molecules bind either calnexin or B2M with binding to the latter releasing the heterodimer from calnexin [45, 46]. Our results suggest that this might be the case for MR1, too, which constitutes a possible mechanism underlying the antigen presentation defect as it may trap MR1 in the ER bound to calnexin in this cell line. While we did not characterize the interaction between calnexin and MR1, the data indicate that such ER retention would likely be due to decreased affinity for B2M as opposed to increased affinity for the chaperone. This hypothesis is further supported by the observation that the loop mutated in MR1 in the D6, D8, and D16 cell lines forms direct contacts with B2M in some fish and chicken MHC class Ia structures [51]. In murine MHC class Ia molecules, this loop is positioned near key contacts within the MHC class Ia:B2M interface [52], but the direct interaction appears to have been lost in the mammalian MHC class Ia isoforms over evolutionary time [51]. Taking into account that MR1 is more closely phylogenetically related to fish and chicken MHC class Ia than human HLA Ia molecules [53], we hypothesize that the loop altered in the panel of MR1 mutants described in this study may directly contact B2M. Indeed, this has been postulated by Dijkstra and colleagues [54] based on published crystal structures of MR1 in complex with B2M and ligand [7, 55]. Consequently, it appears feasible that the introduction or spatial re-arrangement of polar and/or charged residues in this region could interfere with B2M binding to MR1 in the mutant cell lines.

While this would explain the lack of ER-mediated presentation of exogenous antigens, it does not explain why endosomal antigen presentation is not equally affected. Indeed, the molecular determinants that allow D6 to retain the ability to present antigen delivered via endosomal pathways remain to be identified. The fact that D6 presented Mtb antigen and 5-A-RU prodrug despite its reduced binding to B2M indicates that this pathway is less dependent on B2M than presentation of exogenous ligands via the ER. Instead, the results suggest that endosomal antigen presentation is limited by the availability of MR1 molecules themselves as supraphysiological levels of MR1 rescued this function in cells with minimal B2M protein expression. Overexpression of B2M, on the other hand, did not rescue presentation of exogenous MR1 ligands. This could indicate that B2M levels in the transduced cells were not high enough to overcome the reduced ability of MR1 to bind B2M in D6. Alternatively, it could point towards other, as-yet undefined chaperones required for presentation of exogenous antigens on MR1. Of note, B2M itself can act as a folding chaperone for MHC class Ia proteins to enable ER egress without necessarily accompanying the antigen presenting molecule through the anterograde pathway [56, 57]. It is, thus, possible that even with a lower affinity for B2M, MR1 molecules in the D6 cell line could form transient interactions with B2M in the ER, which may allow them to leave this compartment. Another possible explanation for the maintenance of the endosomal antigen presentation pathway when B2M is limiting is that sampling of the endosomal compartment may simply be more efficient and, thus, require fewer MR1:B2M dimers to achieve MAIT cell activation. This would explain why MR1-mediated endosomal antigen presentation in the D6 cell line is functional even though MR1-GFP levels in vesicles are below the limit of detection.

We hypothesize that at baseline, only a small proportion of the MR1 in a cell is funneled into the endosomal compartment whereas the majority remains in the ER, available for loading with exogenous ligands as characterized extensively [13, 18–21, 28, 58]. These endosomal MR1 molecules would then be able to sample antigens derived from intracellular infections. This notion is not without precedent as different subcellular pools of MHC class Ia molecules are employed for ER-based vs cross-presentation of peptides and trafficking of the antigen presenting molecules depends on the nature and source of the antigen [59–61]. For MR1, it is unclear whether those molecules reach the endosomal compartment via the plasma membrane or whether there is direct trafficking from the ER to the endosomal compartment as has been suggested in the context of MHC class Ia cross-presentation [59, 60]. Irrespective of the route of delivery, the data presented here suggest that increasing the flux of MR1 molecules along this pathway suffices to enable presentation of endosomal antigens even in conditions where B2M is limiting. Our data also confirm, however, that this pathway still requires B2M. Thus, MR1 and B2M must both be available in the compartment where loading of endosomal antigen occurs – whether they arrived there together or separately.

In addition to their mode of delivery, another important difference between the synthetic MR1 ligands presented by the D6 mutant cell line (6-FP, high concentrations of 5-OP-RU, and 5-A-RU prodrug) or not (DZ, PLI) is their ability to form a Schiff base with the K43 residue of MR1 as the former covalently bind to MR1 while the latter do not [7, 8, 49, 62]. For MHC class Ia, B2M and ligand binding are cooperative with initial B2M association increasing the affinity of the MHC class Ia heavy chain for peptide ligand and subsequent peptide binding enhancing B2M association, thus stabilizing the heterotrimeric complex [63–65]. Artificially enhancing the association between B2M and MHC class Ia heavy chains consequently improves the thermal stability of MHC class Ia complexes with low-affinity peptides but interestingly does not further increase the stability of complexes with high-affinity ligands [66]. Postulating that B2M and ligand binding in the context of MR1 complex formation and stability are similarly allosterically coupled, we hypothesize that binding of Schiff base-forming ligands such as 6-FP or 5-OP-RU, whether delivered as free 5-OP-RU or derived from the 5-A-RU prodrug, may allow the stable association of the D6ins MR1 mutant with B2M despite its lower affinity for the light chain. Binding of non-covalent, low-affinity lumazine ligands DZ and PLI, on the other hand, may not suffice to support stable complex formation and/or maintenance. Intriguingly, the data would consequently suggest that M. smeg supernatant contains mainly non-Schiff base-forming lumazine ligands whereas antigens derived from intracellular infection with Mtb include covalently binding pyrimidine ligands. Of note, the covalent ligand 6-FP was present throughout the immunoprecipitation experiments which identified the lower B2M association. This indicates that Schiff base formation alone does not completely rescue B2M association for the D6ins and D6del MR1 mutants. Similarly, MR1 in the D8 mutant cell line was capable of presenting endosomal antigen, but undetectable at the cell surface after incubation with 6-FP, further pointing to molecular determinants beyond Schiff base formation. An endosomal chaperone could, for example, enhance the affinity of MR1 for ligands in this compartment and contribute to the cooperative folding in this way. Moreover, since pyrimidine MR1 ligands are highly unstable under physiological conditions [62, 67], they may be more efficiently able to support folding of MR1 in the D6 cell line when released from bacteria directly in endosomal compartments, where MR1 is available to immediately capture the ligands and prevent cyclization into lumazines. Thus, both antigen affinity and route of delivery likely play a role in the differential antigen presentation capacity of the MR1 mutants described here.

## Experimental Procedures

### Human subjects

This study was conducted according to the principles expressed in the Declaration of Helsinki. Study participants, protocols, and consent forms were approved by the Institutional Review Board at Oregon Health & Science University (IRB00000186). All ethical regulations relevant to human research participants were followed. Peripheral blood mononuclear cells (PBMC) and human serum from human subjects were only used to expand T cell clones and in ELISpot medium as described in the methods section. All findings in this manuscript are based on T cell clone function or on other cell line function, e.g. BEAS-2B cell line. No data is presented on the PBMC from human subject participants and sex and gender were not taken into consideration.

### Cells and bacteria

BEAS-2B cells were obtained from the American Type Culture Collection (ATCC) and maintained in Dulbecco’s Modified Eagle Medium (DMEM; Gibco or Corning) supplemented with 10% Fetal Bovine Serum (FBS, Gemini) and 2% L-Glutamine (Gibco). BEAS-2B MR1 KO cells and their reconstitution with a lentiviral construct encoding tetracycline-inducible, GFP-tagged MR1A (tet MR1-GFP) were described previously [25]. BEAS-2B MR1 KO tet MR1-GFP clonal cell lines were established by plating the polyclonal parent cell line in the inner 60 wells of a flat-bottom 96-well plate in conditioned medium at 0.3 cells/well. Plates were incubated at 37°C for about two weeks. Cells from wells containing a single cluster were expanded stepwise into 48-well plates, 12-well plates, and 6-well plates. Clonal cell lines were screened for GFP expression by flow cytometry after addition of dox (Sigma) and interesting clones were selected for further characterization. MR1-restricted T cell clones D481-C7 and D426-G11 were characterized previously [10, 31] and expanded as described in [25]. The BEAS-2B cell line constitutively over-expressing MR1-GFP under a minimal CMV promoter and the BEAS-2B cell line over-expressing tet-inducible MR1-GFP in addition to endogenous MR1 have been published before [49, 50, 68]. The Mtb auxotroph strain MC^2^ 6206 is deficient in the synthesis of leucine and pantothenate (H37Rv ΔpanCD ΔleuCD) [69] and was grown in Middlebrook 7H9 broth supplemented with Middlebrook ADC (ThermoFisher Scientific), 0.5% glycerol (Fisher), 0.05-0.1% Tween-80 (EMD Chemicals), 50 μg/ml leucine (Sigma-Aldrich), and 24 μg/ml pantothenate (Sigma-Aldrich). Fresh cultures were started from frozen stocks for each experiment and grown to an OD600 between 0.3 and 1.1. M. smegmatis MC^2^ 155 was grown in 7H9 broth without leucine and pantothenate. BCG was a gift from Peter Sander and grown in Middlebrook 7H9 broth supplemented with Middlebrook ADC (ThermoFisher Scientific) and 0.2% glycerol before freezing aliquots at-80°C. BCG was used directly from frozen stocks after passing the thawed cell suspension through a 27G needle. The titer of the stock was 1.34e9 colony forming units (CFU)/ml.

### Flow Cytometry

To determine surface expression of MR1 and HLA-Ia, BEAS-2B MR1 KO tet MR1-GFP clonal cell lines were incubated with the indicated concentrations of dox for at least 5h before addition of 100 μM of 6-FP (Schirks Laboratories) or the equivalent volume of 0.01 M NaOH (Sigma) overnight. Cells were harvested and washed in flow buffer (PBS + 5% goat serum + 5% human serum + 0.5% FBS) and then incubated in 25 μl of antibody mix containing APC-conjugated anti-MR1 antibody (clone 26.5; BioLegend, #361108) and, if indicated, PE-conjugated anti-HLA-Ia antibody (clone W6/32; BioLegend, #311406) at 4°C for at least 30 min. Cells were then washed in flow buffer, washed in PBS, and ultimately fixed in 1% paraformaldehyde (Electron Microscopy Sciences) for at least 20 min at room temperature (RT) or overnight at 4°C. BEAS-2B WT cells were used for the unstained controls. For the APC single color controls, WT BEAS-2B cells were stained with W6/32-APC (BioLegend, #311410). For the GFP time course experiment, BEAS-2B MR1 KO tet MR1-GFP clonal cell lines were incubated with 2 μg/ml of dox overnight. Medium containing dox was washed off at the indicated timepoints and replaced with medium without dox. Cells were harvested, washed in PBS and fixed as above. To confirm comparable MR1-GFP expression in assays with adjusted dox concentrations, induced cells were washed with PBS and fixed with 1% paraformaldehyde (4% for Aux Mtb experiments) at the time of addition to the ELISpot plate or cell lysis before analyzing GFP expression by flow cytometry. For the BFA decay assay, BEAS-2B MR1 KO tet MR1-GFP clonal cell lines D4 and D6 were incubated with 4 or 8 μg/ml dox, respectively, overnight. On the next day, dox was refreshed and either 100 μM of 6-FP or the same volume of 0.01 M NaOH was added overnight. At the indicated time points, medium containing dox and 6-FP was washed off and replaced with medium containing dox and 10 μg/ml of BFA. Cells were stained for MR1 surface expression as above. To determine the concentrations of dox that resulted in comparable expression levels of MR1-GFP, BEAS-2B MR1 KO tet MR1-GFP clonal cell lines were incubated with 0, 1, 2, 4, 8, or 16 μg/ml of dox overnight before harvesting and fixing. Samples were acquired on a BD LSR II cytometer at the OHSU Flow Cytometry Shared Resource (RRID: SCR_009974) and data were analyzed with FlowJo version 10.7.1 or 10.8.1 (Treestar, Ashland, OR, USA). Cells were gated based on forward scatter (FSC) area and side scatter (SSC) area, followed by a Single Cell gate based on FSC area versus FSC height and a GFP^+^ gate where appropriate. Rainbow calibration beads (Spherotech) were included in each flow cytometry acquisition to ensure comparable laser settings.

### Western blots

For analysis of whole cell lysates, equal cell numbers were lysed in lysis buffer containing 0.5% NP-40 (Chromotek) + protease inhibitor cocktail (Roche) on ice for at least 1h. Supernatants were collected after centrifugation and whole cell lysates were combined with loading buffer and reducing agent (both Invitrogen) before loading onto a 4-20% polyacrylamide gel (Bio-Rad). Gels were run at 120V for about 1h and transferred onto a polyvinylidene fluoride membrane (Millipore), which was then blocked in Odyssey blocking buffer (Li-Cor) for at least 1h at RT. When indicated, membranes were cut horizontally to blot for multiple proteins with primary antibodies of the same host species in the same samples. Membranes were incubated with primary antibodies against MR1 (Proteintech; polyclonal rabbit antibody #13260-1-AP), B2M (Abcam; monoclonal rabbit antibody #ab75853), CANX (Stressgen; polyclonal rabbit antibody #SPA-865), SQSTM1/p62 (Abcam; monoclonal mouse antibody #ab56416) and loading controls vinculin (Bio-Rad; monoclonal mouse antibody #MCA465GA) or beta-actin (Abcam; polyclonal rabbit antibody #ab8227) on a shaker at 4°C overnight. Membranes were washed repeatedly in PBS + 0.1% Tween-20 (Affymetrics) and incubated with secondary antibodies against mouse (Li-Cor; #925-68072 or #926-68070) and rabbit (Li-Cor; #926-32213 or #926-32211) conjugated to different IRDyes. WBs were imaged on a Li-Cor Odyssey imaging system. Band intensity was quantified using the Analysis functionality of the Image Studio version 5.2 software with the median local background subtracted.

### Next Generation Sequencing

For sequencing of tet MR1A-GFP lentiviral insertions in the clonal cell lines, genomic DNA was extracted from each of the four clonal BEAS-2B MR1 KO tet MR1-GFP cell lines and a clonal cell line expressing the same construct in the BEAS-2B WT background as an unedited control in the absence of the CRISPR/Cas9 system. The region around the sgRNA target site was amplified with a forward primer that binds in the CMV promoter region to avoid amplification of the endogenous locus. Both primers carried overhangs with Illumina adapter sequences (forward: 5’-TCGTCGGCAGCGTCAGATGTGTATAAGAGACAGCACCGGTGGAATTCATGGGGGAA-3’; reverse: 5’-GTCTCGTGGGCTCGGAGATGTGTATAAGAGACAGGAGGTTCTCTGCCATCCATGG-3’). Unique combinations of Nextera XT Index Primers (Illumina) were attached by Index PCR and all five samples pooled at equimolar ratios. The resulting library was sequenced paired-end using the Nano 500 MiSeq Kit (Illumina). Fastq files were then analyzed using the CRISPResso 2.0 web implementation [29]. Sequences were translated using the Expasy Translate Tool (https://web.expasy.org/translate/).

### Enzyme-linked Immunosorbent spot (ELISpot) assays

96-well mixed cellulose esters ELISpot plates (Millipore) were coated with anti–IFNγ antibody (Mabtech; #1-D1K) overnight at 4°C or for 4h at 37°C. After washing with PBS, plates were blocked with Roswell Park Memorial Institute 1640 (RPMI; Gibco or Corning) + 10% human serum + 2% L-Glutamine + 0.1% gentamycin (Gibco) for at least 1h at RT and either used directly or stored at 4°C. BEAS-2B MR1 KO tet MR1-GFP clonal cell lines were seeded in T25 flasks and incubated overnight with either 2 μg/ml or adjusted concentrations (see Figure S3) of dox as indicated in the figure legends. Cells were harvested and added at 1e4 APCs per well in RPMI + human serum containing the appropriate amount of dox to maintain MR1 expression. For M. smeg supernatant, deazalumazine (synthesized in house as previously described [32]), 5-AR-U prodrug (MedChemExpress), free 5-OP-RU (generated from 5-A-RU*HCl as described in [50]), photolumazine I (PLI, synthesized in house as previously described [10]), phytohemagglutinin (PHA, Sigma or Roche) and live M. smeg, titrations of the antigen were added directly to the ELISpot plate and incubated with the APCs for at least 1h at 37°C before addition of the T cell clones. D481-C7 T cells were used at 2.5e3 cells/well whereas D426-G11 T cells were used at 1e3 cells/well except for deazalumazine experiments where 2.5e3 cells/well were used for both clones. Cells were co-cultured for at least 18h before developing the ELISpot. For this, the plate was washed in PBS + 0.05% Tween-20 (Affymetrix) and then incubated with alkaline phosphatase-coupled secondary antibody (Mabtech; #7-B6-1-ALP) for 2h at RT before washing and addition of the BCIP/NBT substrate (Mabtech). Substrate was washed off and plates were dried before enumerating IFNγ spot forming units (SFU) on an AID ELISpot reader. For in-plate infections with M. smeg, bacteria were thawed and cultured in Difco Middlebrook 7H9 Broth (BD) supplemented with ADC Enrichment (BD) overnight before use and added based on OD600. M. smeg supernatant was generated by growing M. smeg strain MC^2^ 155 in Difco Middlebrook 7H9 Broth (BD) supplemented with ADC Enrichment (BD) overnight at 37°C, shaking. The culture was centrifuged at 3000 rpm for 20 min at 4°C and the supernatant was filtered through a 0.22 μm filter before aliquoting and storing at-80°C. Volumes for titrations M. smeg supernatant were empirically determined in functional assays for each batch. For infections with Aux Mtb and BCG, MR1 expression in BEAS-2B MR1 KO tet MR1-GFP clonal cell lines was induced with dox overnight, then cells were harvested, counted and allowed to adhere for at least 4h at 37°C. Aux Mtb was added based on OD600 values of an ongoing culture. BCG was thawed from frozen, dispersed by repeatedly passing through a 27G needle, and added at an MOI of 3 based on titer plates for the frozen stock. Infected cells were harvested after overnight incubation, washed, and diluted for use in ELISpots as above.

### Fluorescence Microscopy

2e5 cells were plated on 8-well #1.5 glass bottom chamber slides (Nunc), incubated with dox (D4: 1 ug/ml, D6: 2 ug/ml) at 37°C and left to adhere for at least 4h before addition of CellLight BacMam 2.0 (Invitrogen) for late endosomes (Rab7-RFP, C10589). The following day, cells were stained with NucBlue Live ReadyProbes (Invitrogen) and imaged in an unbiased manner by their RFP expression. Images were acquired on a motorized Nikon TiE stand with a Yokogawa W1 spinning disk unit and equipped with the following high-powered Agilent lasers and emission filters: 405nm-445/50nm, 488nm-525/36nm, 561nm-617/73nm. A 100x (numerical aperture 1.49) objective was used and images were captured by Andor Zyla 5.5 sCMOS camera with 2by2 camera bin. Co-localization was analyzed using the “Spots” function and “spots colocalization” MatLab Xtension module on Imaris (Bitplane) with spot thresholds (“Quality” values) set as appropriate for each image.

### Co-Immunoprecipitations (Co-IPs)

For the Co-IPs for mass spectrometry, MR1-GFP expression was induced in BEAS-2B MR1 KO tet MR1-GFP clonal cell lines D4 and D6 by incubation with 10 μg/ml dox overnight. On the next day, dox was increased to 20 μg/ml and cells were incubated with 100 μM of 6-FP or the equivalent volume of 0.01 M NaOH overnight. Cells were harvested and 7-9e6 cells per cell line were lysed in 400 μl of wash buffer (Chromotek) + 1% digitonin (Sigma) + protease inhibitor cocktail (Roche) on ice for at least 1h. Lysates were passed through a 27G needle 10 times each and centrifuged. The supernatant was diluted with 650 μl of wash buffer and distributed between two reactions of 50 μl of GFPTrap magnetic agarose (Chromotek) per cell line, which were rotated at 4°C for 1h. Samples were washed using a DynaMag magnetic rack (Invitrogen) and resuspended in 1 ml of PBS final volume to achieve comparable volumes between unbound samples and IP samples for the WB analysis. After collecting the aliquots for WB, samples were concentrated for mass spectrometry analysis. For the validation WBs, 20 μl of the remaining lysates from the three in-gel digest experiments were analyzed for MR1, SQSTM1/p62, and B2M (WB 1) or CANX and B2M (WB2) as above.

### Mass spectrometry – on-bead digestion method

Samples were resuspended in 100 μl of PBS and sent for on-bead digestion. Tryptic digests were conducted using Thermo Scientific’s Pierce In-Solution Tryptic Digestion and Guanidation Kit (Cat # 89895) directly on magnetic beads at 5x volume. Sample cleanup was completed using either Thermo Scientific’s Pierce C18 Spin Columns (Cat #89870), or 10 μl Spin Tips (Cat #87782). Samples were dried via speedvac and resuspended in 6 μl of a solvent composed of 10% acetic acid, 2% acetonitrile, and indexed Retention Time (iRT) peptides (Biognosys, Schlieren, Switzerland) at a 1:50 dilution in HPLC grade water.

Samples were then injected using an Eksigent NanoLC (nanoscale liquid chromatography) 415 reversed-phase HPLC. A trap-elution method was used with a 5-mm-long, 350-μm-internal diameter ChromXP C18 trap column by Optimize Technologies with 3-μm particles and 120-Å pores. After desalting, samples were then loaded onto a 150-mm-long, 75-μm-internal diameter ChromXP Eksigent C18 reversed-phase separation column with 3-μm particles and 120-Å pores. An acetonitrile gradient was run at pH 3.0 using two solvents. Solvent A was 0.1% formic acid in HPLC-grade water, and solvent B was 0.1% formic acid in 95% acetonitrile in HPLC-grade water. After column equilibration with 2% solvent B, samples were loaded at 5 μl over 10 min onto the trap column and injected through the separation column at 300 nl/min with a gradient of 10–40% B for 70 min, followed by a clearance gradient of 40– 80% B for 7 min.

Sample passing through the separation column was ionized using the PicoView ion source by New Objective, through an 11-cm, 20-μm-inner diameter pulled glass emitter that tapered to 10 μm at the tip. The ionized sample was injected into an AB Sciex TripleTOF 5600 quadrupole time of flight (TOF) mass spectrometer in positive ion mode with parameters set at the manufacturer’s default tryptic digest settings.

Peptide sequences were determined using PEAKS Studio X+ software (Bioinformatics Solutions, Waterloo, Canada). A database composed of SwissProt Homo sapiens (Proteome ID UP000005640, Taxon ID 9606), MR1-GFP construct sequence, and iRT peptide sequences was used as the reference for database search. The following variable post-translational modifications were included in the database search: acetylation, deamination, pyroglutamate formation, oxidation, sodium adducts, phosphorylation, cysteinylation, and carbamidomethylation. Identified peptides were further refined using a false discovery rate of 1%, requiring at least two unique, significant peptides for protein identification. Peptide intensity values for each protein ID were normalized by dividing by the corresponding protein’s molecular weight, adjusting for protein size. These were then normalized to MR1-GFP in each sample and averaged from three experiments. Indexed Retention Time peptides and immunoglobulin peptides (HV302, HV319, HV312, HV305, HV304) were removed from the analysis. The mass spectrometry proteomics data have been deposited to the ProteomeXchange [70] Consortium via the PRIDE [71] partner repository with the dataset identifier PXD063725. The list of proteins and peptides obtained during the analysis are in Table S4.

### Mass spectrometry – in-gel digestion method

Agarose-bound proteins were boiled in NuPAGE LDS loading buffer (4X) mixed with 50 mM DTT (both Invitrogen) before briefly running on a 4-20% polyacrylamide gel (Bio-Rad). Gels were stained with Coomassie, and the gel sections containing the proteins were cut into small pieces for in-gel digestion. Briefly, the gel pieces were destained using 60% acetonitrile in 0.2 M ammonium bicarbonate, then dried. The dried gel pieces were mixed with 18 µL of trypsin (at a concentration of 0.083 µg/µL in 0.2 M ammonium bicarbonate) and incubated for 17 hours at 37°C. Extracted peptides were diluted in 3% acetonitrile and 0.1% formic acid in water. All reagents used were of LC/MS grade. The experiment was conducted on three different days, with four injections on day 1 and three injections on days 2 and 3 for each sample.

### LC-MS/MS analytical conditions for Data independent Acquisition (DIA) Proteomics

Peptide concentration was quantified using a nanodrop, and 0.25 µg of peptides per sample were loaded onto a trap cartridge (PepMap™ Neo 5 μm C18 300 μm X 5 mm, Thermo FisherScientific) and separated on an analytical column (10 cm x 75 µm, C18, 1.9 µm, BrukerDaltonics). The gradient was generated using nanoElute liquid chromatography (LC) system (Bruker Daltonics, Bremen, Germany) with solvent A (0.1% formic acid in water) and solvent B (0.1% formic acid in acetonitrile). The gradient was 5%–30% of solvent B in 17.8 min, 30%–95% solvent B in 0.5 min, maintaining at 95% solvent B for 2.4 min at a flow rate of 500 nL.min−1. The LC system was connected to a TIMS TOF Flex mass spectrometer (BrukerDaltonics) via a CaptiveSpray nano-electrospray source, with a capillary voltage of 1600 V, dry gas at 3.0 L/min, and dry temperature at 200 °C. According to the instrument specifications, the MS has a full sensitivity mass resolution of 60,000 at 1,222 m/z. For the spectral library generation, samples were acquired in Data Dependent Acquisition (DDA)–Parallel accumulation Serial Fragmentation (PASEF) mode. Ramp time was set to 75 ms, resulting in an accumulation time of 75 ms and total cycle time of 0.57s. DDA analysis with 6 PASEF scans covered a mass range from 100 m/z to 1700 m/z.

For the quantitative analysis, data independent acquisition (DIA)-PASEF mode was used, with a windows scheme of 7 MS/MS ramps with 19 mass steps per cycle, and a cycle time of 0.65s. Mass overlap and mass width were set to 1.0 Da and of 50 Da respectively, covering a mass-over-charge (m/z) range from 307.8 to 1,239.8 (Figure S14). For both DDA and DIA, a linear collision energy ramp was used for peptide fragmentation: from 20-59, with an ion mobility range from 0.6 to 1.6 Vs/cm2. For all experiments, we calibrated both m/z and ion mobility dimension linearly using three compounds of the following m/z and ion mobility (m/z, 1/K0) values: 622.029, 0.99915 Vs/cm2; 922.0098, 1.1986 Vs/cm2; and 1221.9906, 1.3934 Vs/cm2.

### LC-MS/MS Data and Statistical Analysis

The raw data files were analyzed using FragPipe version 22.0 that contained the database search tool MSFragger version 4.1 for peptide identification [72, 73], diaTracer [74] version 1.1.5 for spectrum deconvolution of our DIA-PASEF data, MSBooster version 1.2.31 for improving peptide identification rates using deep learning predictions [75], EasyPQP version 0.1.50 which is a phyton package for spectral library generation (https://github.com/grosenberger/easypqp), and the built-in DIANN version 1.8.2 Beta 8 [76] for protein quantification, among other embedded validation tools [77–79]. Peptide sequences were searched against the Homo sapiens proteome (UP000005640) retrieved from UniProt on 06/18/2024. Decoy and contaminant protein sequences were added to the human proteome FASTA file. For the MSFragger search, precursor tolerance was set to (-20) to 20 ppm and fragment tolerance was set to 20 ppm, with mass calibration and parameter optimization enabled. Isotope error was set to 0/1/2. One missed cleavage was allowed with trypsin digestion, along with the following variable modifications: oxidation of methionine (+15.9949) and acetylation of protein N-terminal (+42.0106). Additional settings included the top 150 most intense peaks and a minimum of 15 fragment peaks to search for a spectrum. Peptide spectral match (PSM) validation, protein interference, and false discovery rate (FDR) filtering were performed using Percolator, ProteinProphet, and Philosopher, respectively [77–80], to retain only the highest-scoring peptide matches for protein identification validation. A spectral library was generated using DDA data from the same samples run on the same instrument and using the same LC separation method. The default EasyPQP setup was used for spectral library generation, allowing a maximum unimod mass difference of 0.02 units and a maximum mass discrepancy of 15 PPM. Only b and y fragment ions were selected for peptide identification, with no retention time correction applied. The peptide ions in the spectral library were filtered with a 1% global peptide and protein FDR. The spectral library was then used to extract and quantify precursors, peptides, and proteins from the DIA data using DIA-NN.

Statistical analysis was performed using FragPipe-Analyst that has an in-house generated R script based on the ProteinGroup.txt file of DIA-NN [81]. First, contaminant proteins and reverse sequences were filtered out. In addition, proteins that have been only identified by a single peptide and proteins not identified/quantified in at least 66% of the replicates in one condition have been removed as well. The MaxLFQ data was converted to log2 scale, samples were grouped by conditions and missing values were imputed using man method, which uses random draws from a left-shifted Gaussian distribution of 1.8 StDev (standard deviation) apart with a width of 0.3. Protein-wise linear models combined with empirical Bayes statistics were used for the differential expression analyses. The limma package from R Bioconductor was used to generate a list of differentially expressed proteins for each pair-wise comparison. A cutoff of the adjusted p-value of 0.05 (Benjamini-Hochberg method) along with a |log2 fold change| of 0.9 has been applied to determine significantly regulated proteins in each pairwise comparison.

Additionally, we conducted a statistical analysis using linear models in R to assess whether batch (day of analysis) significantly impacted the abundance of proteins. We found that only 10 out of 1,472 identified proteins had a significant batch effect, suggesting that the observed differences in their abundances are likely due to technical variation associated with the different analysis days rather than the experimental conditions alone (Table S10). None of the remaining proteins, including the seven reported as significantly different, had a significant batch effect. The mass spectrometry proteomics data have been deposited to the ProteomeXchange [70] Consortium via the PRIDE [71] partner repository with the dataset identifier PXD062277. The list of proteins and peptides obtained during the analysis are in Table S5.

### Overexpression of B2M and individual MR1 mutants

To overexpress B2M in the D6 clonal cell line, the B2M coding sequence (NCBI entry NM_004048.4) with a downstream internal ribosomal entry site (IRES) GFP [82] was ordered from Twist Biosciences and cloned into the pCI AscI mammalian expression vector previously described [21]. The sequence was confirmed by Sanger sequencing at the OHSU Sequencing Core. The plasmid was transfected into BEAS-2B MR1 KO tet MR1-GFP clonal cell lines D4 and D6 using an Amaxa Nucleofector with Nucleofection Kit T (Lonza) according to the manufacturer’s instructions. Cells were left to rest overnight and then induced with 0.5 (D4) or 1 (D6) μg/ml dox overnight before being used in an ELISpot as described above. B2M protein expression at the time of ELISpot was assessed by WB.To express individual MR1 mutants D4, D6ins, and D6del, the tet MR1-GFP insertions in the D4 and D6 parent cell lines were amplified with primers that covered the 5’ EcoRI restriction site included in the lentiviral insertion and a 3’ PvuII site within the MR1 coding region (forward primer: TCGTCGGCAGCGTCAGATGTGTATAAGAGACAGCACCGGTGGAATTCATGGGGGAA; reverse primer: AGCCCCTCAGCAGCTGAGTGTA). For the V12S mutation, two fragments were amplified, and the point mutation was introduced via overlapping primers (outer primers as above; internal reverse primer: ATGGGATCCGAACTGCCCAGGCGAAA; internal forward primer: TTTCGCCTGGGCAGTTCGGATCCCAT).

Amplicons were annealed and re-amplified. PCR products were then cloned into our pCI AscI MR1 res expression vector which features an IRES GFP (Addgene #214752) [50] and transformed into NEBalpha competent bacteria (NEB). Colonies were screened for plasmids with the correct insertions by Miniprep (QIAGEN) and Sanger sequencing at the OHSU Sequencing Core. The plasmids were nucleofected into BEAS-2B MR1 KO cells to express the MR1 mutants in the absence of endogenous MR1.

### Knock-down of candidate gene expression

For siRNA-mediated knock-down, cells were transfected with 30 pmol of siRNA targeting B2M (Thermo Fisher Scientific, Assay ID s1853) or SQSTM1/p62 (Thermo Fisher Scientific, Assay ID 137334) or missense Silencer Select Control No 1 (Thermo Fisher Scientific, 4390844) using Nucleofector Kit T (Lonza) and an Amaxa Nucleofector according to the manufacturer’s instructions. For assays with exogenous antigens, cells were left to recover and adhere at 37°C for 4-5 h in SQSTM1/p62 KD experiments or overnight for B2M KD experiments before addition of dox overnight if applicable. Cells were used in an ELISpot as above on the next day. For Aux Mtb infections with SQSTM1/p62 KD, cells were induced to express MR1-GFP overnight first. The next day, cells were harvested, nucleofected with the siRNA and allowed to recover for at least 3h. Cells were then re-counted, re-plated, and infected overnight. For Aux Mtb infections with B2M KD, cells were transfected as above, left to recover for at least 4h, and then incubated with dox overnight. They were infected on the next day after which the dox concentrations were lowered to 0.3 and 0.6 μg/ml, respectively. The ELISpot was performed in the usual 0.5 and 1 μg/ml after overnight infection.

### B2M dissociation IP

Cells were induced to express MR1-GFP with 0.5 (D4) and 8 (D6) μg/ml dox overnight to ensure presence of sufficient B2M for comparable WB analysis. On the next day, medium was replaced with medium containing both dox at the respective concentrations and 100 μM 6-FP. After overnight incubation, 4e6 cells per cell line were lysed in 200 μl of wash buffer (Chromotek) + 1% digitonin (Sigma) + protease inhibitor cocktail (Roche) on ice for at least 1h. Lysates were centrifuged and the protein-containing supernatant was diluted with 300 μl of wash buffer (Chromotek). MR1-GFP was immunoprecipitated with 50 μl of GFPTrap magnetic agarose (Chromotek) as described above. After the final wash, magnetic agarose was resuspended in 90 μl of wash buffer supplemented with protease inhibitor cocktail and split into four 21 μl aliquots. These were incubated at 37°C for the indicated amounts of time. At each time point, aliquots were placed on the magnetic rack, the unbound fraction was removed, and the magnetic agarose resuspended in the same volume of wash buffer + protease inhibitor cocktail. The bound and unbound fractions for each cell line at each time point were mixed with loading buffer and reducing agent (both Invitrogen) and boiled for 10 min. Samples were stored at - 20°C and later analyzed by WB as above.

### CRISPR/Cas9-mediated KO of B2M

6e5 cells were electroporated with ribonucleoprotein (RNP) complexes prepared from SpCas9 2NLS Nuclease and either non-targeting control scrambled sgRNA No 1) or three sgRNAs targeting B2M (all Synthego) using a Neon NxT Electroporation system (Invitrogen) according to the manufacturer’s instructions. In short, SpCas9 and sgRNA were mixed in Resuspension Buffer (Synthego) at a 1:9 molar ratio and incubated at RT for 10 min before transfecting the cells using two 20 ms pulses at 1400 V. Editing was confirmed by PCR amplification of the target region in the polyclonal cell line before proceeding. Clonal cell lines were established from the polyclonal transfected cell lines by plating 0.3 cells/well in a flat-bottom 96-well plate. Cells from wells containing a single cluster were expanded stepwise into 24-well plates, 6-well plates, and T25 tissue culture flasks. Clonal cell lines were screened by PCR amplification of the target region in genomic DNA (Figures S12 and S13). Non-targeted (NT) and B2M KO clones that had developed at comparable rates were chosen for further characterization for both D4 and D6 and B2M KO status was confirmed by WB and qRT-PCR (Figures S12 and S13).

### Quantitative reverse transcription polymerase chain reaction (qRT-PCR)

To confirm loss of B2M expression, total RNA was extracted from D4 and D6 NT and B2M KO clones using the RNeasy Plus kit (QIAGEN). RNA was converted to cDNA using the High-Capacity cDNA Reverse Transcription Kit (Life Technologies) including controls without enzyme for each sample. cDNA was purified using the PCR purification kit (QIAGEN) and analyzed using Taqman probes (Thermo Fisher Scientific) to detect transcripts of B2M (Hs00187842_m1) and ACTB (Hs01060665_g1) as an internal reference. The cycle threshold was set to 0.2 across experiments and relative expression levels compared to NT controls and normalized to ACTB were calculated (Figures S12 and S13).

### Data analysis and Statistics

Data were graphed and analyzed in Graphpad Prism version 10.4.1. Figures were assembled in Inkscape. Details of statistical analyses are provided in the Figure Legends.

## Data Availability

The mass spectrometry proteomics data have been deposited to the ProteomeXchange [70] Consortium via the PRIDE [71] partner repository with the dataset identifiers PXD062277 and PXD063725. All other relevant data are contained within the manuscript and its Supporting Information. Raw data can be obtained by contacting the corresponding author at (DML).

## Supporting Information

This article contains supporting information. [29]

## Supporting information

SupportingInformation

## Abbreviations

5-A-RU: 5-Amino-6-D-ribitylaminouracil
5-OP-RU: 5-(2-oxopropylideneamino)-6-D-ribitylaminouracil
6-FP: 6-formylpterin
A.U.: arbitrary units
aa: amino acid
ANOVA: analysis of variance
APC: antigen presenting cell
APC: allophycocyanin
ATCC: American Type Culture Collection
Aux Mtb: Mycobacterium tuberculosis auxotroph B2M β_2_-microglobulin
BCG: Bacillus Calmette–Guérin
BFA: brefeldin A
cDNA: complementary deoxyribonucleic acid
CFU: colony-forming unit
CMV: cytomegalovirus
CT: threshold cycle
DDA: data-dependent acquisition
DIA: data-independent acquisition
DMEM: Dulbecco1s modified Eagle1s medium dox doxycycline
DZ: deazalumazine
ELISpot: enzyme-linked immunosorbent spot assay ER endoplasmic reticulum
FBS: fetal bovine serum
FC: fold change
FDR: false discovery rate
GeoMFI: geometric mean fluorescence intensity HLA human leukocyte antigen
IFNγ: interferon-γ
IP: immunoprecipitation
IRES: internal ribosomal entry site
iRT: indexed retention time
KD: knock down
KO: knock out
M. smeg: Mycobacterium smegmatis
MAIT: mucosal-associated invariant T cell
MHC: major histocompatibility complex
mis: missense
MR1: MHC class I-related protein 1
MR1T: MHC class I-related protein 1-restricted T cell Mtb Mycobacterium tuberculosis
noRT: no reverse transcriptase
NT: non-target
PASEF: parallel accumulation serial fragmentation PBMC peripheral blood mononuclear cells
PE: phycoerythrin
PHA: phytohemagglutinin-L
PLI: photolumazine I
PSM: peptide spectral match
RF: reading frame
RNP: ribonucleoprotein
RPMI: Roswell Park Memorial Institute medium
RT: room temperature
SD: standard deviation
sgRNA: small guide RNA
TCR T: cell receptor
tet: tetracycline
WB: Western blot

## Acknowledgements

We thank Gwendolyn Swarbrick, Oregon Health and Science University, for critical reading of the manuscript and valuable feedback on data analysis and figure composition. We thank Prof. Sebastian Springer, Constructor University, for insightful discussions. We acknowledge expert technical assistance by staff in the Advanced Light Microscopy Core in the Department of Neurology and Jungers Center at Oregon Health and Science University, the staff at the Primate Genetics Core at Oregon National Primate Research Center, the Vollum DNA Sequencing Core at OHSU, and the assistance of the Oregon Clinical & Translational Research Institute, which is supported by the National Center for Advancing Translational Sciences, National Institutes of Health, through Grant Award Number UL1TR002369. The research reported in this publication used computational infrastructure supported by the Office of Research Infrastructure Programs, Office of the Director, of the National Institutes of Health under Award Number S10OD034224. Analytical flow cytometry was performed in the OHSU Flow Cytometry Shared Resource (RRID: SCR_009974). We would like to thank the participants who gave time and dedication to this health research as well as Erin Merrifield, Department of Pediatrics, Oregon Health and Science University, for her contributions to this study. We are grateful to the laboratory of Dr. William Jacobs, Harvard School of Public Health, for sharing the Mtb auxotroph. This work was supported in part by Merit Review #I01 BX000533 from the United States (U.S.) Department of Veterans Affairs Biomedical Laboratory Research and Development Service (DML) and NIH grants AI134790 (DML), T32HL083808 (SK), 1UC7AI180308-01 (KMD), 3G20AI167348-01 (KMD), AI147954 (WHH), and AI141549 (FGT).The content is solely the responsibility of the authors and does not necessarily represent the official views of the National Institutes of Health. The contents do not represent the views of the U.S. Department of Veterans Affairs or the United States Government.

