## SupportingInformation for "Mutations outside the MR1 antigen binding groove differentially inhibit presentation of exogenous antigens"

Figures S1 – S13

Tables S1 – S10

SI References

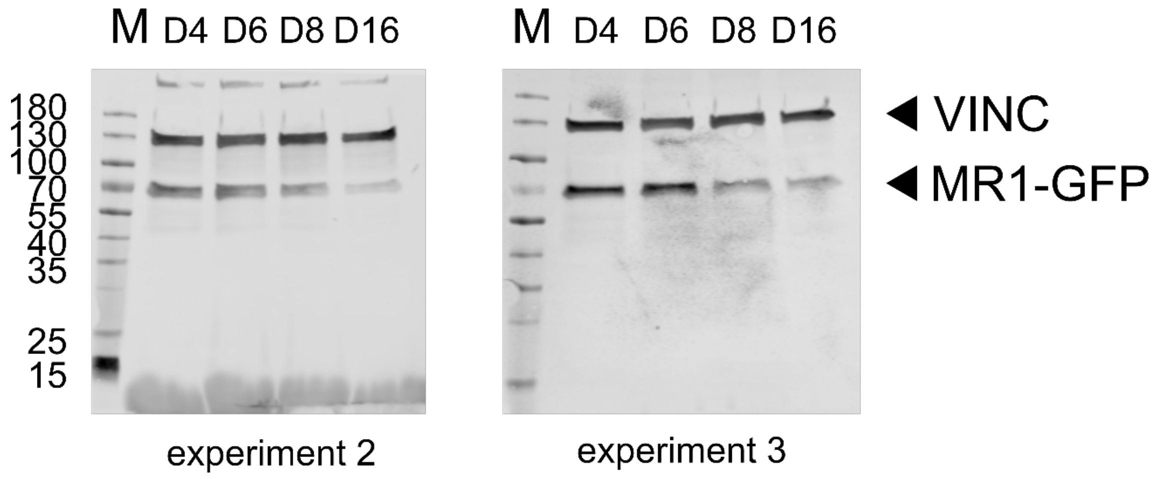

**Figure S1: Repeat experiments of the Western blot shown in Figure 1D.** WB analysis of MR1-GFP expression as in Figure 1D. Whole cell lysates from cell lines induced with 2 µg/ml doxycycline overnight were analyzed for expression of MR1-GFP and loading control Vinculin (VINC) by WB. Primary antibodies were from different species and detected in parallel with species-specific secondary antibodies conjugated to distinct IRDyes and both channels exported as greyscale. Molecular weight markers are indicated in kDa on the left.

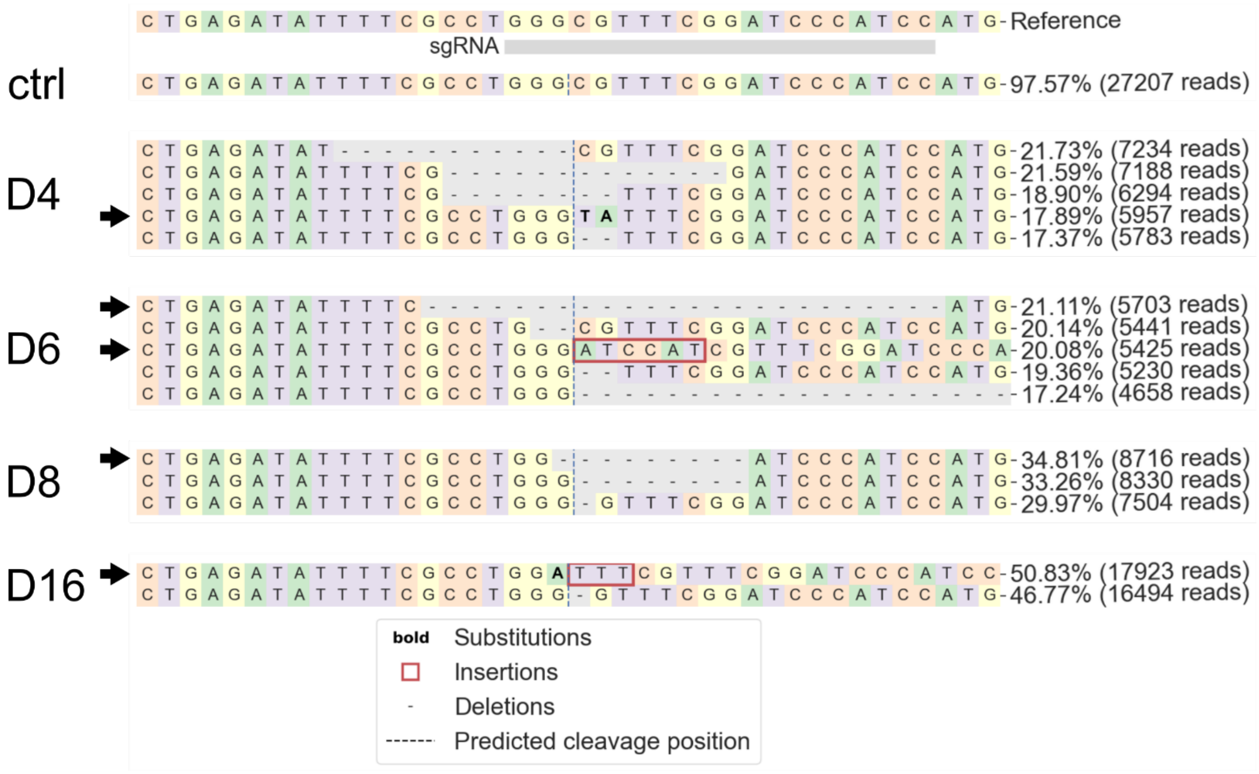

**Figure S2: Sequences of all insertions of the tet MR1-GFP cassette in each clonal cell line.** The region around the sgRNA target site was amplified by PCR and amplicons were sequenced using the MiSeq System. Data were analyzed and visualized with CRISPResso 2.0 [1]. Contributions of each sequence are shown to the right. Black arrows indicate sequences with an intact reading frame.

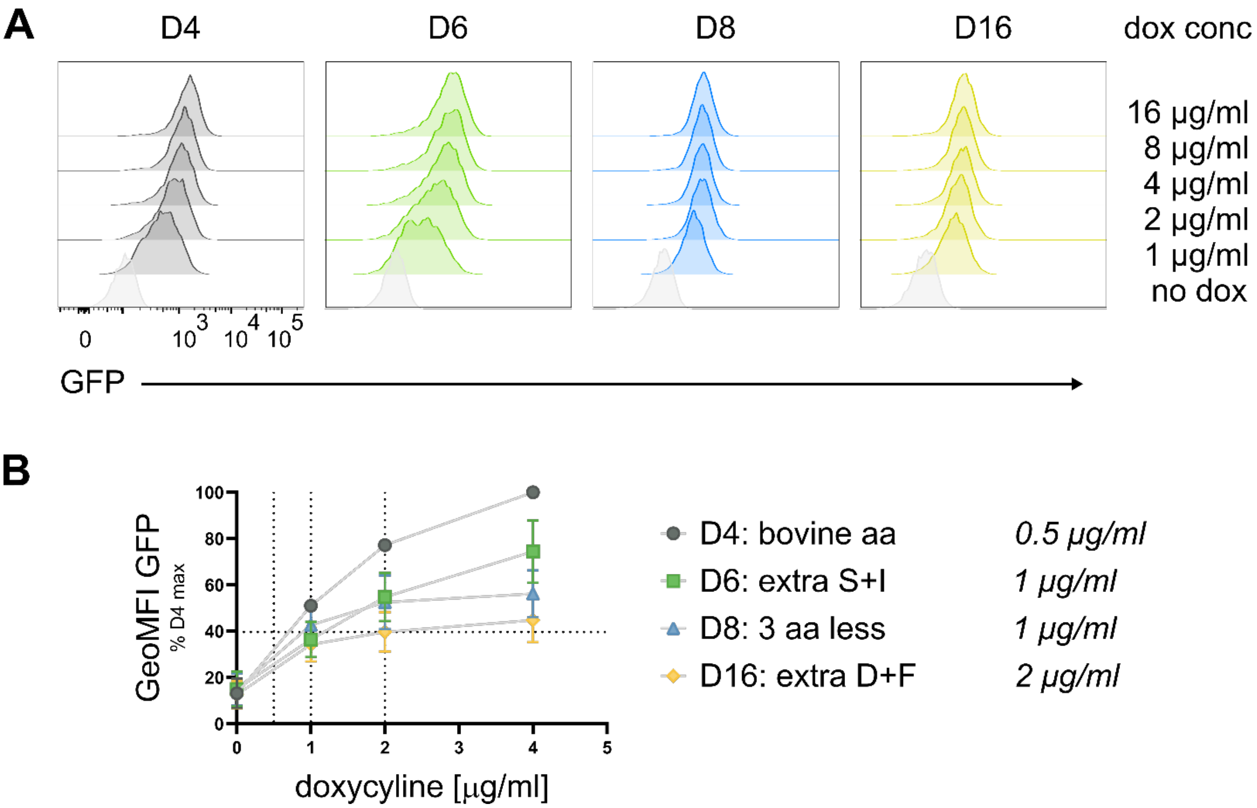

**Figure S3: Doxycyline titration to normalize MR1-GFP expression.** The four clonal cell lines were incubated with the indicated concentration of doxycycline (dox) overnight and analyzed for GFP expression to determine dox concentrations that induced comparable levels of MR1-GFP. Representative histograms are shown in *A*. Data in *B* are pooled from three independent experiments, normalized to D4 at the highest antigen concentration and shown as mean with SD. Calibration beads were included in each experiment. Based on data in *B*, the following concentrations were routinely used in functional experiments: 0.5 µg/ml for D4, 1 µg/ml for D6 and D8, 2 µg/ml for D16. aa = amino acid.

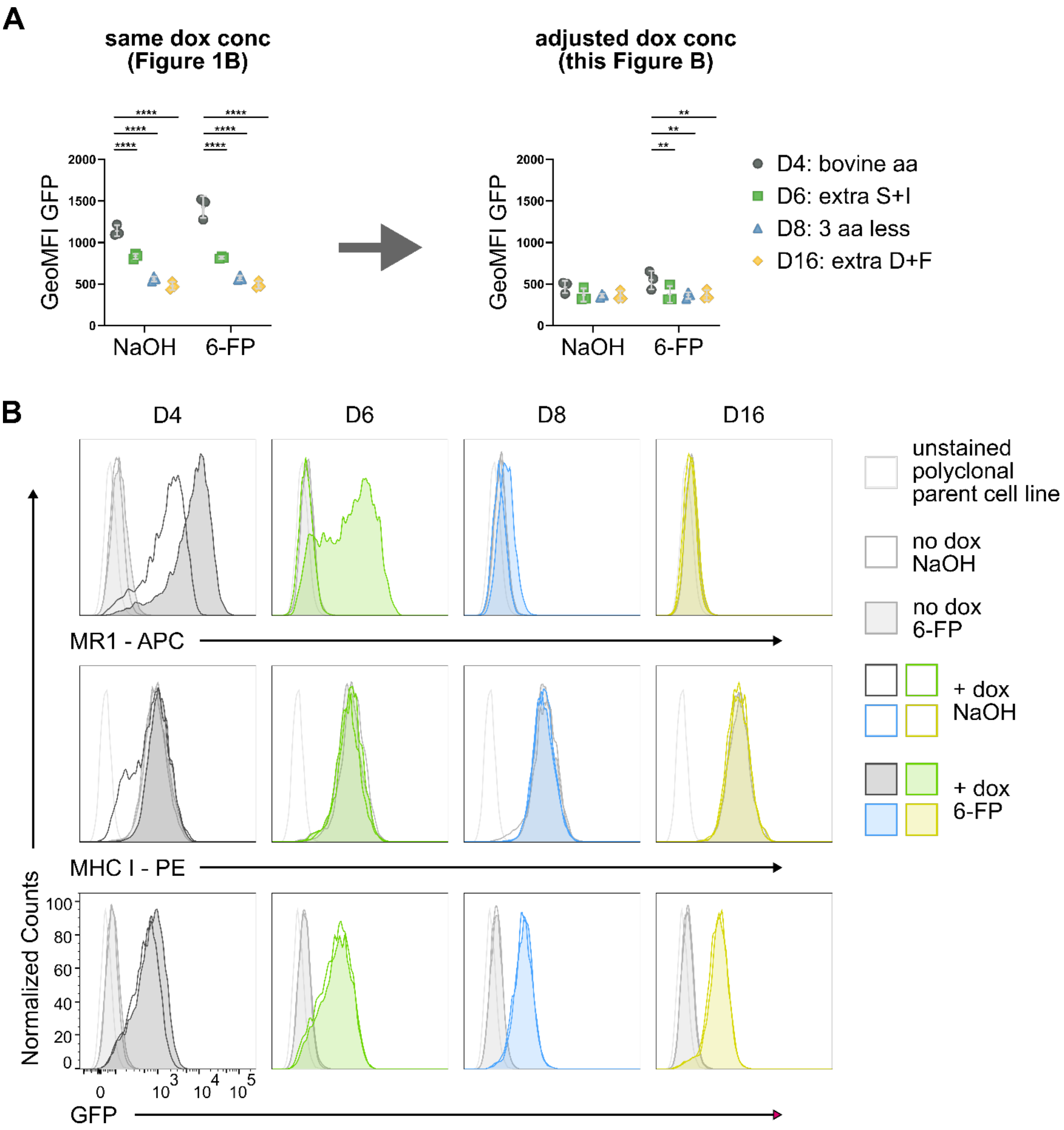

**Figure S4: Repeat of flow cytometric analysis with adjusted dox concentrations.** Experiment performed as in Figure 1B, but with dox concentrations optimized to normalize MR1-GFP expression (see Figure S3). *A.* Quantification of MR1-GFP geometric mean fluorescence intensity (GeoMFI) pooled from three experiments with 2 µg/ml dox for all four cell lines (including the representative experiment shown in Figure 1B) and three experiments with 0.5, 1, or 2 µg/ml dox (including the representative experiment shown in *B*). Experimental groups were compared by repeated-measures ANOVA with Tukey's multiple comparisons test and statistically significant differences are indicated. * = $p \leq 0.05$ ** = $p \leq 0.01$ *** = $p \leq 0.001$ **** = $p \leq 0.0001$. Comparisons of all experimental groups are shown in Table S1. *B*. Histograms representative of three independent experiments.

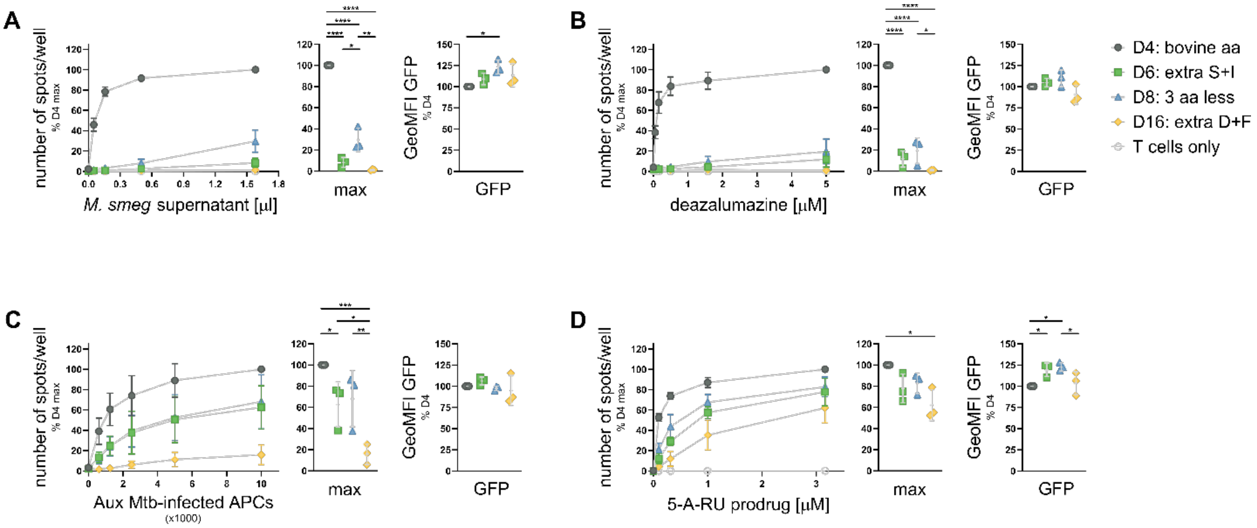

**Figure S5: Repeat of Figure 2 with MAIT cell clone D426-G11.** MR1-GFP expression was induced with adjusted dox concentrations (see Figure S3 and text) overnight and cells were used as APCs in IFNγ ELISPOTs as in Figure 2. Cells were pre-incubated with indicated dilutions of *M.smeg* supernatant (*A*), deazalumazine (*B*), or 5-A-RU prodrug (*D*) for at least one hour before addition of MAIT cell clone D426-G11. APCs in *C* were infected with Aux Mtb overnight and diluted as indicated. IFNγ responses at the highest antigen concentrations are additionally shown as dot plots. MR1-GFP expression at the time of the ELISPOT was measured and is shown to the right of each plot. Data are pooled from three independent experiments, normalized to D4 at the highest antigen concentration and shown as mean with SD. Experimental groups were compared by repeated-measures ANOVA with Tukey's multiple comparisons test and statistically significant differences are indicated. * = $p \leq 0.05$ ** = $p \leq 0.01$ *** = $p \leq 0.001$ **** = $p \leq 0.0001$. Except for *A*, flow cytometry data is the same as shown in Figure 2 as the two MAIT cell clones were assessed in the same experiments using the same APCs. aa=amino acid; GeoMFI = geometric mean fluorescence intensity.

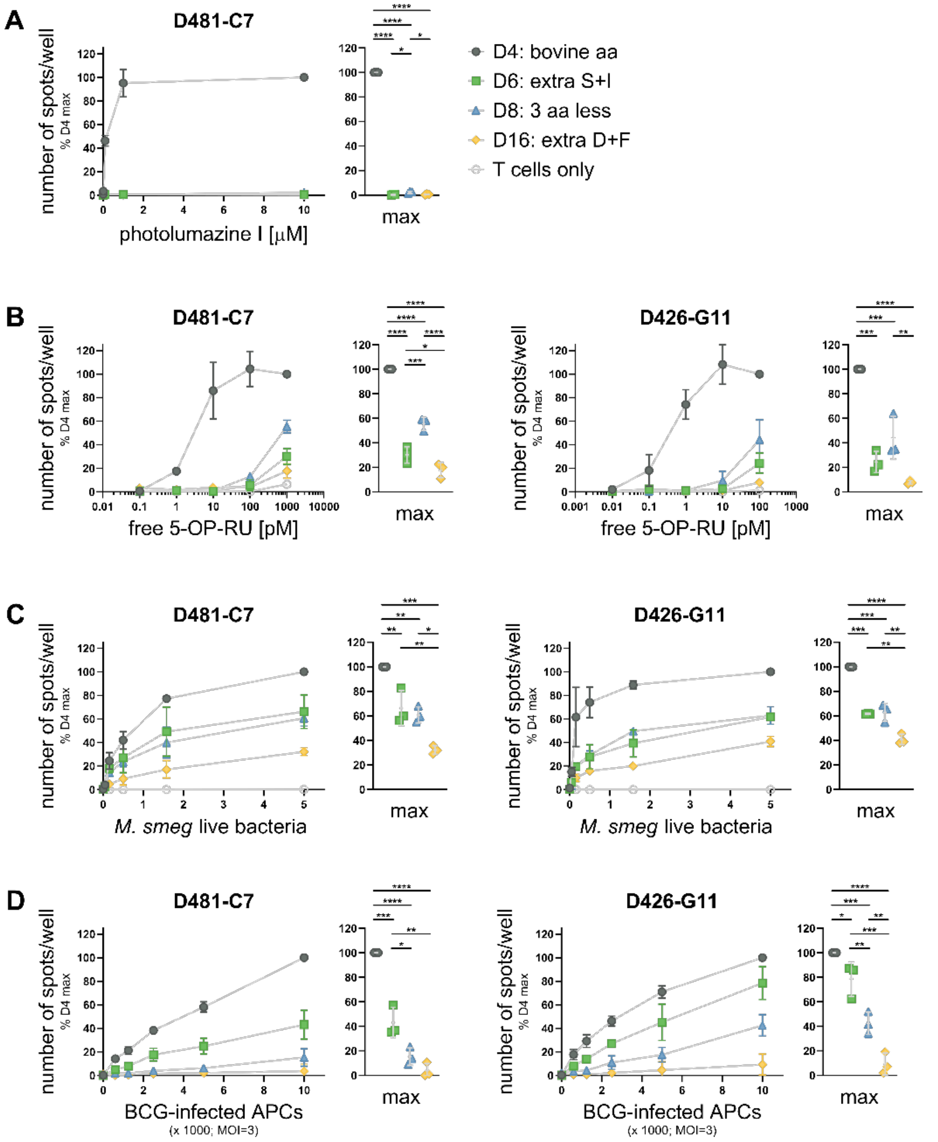

**Figure S6: Additional antigens tested with clonal cell lines induced with equal concentrations of dox.** MR1-GFP expression was induced with 2 μg/ml dox overnight and cells were used as APCs in IFNγ ELISPOTs. Cells were pre-incubated with indicated dilutions of 5-OP-RU (*A*), photolumazine I (*B*), or live *M.smeg* bacteria (*C*) for at least one hour before addition of MAIT cell clones D481-C7 and D426-G11 as indicated. APCs in *D* were infected with BCG at MOI=3 overnight and diluted as indicated. IFNγ responses at the highest antigen concentrations are additionally shown as dot plots. Data are pooled from three independent experiments, normalized to D4 at the highest antigen concentration and shown as mean with SD. Experimental groups were compared by repeated-measures ANOVA with Tukey's multiple comparisons test and statistically significant differences are indicated. * = $p \leq 0.05$ ** = $p \leq 0.01$ *** = $p \leq 0.001$ **** = $p \leq 0.0001$. aa=amino acid.

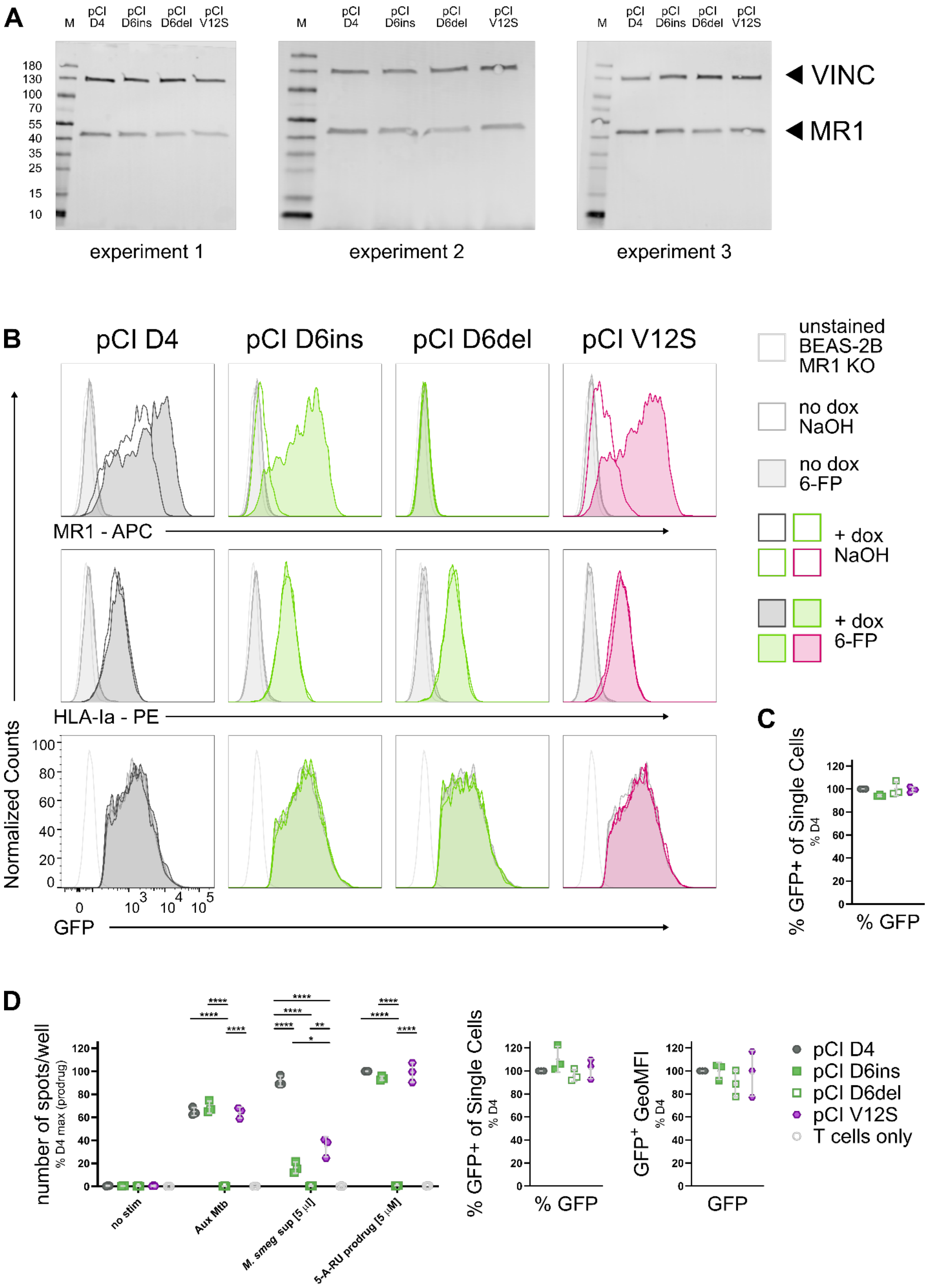
A
M
pCI D4
pCI D6ins
pCI D6del
pCI V12S
180
130
100
70
55
40
35
25
15
10
VINC
MR1
experiment 1
experiment 2
experiment 3
B
pCI D4
pCI D6ins
pCI D6del
pCI V12S
unstained BEAS-2B MR1 KO
no dox NaOH
no dox 6-FP
+ dox NaOH
+ dox 6-FP
MR1 - APC
HLA-Ia - PE
Normalized Counts
GFP
C
% GFP+ of Single Cells
% D4
% GFP
D
number of spots/well
% D4 max (prodrug)
no stim
Aux Mtb
M. smeg sup [5 µl]
5-A-RU prodrug [5 µM]
% GFP+ of Single Cells
% GFP
GFP+ GeoMFI
GFP
pCI D4
pCI D6ins
pCI D6del
pCI V12S
T cells only

**Figure S7: The D6del mutant does not contribute to the functional phenotype.** BEAS-2B MR1 KO cells were transiently transfected with pCI vectors encoding in-frame MR1 mutants identified in the clonal cell lines D4 or D6 (D6ins and D6del) or MR1 V12S with an IRES GFP. Transfected cells were split up and tested for total protein expression by WB (*A*) and MR1 surface expression by flow cytometry (*B*). Transfection efficiency was assessed via the GFP signal (*C*). Data from all three experiments (*A*+*C*) or one representative experiment (*B*) are shown. *D*. BEAS-2B MR1 KO cells were transfected as in A-C and used as APCs in IFNγ ELISPOTs with the indicated antigens. Data are pooled from three independent experiments, normalized to D4 at the highest antigen concentration and shown as mean with SD. Experimental groups were compared by repeated-measures ANOVA with Tukey's multiple comparisons test for each antigen separately (except for no stim controls) and statistically significant differences are indicated. * = $p \leq 0.05$ ** = $p \leq 0.01$ *** = $p \leq 0.001$ **** = $p \leq 0.0001$. Transfection efficiency and GFP expression levels for cells used in ELISPOTs was assessed by flow cytometry and compared by repeated-measures ANOVA as above. GeoMFI = geometric mean fluorescence intensity.

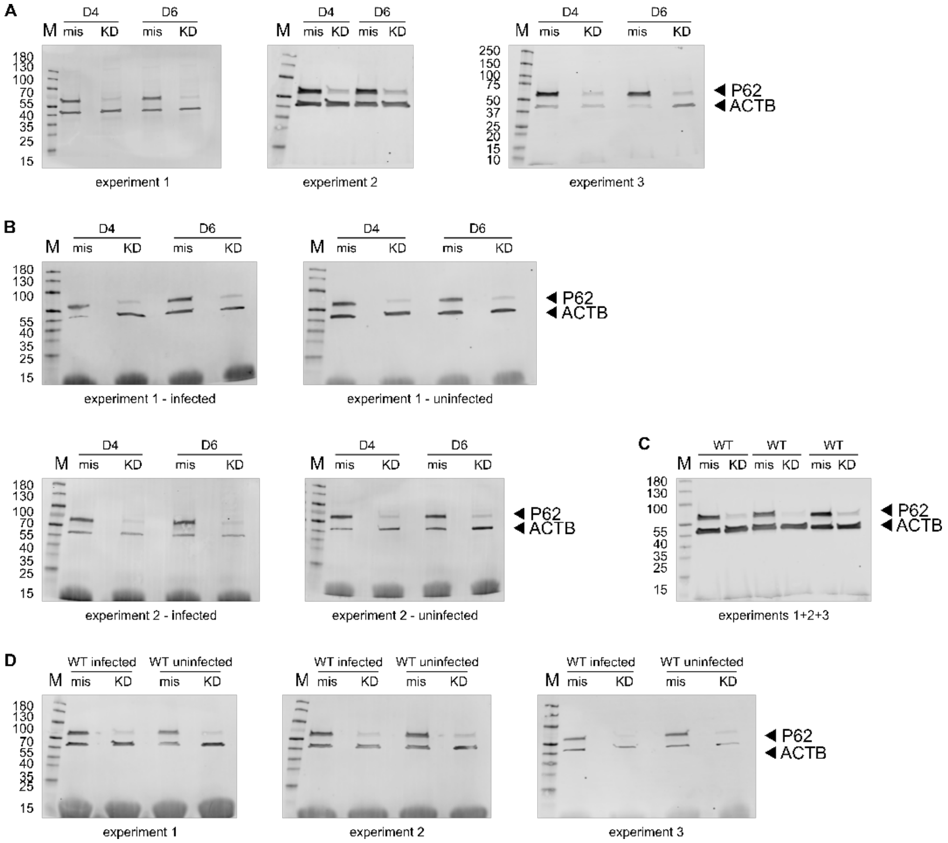

**Figure S8: SQSTM1/p62 expression levels in KD experiments.** At the time the cells were used in the ELISPOTs shown in Figure 5, some cells were lysed and analyzed by WB to assess SQSTM1/p62 KD on the protein level. Lysates from uninfected cells in *B+C* were from approximately 1.1e5 cells each, lysates from infected cells were from approximately 9e4 cells each. Beta actin (ACTB) was used as a loading control. Primary antibodies were from different species and detected in parallel with species-specific secondary antibodies conjugated to distinct IRDyes and both channels exported as greyscale. Molecular weight markers are indicated in kDa on the left. On occasion, lysates from two independent functional experiments were analyzed on the same blot as indicated.

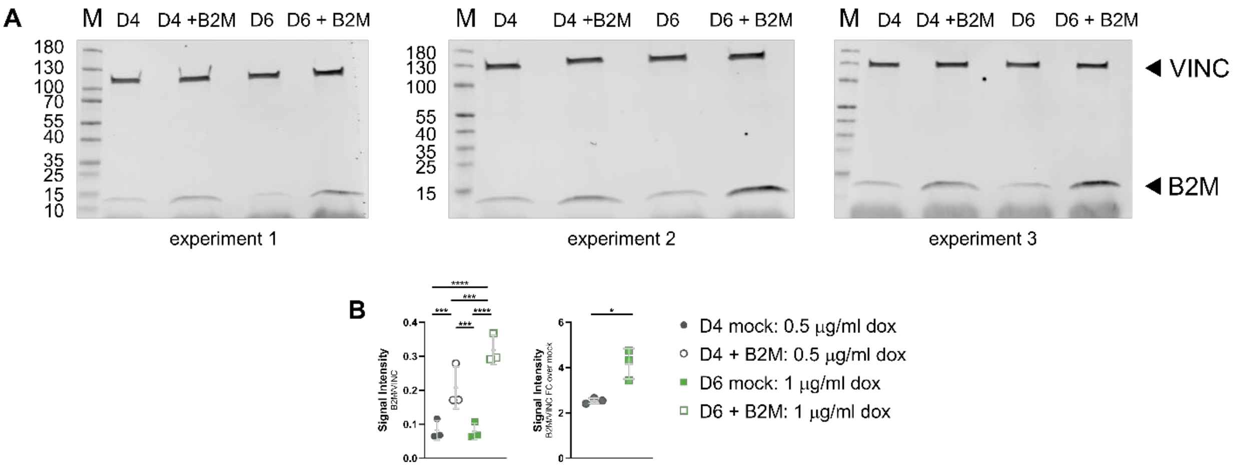

**Figure S9: B2M expression levels in OE experiments.** At the time the cells were used in the ELISPOTs shown in Figure 6A, some cells were lysed and analyzed by WB to assess B2M overexpression on the protein level. Vinculin (VINC) was used as a loading control. Primary antibodies were from different species and detected in parallel with species-specific secondary antibodies conjugated to distinct IRDyes and both channels exported as greyscale. Molecular weight markers are indicated in kDa on the left. Blots are shown in *A* and signal intensity relative to loading control (*left*) and mock-transfected cells (*right*) is shown in *B*. Western blots for experiments 2 and 3 were performed in parallel. Experimental groups were compared by repeated-measures ANOVA with Tukey's multiple comparisons test (*left*) or two-tailed, paired t-test (*right*) and statistically significant differences are indicated. * = p ≤ 0.05 ** = p ≤ 0.01 *** = p ≤ 0.001 **** = p ≤ 0.0001. FC = fold change.

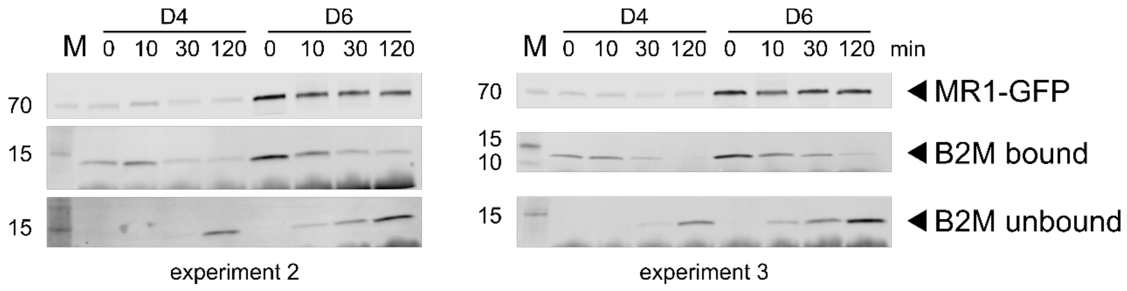

**Figure S10: Repeat experiments of the Western blots shown in Figure 6B.** WB analysis as in Figure 6B. MR1-GFP was immunoprecipitated from clonal cell lines D4 or D6 induced to express MR1-GFP with 0.5 or 8 µg/ml, respectively, overnight followed by incubation with the same concentrations of dox plus 100 µM 6-FP overnight. Beads were incubated at 37°C for the indicated time periods and MR1 bound to the beads, B2M bound to the beads, and B2M in the supernatant (unbound) were measured by WB. Primary antibodies were detected in parallel with species-specific secondary antibodies conjugated to IRDye800 and both channels exported as greyscale to visualize the molecular weight markers. The 800 channel was exported individually for MR1-GFP. Molecular weight markers are indicated in kDa on the left.

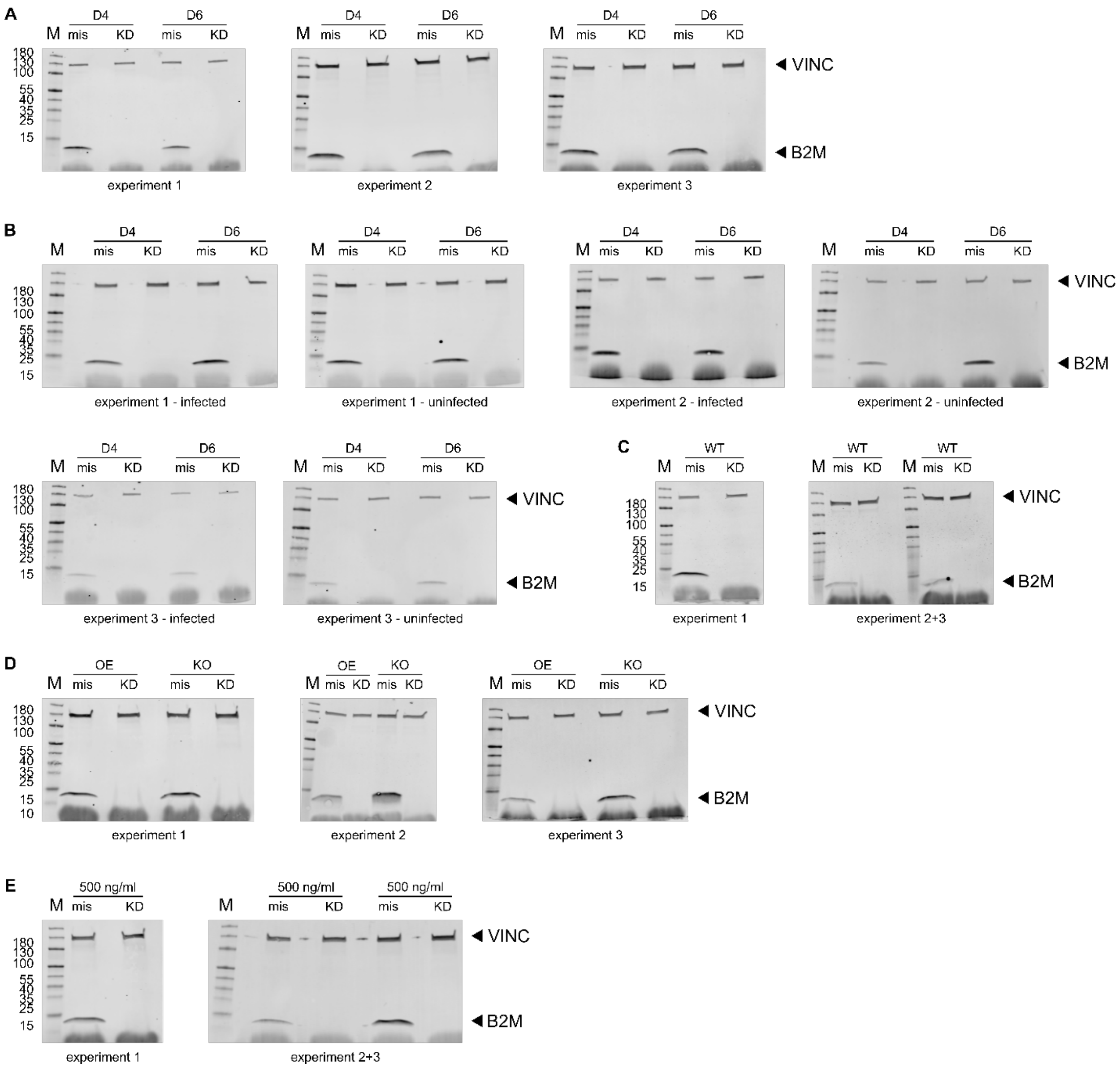

**Figure S11: B2M expression levels in KD experiments.** At the time the cells were used in the ELISPOTs shown in Figures 7 and 8, some cells were lysed and analyzed by WB to assess B2M KD on the protein level. Vinculin (VINC) was used as a loading control. Primary antibodies were from different species and detected in parallel with species-specific secondary antibodies conjugated to distinct IRDyes and both channels exported as greyscale. Molecular weight markers are indicated in kDa on the left. On occasion, lysates from two independent functional experiments were analyzed on the same blot as indicated.

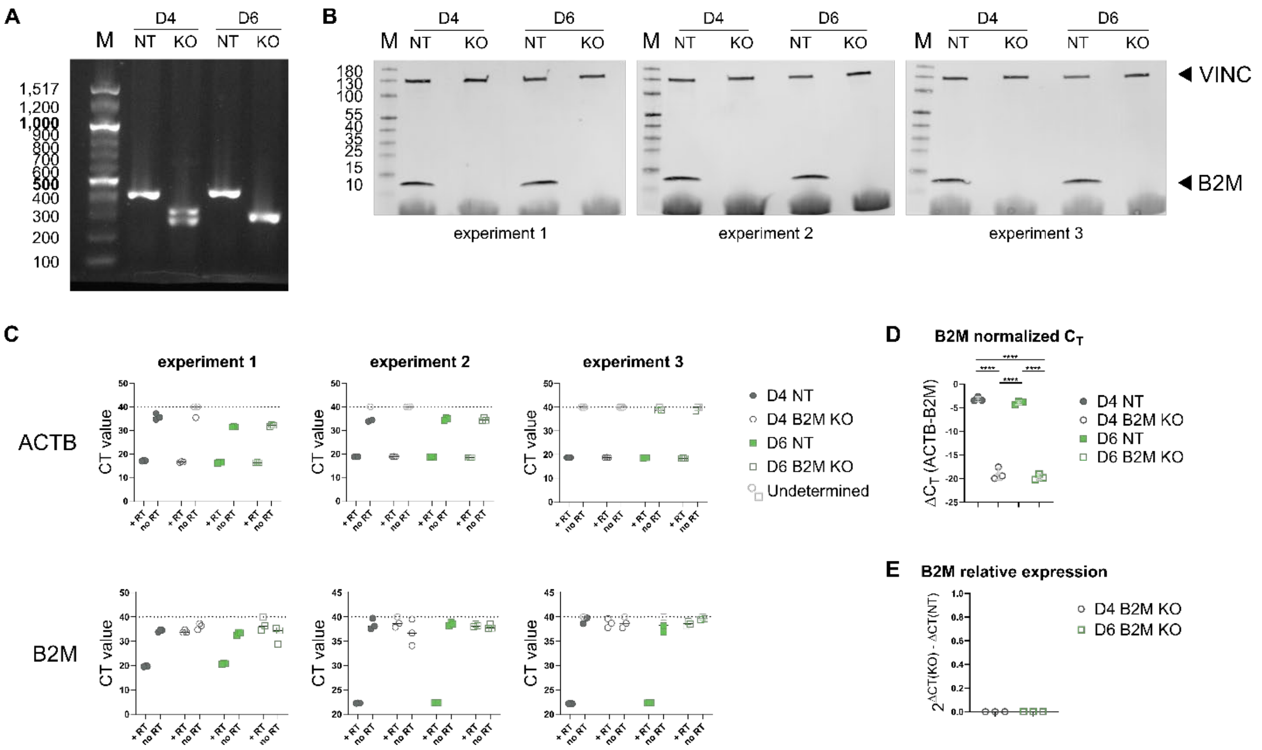

**Figure S12: Characterization of B2M CRISPR clones.** *A*. Genomic DNA was extracted from the clonal B2M KO cell lines (KO) and their respective controls treated with a non-targeted control RNP (NT) used in Figure 9, and the region targeted by the three sgRNAs was amplified by PCR. The amplicons were run on a 2% agarose gel and imaged with Gel Doc XR UV imaging system (Bio-Rad). Base pair (bp) sizes of the DNA ladder are indicated on the left. Expected band size in WT cells: 451 bp. *B*. B2M protein expression at baseline (no dox treatment) in the four clonal cell lines was analyzed by WB. Primary antibodies against B2M and loading control Vinculin (VINC) were from different species and detected in parallel with species-specific secondary antibodies conjugated to distinct IRDyes and both channels exported as greyscale. Molecular weight markers are indicated in kDa on the left. *C-E*. B2M transcript levels in the four clonal cell lines were measured by qRT-PCR. *C* shows the raw threshold cycle values ($C_T$) for each sample and its control without reverse transcriptase (noRT) for each experiment. Each dot represents a technical replicate (i.e. one well). “Undetermined” values were set to $C_T$=40 and colored in light grey for visualization. “Undetermined” values were excluded from the calculations of normalized $C_T$ values ($\Delta C_T$) in *D* and relative expression in *E*. Thus, each dot in *D* and *E* represents the mean of 1-3 technical replicates from one of the experiments shown in *C*. Experimental groups in *D* were compared by repeated-measures ANOVA with Tukey’s multiple comparisons test and statistically significant differences are indicated. * = $p \leq 0.05$ ** = $p \leq 0.01$ *** = $p \leq 0.001$ **** = $p \leq 0.0001$.

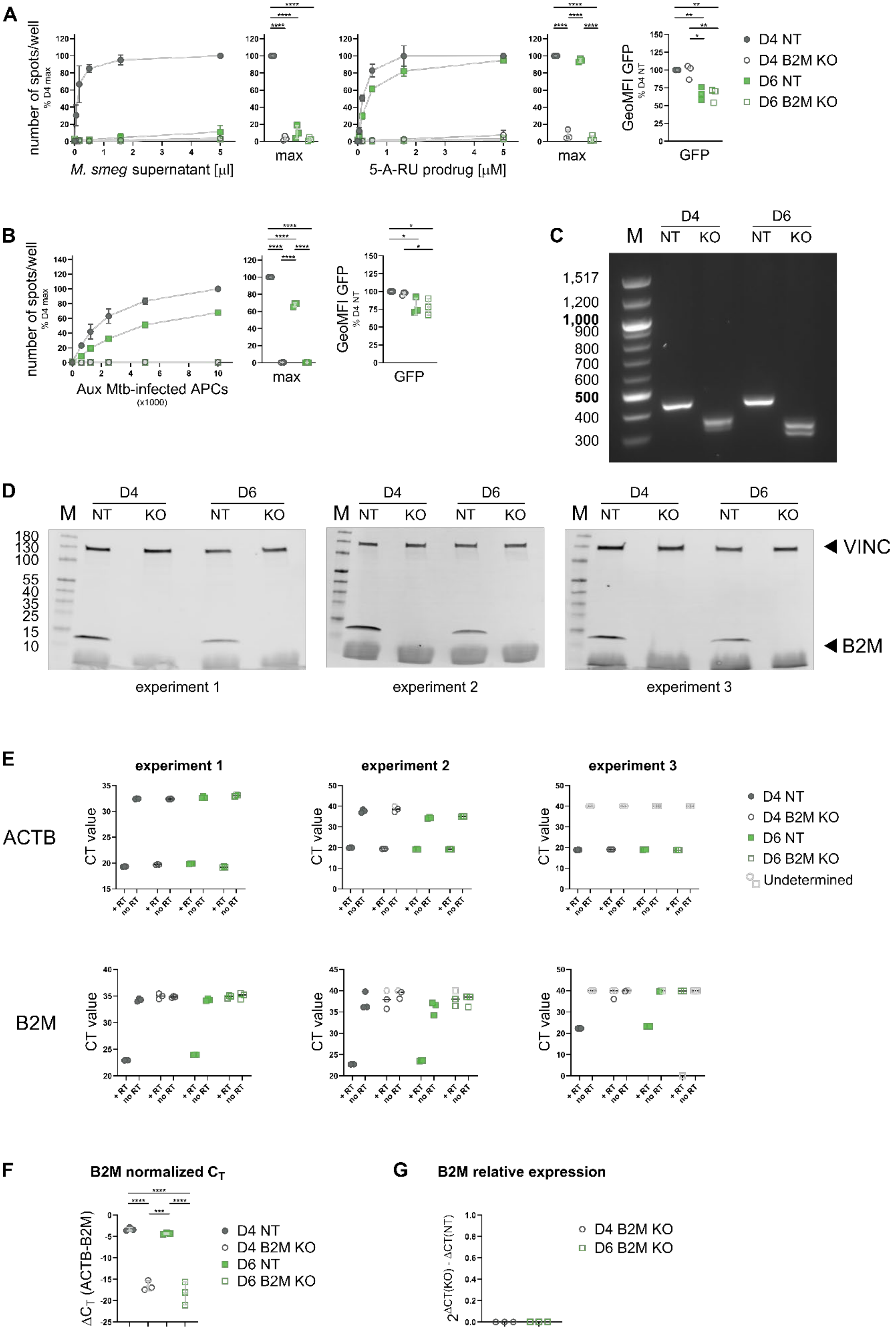
A
number of spots/well
% D4 max
M. smeg supernatant [µl]
max
5-A-RU prodrug [µM]
GeoMFI GFP
% D4 NT
GFP
D4 NT
D4 B2M KO
D6 NT
D6 B2M KO
B
Aux Mtb-infected APCs
(x1000)
C
M
D4
D6
NT
KO
1,517
1,200
1,000
900
800
700
600
500
400
300
D
180
130
100
55
40
35
25
15
10
VINC
B2M
experiment 1
experiment 2
experiment 3
E
ACTB
B2M
CT value
+ RT
no RT
Undetermined
F
B2M normalized $C_T$
$\Delta C_T$ (ACTB-B2M)
G
B2M relative expression
$2^{\Delta CT(KO) - \Delta CT(NT)}$

**Figure S13: Analysis of a second set of B2M KO clonal cell lines.** To confirm the B2M KO phenotype, a second set of B2M KO (KO) and non-target control (NT) clonal cell lines derived from D4 and D6 were analyzed as in Figures 9 and S12. *A*. Single clones were induced to express MR1-GFP with adjusted dox concentrations (see Figure S3) and used as APCs in IFNγ ELISPOTs with the indicated antigens. MR1-GFP expression was measured by flow cytometry at the time of the ELISPOT and shown to the right. Data are pooled from three independent experiments, normalized to D4 at the highest antigen concentration and shown as mean with SD. IFNγ responses at the highest antigen concentrations are additionally shown as dot plots. *C*. Genomic DNA was extracted and the region targeted by the three sgRNAs was amplified by PCR. The amplicons were run on a 2% agarose gel and imaged with Gel Doc XR UV imaging system (Bio-Rad). Base pair (bp) sizes of the DNA ladder are indicated on the left. Expected band size in WT cells: 451 bp. *D*. B2M protein expression at baseline (no dox treatment) in the four clonal cell lines was analyzed by WB. Primary antibodies against B2M and loading control Vinculin (VINC) were from different species and detected in parallel with species-specific secondary antibodies conjugated to distinct IRDyes and both channels exported as greyscale. Molecular weight markers are indicated in kDa on the left. *E-G*. B2M transcript levels in the four clonal cell lines were measured by qRT-PCR. *E* shows the raw threshold cycle values ($C_T$) for each sample and its control without reverse transcriptase (noRT) for each experiment. Each dot represents a technical replicate (i.e. one well). “Undetermined” values were set to $C_T$=40 and colored in light grey for visualization. “Undetermined” values were excluded from the calculations of normalized $C_T$ values ($\Delta C_T$) in *F* and relative expression in *G*. Thus, each dot in *F* and *G* represents the mean of 1-3 technical replicates from one of the experiments shown in *E*. Experimental groups in *A, B* + *F* were compared by repeated-measures ANOVA with Tukey’s multiple comparisons test and statistically significant differences are indicated. * = $p \leq 0.05$ ** = $p \leq 0.01$ *** = $p \leq 0.001$ **** = $p \leq 0.0001$. GeoMFI = geometric mean fluorescence intensity.

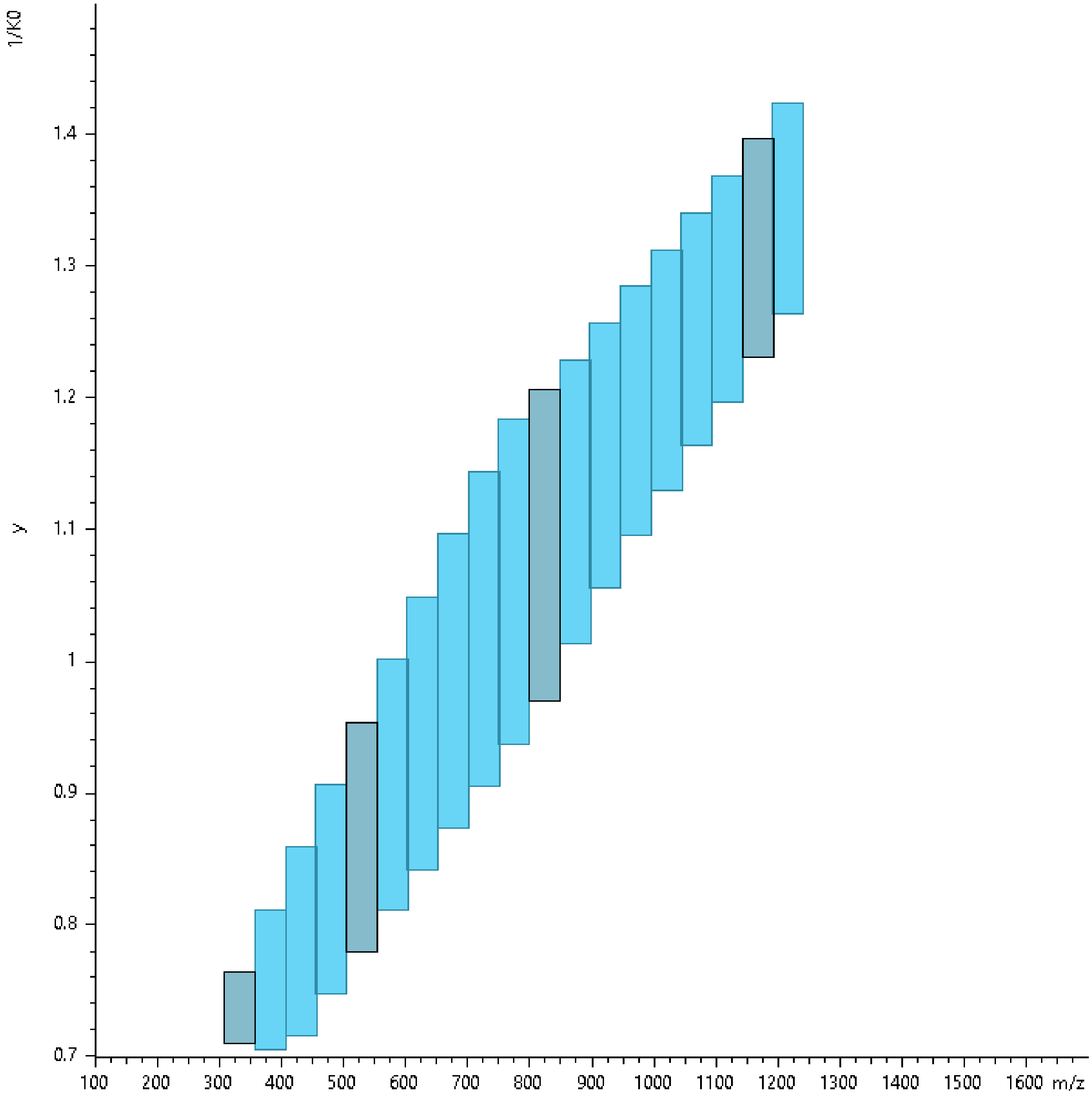

**Figure S14: dia-PASEF acquisition schemes on Bruker timsTOF Flex.** Scheme visualization was performed using the Bruker timsControl software.

**Table S1: Nonlinear fit Table of results for Fig 1C.** Table of Results for nonlinear regression analysis to determine whether data sets differ from each other. Straight line models were fit using least squares regression and compared using the extra sum-of-squares F test in Graphpad Prism 10.4.1.

| | | | | | |
|---|---|---|---|---|---|
| **Comparison of Fits** | | | | | |
| Null hypothesis | One curve for all data sets | | | | |
| Alternative hypothesis | Different curve for at least one data set | | | | |
| P value | 0.0126 | | | | |
| F (DFn, DFd) | 2.953 (6, 70) | | | | |
| **Different curve for at least one data set** | | | | | |
| Best-fit values | D4 | D6 | D8 | D16 | Global |
| YIntercept | 98.76 | 98.34 | 97.63 | 96.94 | |
| Slope | -2.941 | -2.848 | -2.44 | -2.692 | |
| 95% CI (profile likelihood) | | | | | |
| YIntercept | 92.00 to 105.5 | 92.60 to 104.1 | 92.80 to 102.5 | 91.94 to 101.9 | |
| Slope | -3.348 to -2.535 | -3.198 to -2.498 | -2.732 to -2.149 | -2.994 to -2.390 | |
| Goodness of Fit | | | | | |
| Degrees of Freedom | 17 | 17 | 18 | 18 | |
| R squared | 0.932 | 0.9454 | 0.9451 | 0.9512 | |
| Sum of Squares | 854.4 | 620 | 469.7 | 504.1 | |
| Sy.x | 7.089 | 6.039 | 5.108 | 5.292 | |
| **One curve for all data sets** | | | | | |
| Best-fit values | | | | | |
| YIntercept | 97.94 | 97.94 | 97.94 | 97.94 | 97.94 |
| Slope | -2.728 | -2.728 | -2.728 | -2.728 | -2.728 |
| 95% CI (profile likelihood) | | | | | |
| YIntercept | 95.08 to 100.8 | 95.08 to 100.8 | 95.08 to 100.8 | 95.08 to 100.8 | 95.08 to 100.8 |
| Slope | -2.900 to -2.555 | -2.900 to -2.555 | -2.900 to -2.555 | -2.900 to -2.555 | -2.900 to -2.555 |
| Goodness of Fit | | | | | |
| Degrees of Freedom | | | | | 76 |
| R squared | 0.9195 | 0.9409 | 0.8976 | 0.9506 | 0.929 |
| Sum of Squares | 1011 | 671 | 875.3 | 510.4 | 3068 |
| Sy.x | | | | | 6.354 |
| Constraints | | | | | |
| YIntercept | YIntercept is shared | YIntercept is shared | YIntercept is shared | YIntercept is shared | |
| Slope | Slope is shared | Slope is shared | Slope is shared | Slope is shared | |
| Number of points | | | | | |
| # of X values | 20 | 20 | 20 | 20 | |
| # Y values analyzed | 19 | 19 | 20 | 20 | |

**Table S2: ANOVA table for GFP GeoMFI with equal dox concentrations.** Relates to Fig 1B and Fig S4A. Table of Results for repeated-measures one-way ANOVA with Tukey's multiple comparisons test comparing GFP GeoMFI between experimental groups in Graphpad Prism 10.4.1.

| | | | | | |
|---|---|---|---|---|---|
| **Repeated measures ANOVA summary** | | | | | |
| Assume sphericity? | Yes | | | | |
| F | 138 | | | | |
| P value | <0.0001 | | | | |
| P value summary | **** | | | | |
| Statistically significant (P < 0.05)? | Yes | | | | |
| R squared | 0.9857 | | | | |
| Was the matching effective? | | | | | |
| F | 4.558 | | | | |
| P value | 0.0299 | | | | |
| P value summary | * | | | | |
| Is there significant matching (P < 0.05)? | Yes | | | | |
| R squared | 0.009219 | | | | |
| ANOVA table | SS | DF | MS | F (DFn, DFd) | P value |
| Treatment (between columns) | 2459701 | 7 | 351386 | F (7, 14) = 138.0 | P<0.0001 |
| Individual (between rows) | 23219 | 2 | 11610 | F (2, 14) = 4.558 | P=0.0299 |
| Residual (random) | 35660 | 14 | 2547 | | |
| Total | 2518579 | 23 | | | |
| Data summary | | | | | |
| Number of treatments (columns) | 8 | | | | |
| Number of subjects (rows) | 3 | | | | |
| Number of missing values | 0 | | | | |

| | | | | | |
|---|---|---|---|---|---|
| Number of families | | | 1 | | |
| Number of comparisons per family | | | 28 | | |
| Alpha | | | 0.05 | | |
| **Tukey's multiple comparisons test** | | | | | |
| | Mean Diff. | 95.00% CI of diff. | Below threshold? | Summary | Adjusted P Value |
| D4 NaOH vs. D6 NaOH | 306.3 | 160.9 to 451.7 | Yes | **** | <0.0001 |
| D4 NaOH vs. D8 NaOH | 580 | 434.6 to 725.4 | Yes | **** | <0.0001 |
| D4 NaOH vs. D16 NaOH | 663.3 | 517.9 to 808.7 | Yes | **** | <0.0001 |
| D4 NaOH vs. D4 6FP | -286.7 | -432.1 to -141.3 | Yes | *** | 0.0001 |
| D4 NaOH vs. D6 6FP | 321.7 | 176.3 to 467.1 | Yes | **** | <0.0001 |
| D4 NaOH vs. D8 6FP | 568.3 | 422.9 to 713.7 | Yes | **** | <0.0001 |
| D4 NaOH vs. D16 6FP | 651 | 505.6 to 796.4 | Yes | **** | <0.0001 |

| | | | | | | | | |
|---|---|---|---|---|---|---|---|---|
| D6 NaOH vs. D8 NaOH | 273.7 | 128.3 to 419.1 | Yes | *** | 0.0002 | | | |
| D6 NaOH vs. D16 NaOH | 357 | 211.6 to 502.4 | Yes | **** | <0.0001 | | | |
| D6 NaOH vs. D4 6FP | -593 | -738.4 to -447.6 | Yes | **** | <0.0001 | | | |
| D6 NaOH vs. D6 6FP | 15.33 | -130.1 to 160.7 | No | ns | >0.9999 | | | |
| D6 NaOH vs. D8 6FP | 262 | 116.6 to 407.4 | Yes | *** | 0.0004 | | | |
| D6 NaOH vs. D16 6FP | 344.7 | 199.3 to 490.1 | Yes | **** | <0.0001 | | | |
| D8 NaOH vs. D16 NaOH | 83.33 | -62.08 to 228.7 | No | ns | 0.5016 | | | |
| D8 NaOH vs. D4 6FP | -866.7 | -1012 to -721.3 | Yes | **** | <0.0001 | | | |
| D8 NaOH vs. D6 6FP | -258.3 | -403.7 to -112.9 | Yes | *** | 0.0004 | | | |
| D8 NaOH vs. D8 6FP | -11.67 | -157.1 to 133.7 | No | ns | >0.9999 | | | |
| D8 NaOH vs. D16 6FP | 71 | -74.41 to 216.4 | No | ns | 0.6745 | | | |
| D16 NaOH vs. D4 6FP | -950 | -1095 to -804.6 | Yes | **** | <0.0001 | | | |
| D16 NaOH vs. D6 6FP | -341.7 | -487.1 to -196.3 | Yes | **** | <0.0001 | | | |
| D16 NaOH vs. D8 6FP | -95 | -240.4 to 50.41 | No | ns | 0.3542 | | | |
| D16 NaOH vs. D16 6FP | -12.33 | -157.7 to 133.1 | No | ns | >0.9999 | | | |
| D4 6FP vs. D6 6FP | 608.3 | 462.9 to 753.7 | Yes | **** | <0.0001 | | | |
| D4 6FP vs. D8 6FP | 855 | 709.6 to 1000 | Yes | **** | <0.0001 | | | |
| D4 6FP vs. D16 6FP | 937.7 | 792.3 to 1083 | Yes | **** | <0.0001 | | | |
| D6 6FP vs. D8 6FP | 246.7 | 101.3 to 392.1 | Yes | *** | 0.0007 | | | |
| D6 6FP vs. D16 6FP | 329.3 | 183.9 to 474.7 | Yes | **** | <0.0001 | | | |
| D8 6FP vs. D16 6FP | 82.67 | -62.74 to 228.1 | No | ns | 0.5107 | | | |
| **Test details** | Mean 1 | Mean 2 | Mean Diff. | SE of diff. | n1 | n2 | q | DF |
| D4 NaOH vs. D6 NaOH | 1141 | 834.3 | 306.3 | 41.21 | 3 | 3 | 10.51 | 14 |
| D4 NaOH vs. D8 NaOH | 1141 | 560.7 | 580 | 41.21 | 3 | 3 | 19.91 | 14 |
| D4 NaOH vs. D16 NaOH | 1141 | 477.3 | 663.3 | 41.21 | 3 | 3 | 22.77 | 14 |
| D4 NaOH vs. D4 6FP | 1141 | 1427 | -286.7 | 41.21 | 3 | 3 | 9.838 | 14 |
| D4 NaOH vs. D6 6FP | 1141 | 819 | 321.7 | 41.21 | 3 | 3 | 11.04 | 14 |
| D4 NaOH vs. D8 6FP | 1141 | 572.3 | 568.3 | 41.21 | 3 | 3 | 19.5 | 14 |
| D4 NaOH vs. D16 6FP | 1141 | 489.7 | 651 | 41.21 | 3 | 3 | 22.34 | 14 |
| D6 NaOH vs. D8 NaOH | 834.3 | 560.7 | 273.7 | 41.21 | 3 | 3 | 9.392 | 14 |
| D6 NaOH vs. D16 NaOH | 834.3 | 477.3 | 357 | 41.21 | 3 | 3 | 12.25 | 14 |
| D6 NaOH vs. D4 6FP | 834.3 | 1427 | -593 | 41.21 | 3 | 3 | 20.35 | 14 |
| D6 NaOH vs. D6 6FP | 834.3 | 819 | 15.33 | 41.21 | 3 | 3 | 0.5262 | 14 |
| D6 NaOH vs. D8 6FP | 834.3 | 572.3 | 262 | 41.21 | 3 | 3 | 8.992 | 14 |
| D6 NaOH vs. D16 6FP | 834.3 | 489.7 | 344.7 | 41.21 | 3 | 3 | 11.83 | 14 |
| D8 NaOH vs. D16 NaOH | 560.7 | 477.3 | 83.33 | 41.21 | 3 | 3 | 2.86 | 14 |
| D8 NaOH vs. D4 6FP | 560.7 | 1427 | -866.7 | 41.21 | 3 | 3 | 29.74 | 14 |
| D8 NaOH vs. D6 6FP | 560.7 | 819 | -258.3 | 41.21 | 3 | 3 | 8.866 | 14 |
| D8 NaOH vs. D8 6FP | 560.7 | 572.3 | -11.67 | 41.21 | 3 | 3 | 0.4004 | 14 |
| D8 NaOH vs. D16 6FP | 560.7 | 489.7 | 71 | 41.21 | 3 | 3 | 2.437 | 14 |

| | | | | | | | | |
|---|---|---|---|---|---|---|---|---|
| D16 NaOH vs. D4 6FP | 477.3 | 1427 | -950 | 41.21 | 3 | 3 | 32.6 | 14 |
| D16 NaOH vs. D6 6FP | 477.3 | 819 | -341.7 | 41.21 | 3 | 3 | 11.73 | 14 |
| D16 NaOH vs. D8 6FP | 477.3 | 572.3 | -95 | 41.21 | 3 | 3 | 3.26 | 14 |
| D16 NaOH vs. D16 6FP | 477.3 | 489.7 | -12.33 | 41.21 | 3 | 3 | 0.4233 | 14 |
| D4 6FP vs. D6 6FP | 1427 | 819 | 608.3 | 41.21 | 3 | 3 | 20.88 | 14 |
| D4 6FP vs. D8 6FP | 1427 | 572.3 | 855 | 41.21 | 3 | 3 | 29.34 | 14 |
| D4 6FP vs. D16 6FP | 1427 | 489.7 | 937.7 | 41.21 | 3 | 3 | 32.18 | 14 |
| D6 6FP vs. D8 6FP | 819 | 572.3 | 246.7 | 41.21 | 3 | 3 | 8.465 | 14 |
| D6 6FP vs. D16 6FP | 819 | 489.7 | 329.3 | 41.21 | 3 | 3 | 11.3 | 14 |
| D8 6FP vs. D16 6FP | 572.3 | 489.7 | 82.67 | 41.21 | 3 | 3 | 2.837 | 14 |

**Table S3: ANOVA table for GFP GeoMFI with adjusted dox concentrations.** Relates to Fig S4. Table of Results for repeated-measures one-way ANOVA with Tukey's multiple comparisons test comparing GFP GeoMFI between experimental groups in Graphpad Prism 10.4.1.

| **Repeated measures ANOVA summary** | | | | | |
|---|---|---|---|---|---|
| Assume sphericity? | Yes | | | | |
| F | 7.531 | | | | |
| P value | 0.0007 | | | | |
| P value summary | *** | | | | |
| Statistically significant (P < 0.05)? | Yes | | | | |
| R squared | 0.7902 | | | | |
| Was the matching effective? | | | | | |
| F | 15.94 | | | | |
| P value | 0.0002 | | | | |
| P value summary | *** | | | | |
| Is there significant matching (P < 0.05)? | Yes | | | | |
| R squared | 0.3233 | | | | |
| ANOVA table | SS | DF | MS | F (DFn, DFd) | P value |
| Treatment (between columns) | 100910 | 7 | 14416 | F (7, 14) = 7.531 | P=0.0007 |
| Individual (between rows) | 61021 | 2 | 30511 | F (2, 14) = 15.94 | P=0.0002 |
| Residual (random) | 26798 | 14 | 1914 | | |
| Total | 188729 | 23 | | | |
| Data summary | | | | | |
| Number of treatments (columns) | 8 | | | | |
| Number of subjects (rows) | 3 | | | | |
| Number of missing values | 0 | | | | |

| | | | | | |
|---|---|---|---|---|---|
| Number of families | | | 1 | | |
| Number of comparisons per family | | | 28 | | |
| Alpha | | | 0.05 | | |
| **Tukey's multiple comparisons test** | | | | | |
| | Mean Diff. | 95.00% CI of diff. | Below threshold? | Summary | Adjusted P Value |
| D4 NaOH vs. D6 NaOH | 97.33 | -28.72 to 223.4 | No | ns | 0.1929 |
| D4 NaOH vs. D8 NaOH | 107.3 | -18.72 to 233.4 | No | ns | 0.123 |
| D4 NaOH vs. D16 NaOH | 106 | -20.05 to 232.1 | No | ns | 0.1308 |
| D4 NaOH vs. D4 6FP | -82.33 | -208.4 to 43.72 | No | ns | 0.3544 |
| D4 NaOH vs. D6 6FP | 89.67 | -36.39 to 215.7 | No | ns | 0.2663 |

| | | | | | | | | |
|---|---|---|---|---|---|---|---|---|
| D4 NaOH vs. D8 6FP | 103 | -23.05 to 229.1 | No | ns | 0.15 | | | |
| D4 NaOH vs. D16 6FP | 98.67 | -27.39 to 224.7 | No | ns | 0.1819 | | | |
| D6 NaOH vs. D8 NaOH | 10 | -116.1 to 136.1 | No | ns | >0.9999 | | | |
| D6 NaOH vs. D16 NaOH | 8.667 | -117.4 to 134.7 | No | ns | >0.9999 | | | |
| D6 NaOH vs. D4 6FP | -179.7 | -305.7 to -53.61 | Yes | ** | 0.0034 | | | |
| D6 NaOH vs. D6 6FP | -7.667 | -133.7 to 118.4 | No | ns | >0.9999 | | | |
| D6 NaOH vs. D8 6FP | 5.667 | -120.4 to 131.7 | No | ns | >0.9999 | | | |
| D6 NaOH vs. D16 6FP | 1.333 | -124.7 to 127.4 | No | ns | >0.9999 | | | |
| D8 NaOH vs. D16 NaOH | -1.333 | -127.4 to 124.7 | No | ns | >0.9999 | | | |
| D8 NaOH vs. D4 6FP | -189.7 | -315.7 to -63.61 | Yes | ** | 0.0021 | | | |
| D8 NaOH vs. D6 6FP | -17.67 | -143.7 to 108.4 | No | ns | 0.9995 | | | |
| D8 NaOH vs. D8 6FP | -4.333 | -130.4 to 121.7 | No | ns | >0.9999 | | | |
| D8 NaOH vs. D16 6FP | -8.667 | -134.7 to 117.4 | No | ns | >0.9999 | | | |
| D16 NaOH vs. D4 6FP | -188.3 | -314.4 to -62.28 | Yes | ** | 0.0022 | | | |
| D16 NaOH vs. D6 6FP | -16.33 | -142.4 to 109.7 | No | ns | 0.9997 | | | |
| D16 NaOH vs. D8 6FP | -3 | -129.1 to 123.1 | No | ns | >0.9999 | | | |
| D16 NaOH vs. D16 6FP | -7.333 | -133.4 to 118.7 | No | ns | >0.9999 | | | |
| D4 6FP vs. D6 6FP | 172 | 45.95 to 298.1 | Yes | ** | 0.005 | | | |
| D4 6FP vs. D8 6FP | 185.3 | 59.28 to 311.4 | Yes | ** | 0.0026 | | | |
| D4 6FP vs. D16 6FP | 181 | 54.95 to 307.1 | Yes | ** | 0.0032 | | | |
| D6 6FP vs. D8 6FP | 13.33 | -112.7 to 139.4 | No | ns | >0.9999 | | | |
| D6 6FP vs. D16 6FP | 9 | -117.1 to 135.1 | No | ns | >0.9999 | | | |
| D8 6FP vs. D16 6FP | -4.333 | -130.4 to 121.7 | No | ns | >0.9999 | | | |
| **Test details** | Mean 1 | Mean 2 | Mean Diff. | SE of diff. | n1 | n2 | q | DF |
| D4 NaOH vs. D6 NaOH | 466 | 368.7 | 97.33 | 35.72 | 3 | 3 | 3.853 | 14 |
| D4 NaOH vs. D8 NaOH | 466 | 358.7 | 107.3 | 35.72 | 3 | 3 | 4.249 | 14 |
| D4 NaOH vs. D16 NaOH | 466 | 360 | 106 | 35.72 | 3 | 3 | 4.196 | 14 |
| D4 NaOH vs. D4 6FP | 466 | 548.3 | -82.33 | 35.72 | 3 | 3 | 3.259 | 14 |
| D4 NaOH vs. D6 6FP | 466 | 376.3 | 89.67 | 35.72 | 3 | 3 | 3.55 | 14 |
| D4 NaOH vs. D8 6FP | 466 | 363 | 103 | 35.72 | 3 | 3 | 4.078 | 14 |
| D4 NaOH vs. D16 6FP | 466 | 367.3 | 98.67 | 35.72 | 3 | 3 | 3.906 | 14 |
| D6 NaOH vs. D8 NaOH | 368.7 | 358.7 | 10 | 35.72 | 3 | 3 | 0.3959 | 14 |
| D6 NaOH vs. D16 NaOH | 368.7 | 360 | 8.667 | 35.72 | 3 | 3 | 0.3431 | 14 |
| D6 NaOH vs. D4 6FP | 368.7 | 548.3 | -179.7 | 35.72 | 3 | 3 | 7.113 | 14 |
| D6 NaOH vs. D6 6FP | 368.7 | 376.3 | -7.667 | 35.72 | 3 | 3 | 0.3035 | 14 |
| D6 NaOH vs. D8 6FP | 368.7 | 363 | 5.667 | 35.72 | 3 | 3 | 0.2243 | 14 |
| D6 NaOH vs. D16 6FP | 368.7 | 367.3 | 1.333 | 35.72 | 3 | 3 | 0.05278 | 14 |
| D8 NaOH vs. D16 NaOH | 358.7 | 360 | -1.333 | 35.72 | 3 | 3 | 0.05278 | 14 |

| | | | | | | | | |
|---|---|---|---|---|---|---|---|---|
| D8 NaOH vs. D4 6FP | 358.7 | 548.3 | -189.7 | 35.72 | 3 | 3 | 7.509 | 14 |
| D8 NaOH vs. D6 6FP | 358.7 | 376.3 | -17.67 | 35.72 | 3 | 3 | 0.6994 | 14 |
| D8 NaOH vs. D8 6FP | 358.7 | 363 | -4.333 | 35.72 | 3 | 3 | 0.1716 | 14 |
| D8 NaOH vs. D16 6FP | 358.7 | 367.3 | -8.667 | 35.72 | 3 | 3 | 0.3431 | 14 |
| D16 NaOH vs. D4 6FP | 360 | 548.3 | -188.3 | 35.72 | 3 | 3 | 7.456 | 14 |
| D16 NaOH vs. D6 6FP | 360 | 376.3 | -16.33 | 35.72 | 3 | 3 | 0.6466 | 14 |
| D16 NaOH vs. D8 6FP | 360 | 363 | -3 | 35.72 | 3 | 3 | 0.1188 | 14 |
| D16 NaOH vs. D16 6FP | 360 | 367.3 | -7.333 | 35.72 | 3 | 3 | 0.2903 | 14 |
| D4 6FP vs. D6 6FP | 548.3 | 376.3 | 172 | 35.72 | 3 | 3 | 6.809 | 14 |
| D4 6FP vs. D8 6FP | 548.3 | 363 | 185.3 | 35.72 | 3 | 3 | 7.337 | 14 |
| D4 6FP vs. D16 6FP | 548.3 | 367.3 | 181 | 35.72 | 3 | 3 | 7.166 | 14 |
| D6 6FP vs. D8 6FP | 376.3 | 363 | 13.33 | 35.72 | 3 | 3 | 0.5278 | 14 |
| D6 6FP vs. D16 6FP | 376.3 | 367.3 | 9 | 35.72 | 3 | 3 | 0.3563 | 14 |
| D8 6FP vs. D16 6FP | 363 | 367.3 | -4.333 | 35.72 | 3 | 3 | 0.1716 | 14 |

**Table S4: List of protein identified by mass spectrometry after on-bead digest and DDA analysis.** See separate Excel file “SupportingInformationTable_S4”.

**Table S5: Lists of proteins and peptides identified by mass spectrometry after in-gel digest and DIA analysis.** See separate Excel file “SupportingInformationTable_S5”.

**Table S6: Nonlinear fit Table of results for Figure 6B.** Table of Results for nonlinear regression analysis to determine whether data sets differ from each other. One phase decay models were fit using least squares regression and compared using the extra sum-of-squares F test in Graphpad Prism 10.4.1.

| | | | |
|---|---|---|---|
| **Comparison of Fits** | | | |
| Null hypothesis | | | One curve for all data sets |
| Alternative hypothesis | | | Different curve for at least one data set |
| P value | | | 0.0007 |
| F (DFn, DFd) | | | 9.165 (3, 18) |
| **Different curve for at least one data set** | | | |
| Best-fit values | D4 | D6 | Global |
| Y0 | 104.1 | 101.2 | |
| Plateau | 15.59 | 13.65 | |
| K | 0.01665 | 0.04936 | |
| Half Life | 41.62 | 14.04 | |
| Tau | 60.05 | 20.26 | |
| Span | 88.55 | 87.56 | |
| 95% CI (profile likelihood) | | | |
| Y0 | 87.99 to 120.8 | 94.76 to 107.7 | |
| Plateau | ??? to 44.22 | 6.759 to 20.27 | |
| K | ??? to 0.04246 | 0.03868 to 0.06289 | |
| Half Life | 16.32 to ??? | 11.02 to 17.92 | |
| Tau | 23.55 to ??? | 15.90 to 25.85 | |
| Goodness of Fit | | | |
| Degrees of Freedom | 9 | 9 | |
| R squared | 0.8484 | 0.982 | |
| Sum of Squares | 1797 | 243.7 | |
| Sy.x | 14.13 | 5.203 | |
| Constraints | | | |
| K | K > 0 | K > 0 | |
| **One curve for all data sets** | | | |
| Best-fit values | | | |
| Y0 | 102.2 | 102.2 | 102.2 |
| Plateau | 18.94 | 18.94 | 18.94 |
| K | 0.03168 | 0.03168 | 0.03168 |
| Half Life | 21.88 | 21.88 | 21.88 |
| Tau | 31.57 | 31.57 | 31.57 |
| Span | 83.23 | 83.23 | 83.23 |
| 95% CI (profile likelihood) | | | |
| Y0 | 90.03 to 114.5 | 90.03 to 114.5 | 90.03 to 114.5 |

| | | | |
|---|---|---|---|
| Plateau | -1.844 to 33.44 | -1.844 to 33.44 | -1.844 to 33.44 |
| K | 0.01528 to 0.05376 | 0.01528 to 0.05376 | 0.01528 to 0.05376 |
| Half Life | 12.89 to 45.38 | 12.89 to 45.38 | 12.89 to 45.38 |
| Tau | 18.60 to 65.46 | 18.60 to 65.46 | 18.60 to 65.46 |
| Goodness of Fit | | | |
| Degrees of Freedom | | | 21 |
| R squared | 0.718 | 0.8659 | 0.8127 |
| Sum of Squares | 3344 | 1813 | 5157 |
| Sy.x | | | 15.67 |
| Constraints | | | |
| Y0 | Y0 is shared | Y0 is shared | |
| Plateau | Plateau is shared | Plateau is shared | |
| K | K > 0 and shared | K > 0 and shared | |
| Number of points | | | |
| # of X values | 12 | 12 | |
| # Y values analyzed | 12 | 12 | |

**Table S7: Nonlinear fit Table of results for surface MR1 in Figure 6C.** Table of Results for nonlinear regression analysis to determine whether data sets differ from each other. One phase decay models were fit using least squares regression with K constrained to > 0 and compared using the extra sum-of-squares F test in Graphpad Prism 10.4.1.

| | | | |
|---|---|---|---|
| **Comparison of Fits** | | | |
| Null hypothesis | | | One curve for all data sets |
| Alternative hypothesis | | | Different curve for at least one data set |
| P value | | | <0.0001 |
| F (DFn, DFd) | | | 66.77 (3, 24) |
| **Different curve for at least one data set** | | | |
| Best-fit values | D4 | D6 | Global |
| Y0 | 100.6 | 99.95 | |
| Plateau | 3.366E-09 | 5.894 | |
| K | 0.1167 | 0.3215 | |
| Half Life | 5.938 | 2.156 | |
| Tau | 8.566 | 3.111 | |
| Span | 100.6 | 94.06 | |
| 95% CI (profile likelihood) | | | |
| Y0 | 96.09 to 105.1 | 94.62 to 105.3 | |
| Plateau | ??? to 36.33 | ??? to 17.79 | |
| K | 0.09888 to 0.2331 | 0.3151 to 0.4419 | |
| Half Life | 2.974 to 7.010 | 1.569 to 2.200 | |
| Tau | 4.290 to 10.11 | 2.263 to 3.174 | |
| Goodness of Fit | | | |
| Degrees of Freedom | 12 | 12 | |
| R squared | 0.9514 | 0.9813 | |
| Sum of Squares | 255 | 238.6 | |
| Sy.x | 4.61 | 4.459 | |
| Constraints | | | |
| Plateau | Plateau > 0 | Plateau > 0 | |
| K | K > 0 | K > 0 | |
| **One curve for all data sets** | | | |
| Best-fit values | | | |
| Y0 | 99.54 | 99.54 | 99.54 |
| Plateau | 8.295 | 8.295 | 8.295 |
| K | 0.2104 | 0.2104 | 0.2104 |
| Half Life | 3.294 | 3.294 | 3.294 |
| Tau | 4.752 | 4.752 | 4.752 |
| Span | 91.24 | 91.24 | 91.24 |
| 95% CI (profile likelihood) | | | |

| | | | |
|---|---|---|---|
| Y0 | 90.15 to 110.0 | 90.15 to 110.0 | 90.15 to 110.0 |
| Plateau | -infinity to 39.63 | -infinity to 39.63 | -infinity to 39.63 |
| K | 0.1392 to 0.5012 | 0.1392 to 0.5012 | 0.1392 to 0.5012 |
| Half Life | 1.383 to 4.979 | 1.383 to 4.979 | 1.383 to 4.979 |
| Tau | 1.995 to 7.183 | 1.995 to 7.183 | 1.995 to 7.183 |
| Goodness of Fit | | | |
| Degrees of Freedom | | | 27 |
| R squared | 0.5609 | 0.8189 | 0.7813 |
| Sum of Squares | 2306 | 2308 | 4614 |
| Sy.x | | | 13.07 |
| Constraints | | | |
| Y0 | Y0 is shared | Y0 is shared | |
| Plateau | Plateau > 0 and shared | Plateau > 0 and shared | |
| K | K > 0 and shared | K > 0 and shared | |
| Number of points | | | |
| # of X values | 15 | 15 | |
| # Y values analyzed | 15 | 15 | |

**Table S8: Nonlinear fit Table of results for total MR1 (GFP) in Figure 6C.** Table of Results for nonlinear regression analysis to determine whether data sets differ from each other. One phase decay models were fit using least squares regression with K constrained to > 0 and compared using the extra sum-of-squares F test in Graphpad Prism 10.4.1.

| | | | |
|---|---|---|---|
| **Comparison of Fits** | | | |
| Null hypothesis | | | One curve for all data sets |
| Alternative hypothesis | | | Different curve for at least one data set |
| P value | | | <0.0001 |
| F (DFn, DFd) | | | 61.94 (3, 24) |
| **Different curve for at least one data set** | | | |
| Best-fit values | | | |
| Y0 | 102.1 | 100.6 | |
| Plateau | 3.958E-09 | 21 | |
| K | 0.08881 | 0.2096 | |
| Half Life | 7.805 | 3.307 | |
| Tau | 11.26 | 4.771 | |
| Span | 102.1 | 79.57 | |
| 95% CI (profile likelihood) | | | |
| Y0 | 99.65 to 104.6 | 97.89 to 103.3 | |
| Plateau | ??? to 30.42 | ??? to 33.37 | |
| K | 0.07975 to 0.1441 | 0.1390 to 0.2876 | |
| Half Life | 4.811 to 8.692 | 2.410 to 4.986 | |
| Tau | 6.941 to 12.54 | 3.477 to 7.193 | |
| Goodness of Fit | | | |
| Degrees of Freedom | 12 | 12 | |
| R squared | 0.977 | 0.9898 | |
| Sum of Squares | 81.81 | 64.28 | |
| Sy.x | 2.611 | 2.314 | |
| Constraints | | | |
| Plateau | Plateau > 0 | Plateau > 0 | |
| K | K > 0 | K > 0 | |
| **One curve for all data sets** | | | |
| Best-fit values | | | |
| Y0 | 100.7 | 100.7 | 100.7 |
| Plateau | 3.877 | 3.877 | 3.877 |
| K | 0.1196 | 0.1196 | 0.1196 |
| Half Life | 5.795 | 5.795 | 5.795 |
| Tau | 8.36 | 8.36 | 8.36 |
| Span | 96.81 | 96.81 | 96.81 |
| 95% CI (profile likelihood) | | | |

| | | | |
|---|---|---|---|
| Y0 | 96.14 to 106.0 | 96.14 to 106.0 | 96.14 to 106.0 |
| Plateau | -infinity to 45.12 | -infinity to 45.12 | -infinity to 45.12 |
| K | 0.09566 to 0.2942 | 0.09566 to 0.2942 | 0.09566 to 0.2942 |
| Half Life | 2.356 to 7.246 | 2.356 to 7.246 | 2.356 to 7.246 |
| Tau | 3.399 to 10.45 | 3.399 to 10.45 | 3.399 to 10.45 |
| Goodness of Fit | | | |
| Degrees of Freedom | | | 27 |
| R squared | 0.8204 | 0.8986 | 0.881 |
| Sum of Squares | 637.4 | 639.9 | 1277 |
| Sy.x | | | 6.878 |
| Constraints | | | |
| Y0 | Y0 is shared | Y0 is shared | |
| Plateau | Plateau > 0 and shared | Plateau > 0 and shared | |
| K | K > 0 and shared | K > 0 and shared | |
| Number of points | | | |
| # of X values | 15 | 15 | |
| # Y values analyzed | 15 | 15 | |

**Table S9: ANOVA table for GFP GeoMFI with dox titration.** Relates to Figure 8C. Table of Results for repeated-measures one-way ANOVA with Tukey's multiple comparisons test comparing GFP GeoMFI between experimental groups in Graphpad Prism 10.4.1.

| **Repeated measures ANOVA summary** | | | | | |
|---|---|---|---|---|---|
| Assume sphericity? | Yes | | | | |
| F | 124.6 | | | | |
| P value | <0.0001 | | | | |
| P value summary | **** | | | | |
| Statistically significant (P < 0.05)? | Yes | | | | |
| R squared | 0.9842 | | | | |
| Was the matching effective? | | | | | |
| F | 39.87 | | | | |
| P value | <0.0001 | | | | |
| P value summary | **** | | | | |
| Is there significant matching (P < 0.05)? | Yes | | | | |
| R squared | 0.06543 | | | | |
| ANOVA table | SS | DF | MS | F (DFn, DFd) | P value |
| Treatment (between columns) | 18377 | 9 | 2042 | F (9, 18) = 124.6 | P<0.0001 |
| Individual (between rows) | 1307 | 2 | 653.6 | F (2, 18) = 39.87 | P<0.0001 |
| Residual (random) | 295.1 | 18 | 16.39 | | |
| Total | 19979 | 29 | | | |
| Data summary | | | | | |
| Number of treatments (columns) | 10 | | | | |
| Number of subjects (rows) | 3 | | | | |
| Number of missing values | 0 | | | | |

| | | | | | |
|---|---|---|---|---|---|
| Number of families | | | 1 | | |
| Number of comparisons per family | | | 45 | | |
| Alpha | | | 0.05 | | |
| **Tukey's multiple comparisons test** | | | | | |
| | Mean Diff. | 95.00% CI of diff. | Below threshold? | Summary | Adjusted P Value |
| mis 0 vs. mis 62.5 | -3.646 | -15.50 to 8.208 | No | ns | 0.978 |
| mis 0 vs. mis 125 | -11.32 | -23.17 to 0.5335 | No | ns | 0.0682 |
| mis 0 vs. mis 250 | -35.49 | -47.34 to -23.64 | Yes | **** | <0.0001 |
| mis 0 vs. mis 500 | -77.56 | -89.41 to -65.71 | Yes | **** | <0.0001 |
| mis 0 vs. KD 0 | -1.02 | -12.87 to 10.83 | No | ns | >0.9999 |

| | | | | | |
|---|---|---|---|---|---|
| mis 0 vs. KD 62.5 | -2.803 | -14.66 to 9.050 | No | ns | 0.9964 |
| mis 0 vs. KD 125 | -8.919 | -20.77 to 2.935 | No | ns | 0.2449 |
| mis 0 vs. KD 250 | -23.05 | -34.90 to -11.19 | Yes | **** | <0.0001 |
| mis 0 vs. KD 500 | -52.7 | -64.55 to -40.84 | Yes | **** | <0.0001 |
| mis 62.5 vs. mis 125 | -7.674 | -19.53 to 4.179 | No | ns | 0.4214 |
| mis 62.5 vs. mis 250 | -31.84 | -43.70 to -19.99 | Yes | **** | <0.0001 |
| mis 62.5 vs. mis 500 | -73.91 | -85.77 to -62.06 | Yes | **** | <0.0001 |
| mis 62.5 vs. KD 0 | 2.626 | -9.227 to 14.48 | No | ns | 0.9978 |
| mis 62.5 vs. KD 62.5 | 0.8427 | -11.01 to 12.70 | No | ns | >0.9999 |
| mis 62.5 vs. KD 125 | -5.273 | -17.13 to 6.580 | No | ns | 0.8344 |
| mis 62.5 vs. KD 250 | -19.4 | -31.25 to -7.548 | Yes | *** | 0.0005 |
| mis 62.5 vs. KD 500 | -49.05 | -60.90 to -37.20 | Yes | **** | <0.0001 |
| mis 125 vs. mis 250 | -24.17 | -36.02 to -12.32 | Yes | **** | <0.0001 |
| mis 125 vs. mis 500 | -66.24 | -78.09 to -54.39 | Yes | **** | <0.0001 |
| mis 125 vs. KD 0 | 10.3 | -1.553 to 22.15 | No | ns | 0.1208 |
| mis 125 vs. KD 62.5 | 8.517 | -3.336 to 20.37 | No | ns | 0.2951 |
| mis 125 vs. KD 125 | 2.401 | -9.452 to 14.25 | No | ns | 0.9989 |
| mis 125 vs. KD 250 | -11.73 | -23.58 to 0.1257 | No | ns | 0.0538 |
| mis 125 vs. KD 500 | -41.38 | -53.23 to -29.52 | Yes | **** | <0.0001 |
| mis 250 vs. mis 500 | -42.07 | -53.92 to -30.22 | Yes | **** | <0.0001 |
| mis 250 vs. KD 0 | 34.47 | 22.62 to 46.32 | Yes | **** | <0.0001 |
| mis 250 vs. KD 62.5 | 32.69 | 20.83 to 44.54 | Yes | **** | <0.0001 |
| mis 250 vs. KD 125 | 26.57 | 14.72 to 38.42 | Yes | **** | <0.0001 |
| mis 250 vs. KD 250 | 12.44 | 0.5891 to 24.30 | Yes | * | 0.0352 |
| mis 250 vs. KD 500 | -17.21 | -29.06 to -5.355 | Yes | ** | 0.0018 |
| mis 500 vs. KD 0 | 76.54 | 64.69 to 88.39 | Yes | **** | <0.0001 |
| mis 500 vs. KD 62.5 | 74.76 | 62.90 to 86.61 | Yes | **** | <0.0001 |
| mis 500 vs. KD 125 | 68.64 | 56.79 to 80.49 | Yes | **** | <0.0001 |
| mis 500 vs. KD 250 | 54.51 | 42.66 to 66.36 | Yes | **** | <0.0001 |
| mis 500 vs. KD 500 | 24.86 | 13.01 to 36.71 | Yes | **** | <0.0001 |
| KD 0 vs. KD 62.5 | -1.783 | -13.64 to 10.07 | No | ns | 0.9999 |
| KD 0 vs. KD 125 | -7.899 | -19.75 to 3.954 | No | ns | 0.3851 |
| KD 0 vs. KD 250 | -22.03 | -33.88 to -10.17 | Yes | *** | 0.0001 |
| KD 0 vs. KD 500 | -51.68 | -63.53 to -39.82 | Yes | **** | <0.0001 |
| KD 62.5 vs. KD 125 | -6.116 | -17.97 to 5.737 | No | ns | 0.6989 |
| KD 62.5 vs. KD 250 | -20.24 | -32.10 to -8.391 | Yes | *** | 0.0003 |
| KD 62.5 vs. KD 500 | -49.89 | -61.75 to -38.04 | Yes | **** | <0.0001 |
| KD 125 vs. KD 250 | -14.13 | -25.98 to -2.275 | Yes | * | 0.0126 |
| KD 125 vs. KD 500 | -43.78 | -55.63 to -31.93 | Yes | **** | <0.0001 |

| | | | | | | | | |
|---|---|---|---|---|---|---|---|---|
| KD 250 vs. KD 500 | -29.65 | -41.50 to -17.80 | Yes | **** | <0.0001 | | | |
| Test details | Mean 1 | Mean 2 | Mean Diff. | SE of diff. | n1 | n2 | q | DF |
| mis 0 vs. mis 62.5 | 22.44 | 26.09 | -3.646 | 3.306 | 3 | 3 | 1.56 | 18 |
| mis 0 vs. mis 125 | 22.44 | 33.76 | -11.32 | 3.306 | 3 | 3 | 4.842 | 18 |
| mis 0 vs. mis 250 | 22.44 | 57.93 | -35.49 | 3.306 | 3 | 3 | 15.18 | 18 |
| mis 0 vs. mis 500 | 22.44 | 100 | -77.56 | 3.306 | 3 | 3 | 33.18 | 18 |
| mis 0 vs. KD 0 | 22.44 | 23.46 | -1.02 | 3.306 | 3 | 3 | 0.4362 | 18 |
| mis 0 vs. KD 62.5 | 22.44 | 25.24 | -2.803 | 3.306 | 3 | 3 | 1.199 | 18 |
| mis 0 vs. KD 125 | 22.44 | 31.36 | -8.919 | 3.306 | 3 | 3 | 3.815 | 18 |
| mis 0 vs. KD 250 | 22.44 | 45.49 | -23.05 | 3.306 | 3 | 3 | 9.859 | 18 |
| mis 0 vs. KD 500 | 22.44 | 75.14 | -52.7 | 3.306 | 3 | 3 | 22.54 | 18 |
| mis 62.5 vs. mis 125 | 26.09 | 33.76 | -7.674 | 3.306 | 3 | 3 | 3.283 | 18 |
| mis 62.5 vs. mis 250 | 26.09 | 57.93 | -31.84 | 3.306 | 3 | 3 | 13.62 | 18 |
| mis 62.5 vs. mis 500 | 26.09 | 100 | -73.91 | 3.306 | 3 | 3 | 31.62 | 18 |
| mis 62.5 vs. KD 0 | 26.09 | 23.46 | 2.626 | 3.306 | 3 | 3 | 1.123 | 18 |
| mis 62.5 vs. KD 62.5 | 26.09 | 25.24 | 0.8427 | 3.306 | 3 | 3 | 0.3605 | 18 |
| mis 62.5 vs. KD 125 | 26.09 | 31.36 | -5.273 | 3.306 | 3 | 3 | 2.256 | 18 |
| mis 62.5 vs. KD 250 | 26.09 | 45.49 | -19.4 | 3.306 | 3 | 3 | 8.299 | 18 |
| mis 62.5 vs. KD 500 | 26.09 | 75.14 | -49.05 | 3.306 | 3 | 3 | 20.98 | 18 |
| mis 125 vs. mis 250 | 33.76 | 57.93 | -24.17 | 3.306 | 3 | 3 | 10.34 | 18 |
| mis 125 vs. mis 500 | 33.76 | 100 | -66.24 | 3.306 | 3 | 3 | 28.34 | 18 |
| mis 125 vs. KD 0 | 33.76 | 23.46 | 10.3 | 3.306 | 3 | 3 | 4.406 | 18 |
| mis 125 vs. KD 62.5 | 33.76 | 25.24 | 8.517 | 3.306 | 3 | 3 | 3.643 | 18 |
| mis 125 vs. KD 125 | 33.76 | 31.36 | 2.401 | 3.306 | 3 | 3 | 1.027 | 18 |
| mis 125 vs. KD 250 | 33.76 | 45.49 | -11.73 | 3.306 | 3 | 3 | 5.017 | 18 |
| mis 125 vs. KD 500 | 33.76 | 75.14 | -41.38 | 3.306 | 3 | 3 | 17.7 | 18 |
| mis 250 vs. mis 500 | 57.93 | 100 | -42.07 | 3.306 | 3 | 3 | 18 | 18 |
| mis 250 vs. KD 0 | 57.93 | 23.46 | 34.47 | 3.306 | 3 | 3 | 14.75 | 18 |
| mis 250 vs. KD 62.5 | 57.93 | 25.24 | 32.69 | 3.306 | 3 | 3 | 13.98 | 18 |
| mis 250 vs. KD 125 | 57.93 | 31.36 | 26.57 | 3.306 | 3 | 3 | 11.37 | 18 |
| mis 250 vs. KD 250 | 57.93 | 45.49 | 12.44 | 3.306 | 3 | 3 | 5.323 | 18 |
| mis 250 vs. KD 500 | 57.93 | 75.14 | -17.21 | 3.306 | 3 | 3 | 7.361 | 18 |
| mis 500 vs. KD 0 | 100 | 23.46 | 76.54 | 3.306 | 3 | 3 | 32.74 | 18 |
| mis 500 vs. KD 62.5 | 100 | 25.24 | 74.76 | 3.306 | 3 | 3 | 31.98 | 18 |
| mis 500 vs. KD 125 | 100 | 31.36 | 68.64 | 3.306 | 3 | 3 | 29.36 | 18 |
| mis 500 vs. KD 250 | 100 | 45.49 | 54.51 | 3.306 | 3 | 3 | 23.32 | 18 |
| mis 500 vs. KD 500 | 100 | 75.14 | 24.86 | 3.306 | 3 | 3 | 10.63 | 18 |
| KD 0 vs. KD 62.5 | 23.46 | 25.24 | -1.783 | 3.306 | 3 | 3 | 0.7628 | 18 |

| | | | | | | | | |
|---|---|---|---|---|---|---|---|---|
| KD 0 vs. KD 125 | 23.46 | 31.36 | -7.899 | 3.306 | 3 | 3 | 3.379 | 18 |
| KD 0 vs. KD 250 | 23.46 | 45.49 | -22.03 | 3.306 | 3 | 3 | 9.423 | 18 |
| KD 0 vs. KD 500 | 23.46 | 75.14 | -51.68 | 3.306 | 3 | 3 | 22.11 | 18 |
| KD 62.5 vs. KD 125 | 25.24 | 31.36 | -6.116 | 3.306 | 3 | 3 | 2.616 | 18 |
| KD 62.5 vs. KD 250 | 25.24 | 45.49 | -20.24 | 3.306 | 3 | 3 | 8.66 | 18 |
| KD 62.5 vs. KD 500 | 25.24 | 75.14 | -49.89 | 3.306 | 3 | 3 | 21.34 | 18 |
| KD 125 vs. KD 250 | 31.36 | 45.49 | -14.13 | 3.306 | 3 | 3 | 6.044 | 18 |
| KD 125 vs. KD 500 | 31.36 | 75.14 | -43.78 | 3.306 | 3 | 3 | 18.73 | 18 |
| KD 250 vs. KD 500 | 45.49 | 75.14 | -29.65 | 3.306 | 3 | 3 | 12.68 | 18 |

**Table S10: Proteins identified by mass spectrometry after in-gel digest and DIA analysis that showed a statistically significant batch effect.**

| ProteinID | Gene | ConditionWT | BatchDay2 | BatchDay3 | AveExpr | F | P.Value | adj.P.Val |
|---|---|---|---|---|---|---|---|---|
| Q9UQ35 | SRRM2 | 0.28528 | -1.05295 | -6.06925 | 16.35781 | 1654.77 | 3.61E-22 | 5.33E-19 |
| Q92572 | AP3S1 | 0.076903 | 0.522364 | -3.97102 | 14.57861 | 1395.993 | 1.66E-21 | 1.22E-18 |
| Q13144 | EIF2B5 | 0.064008 | 0.943977 | -5.69049 | 13.01635 | 1026.646 | 2.60E-20 | 1.28E-17 |
| Q8NC51 | SERBP1 | 0.022965 | -0.29102 | -2.72809 | 14.91425 | 963.8496 | 4.58E-20 | 1.69E-17 |
| P52272 | HNRNPM | -0.16751 | -0.52085 | -2.55946 | 15.11057 | 931.0728 | 6.24E-20 | 1.84E-17 |
| P26038 | MSN | -0.0166 | -1.33717 | -2.95219 | 13.89116 | 907.8099 | 7.82E-20 | 1.92E-17 |
| A0A0A0MRM9 | NOLC1 | 0.008183 | -0.68494 | -5.87713 | 15.03803 | 838.0488 | 1.60E-19 | 3.37E-17 |
| Q07065 | CKAP4 | 0.081128 | -0.26723 | -2.37724 | 14.6895 | 674.0332 | 1.12E-18 | 2.06E-16 |
| O14974 | PPP1R12A | -0.15354 | 0.241904 | -2.54269 | 14.0712 | 655.8396 | 1.43E-18 | 2.34E-16 |
| Q86UE4 | MTDH | -0.15537 | -0.43513 | -3.09421 | 14.36369 | 642.3895 | 1.72E-18 | 2.53E-16 |

### Supporting Information References

1. Clement, K., et al., *CRISPResso2 provides accurate and rapid genome editing sequence analysis.* Nat Biotechnol, 2019. **37**(3): p. 224-226.
